# Boosting reversible photocontrol of a photoxenase by an engineered conformational shift

**DOI:** 10.1101/2025.08.25.672088

**Authors:** Sabrina Mandl, Janet Sánchez, Miquel Estévez, Astrid Bruckmann, Caroline Hiefinger, Sílvia Osuna, Andrea Hupfeld

## Abstract

Our study successfully explores strategies to effectively improve the photocontrol efficiency of light-sensitive enzymes, dubbed photoxenases, with photoswitchable unnatural amino acids (UAAs). The engineering of photoxenases is a versatile method for the reversible photocontrol in various applications. To boost the photocontrol of an established allosteric and heterodimeric photoxenase based on imidazole glycerol phosphate synthase, we turned from an ineffective tuning of the UAA photochemistry to a semi-rational enzyme design. Remarkably, mutations at the catalytically important heterodimer interface increased the light-regulation factor (LRF) for the *k*_*cat*_ up to ~100 with near-quantitative reversibility. Steady-state kinetic investigations combined with computationally determined correlation-based Shortest-Path-Map analysis and conformational landscapes revealed how photocontrol was altered in the two best hits. The LRF(*k*_*cat*_) correlated with a shift of a conformational equilibrium between an active and inactive population at the targeted active site and a tuned population productivity upon irradiation. While the overall reduced *k*_*cat*_ values originated from a rewiring of the allosteric signal transmission, the increased LRF(*k*_*cat*_) resulted from a change in i) the size of the conformational shift, ii) the population productivity, and iii) the conformational heterogeneity. With this, our findings provide initial guidelines to boost photocontrol and underscore the power of photoxenase engineering.

## Introduction

Life is based on many actions and reactions, which requires a high adaptability of natural processes. Enzymes are at the bottom of most of these processes and comply with this requirement through their high flexibility allowing different mechanisms, such as allostery,^[1,2]^ to exactly control their activity. Within the last decades, enzymes have gained a strong foothold in various biotechnological processes, e.g., as biosensors,^[3,4]^ drugs^[5]^ and green catalysts.^[6–8]^ Similar to nature, these applications strive for the precise and reversible control of enzymes. Besides de novo design of allostery,^[9–11]^ light can be used to manipulate enzyme activity with high spatiotemporal resolution that is cost-effective, environmentally friendly and easy to apply.

In recent years, engineering of enzymes with photoswitchable unnatural amino acids (UAAs) has emerged as a versatile method for the reversible regulation with light.^[12–18]^ We named the resulting enzyme family photoxenases (“photo”: light; “xeno”: foreign to nature; “-ase”: common suffix for enzymes). Light-sensitivity is thereby genetically encoded via amber suppression and the use of engineered, orthogonal aminoacyl-tRNA-synthetase (aaRS)/tRNA pairs.^[19]^ The so far most commonly used photoswitchable UAA, termed AzoF, contains an azobenzene extension to the natural phenylalanine.^[20,21]^ AzoF exists in two sterically different isomers, a thermodynamically favored *E* and a metastable [*t*_*½*_(25°C) ~16 d]^[22]^ *Z* isomer. Since the absorbance spectra of *E* and *Z* overlap, irradiation induces the formation of an isomeric equilibrium called photostationary state (PSS) that depends on the wavelength of the applied light. As a result, light of 365 nm shifts the *E*-enriched thermal equilibrium (TEQ) towards a *Z*-enriched PSS^365^ (**Figure 1A**), whereas light of 420 nm obtains an *E*-enriched PSS^420^. Even though synthetic photoswitches are well-established tools for the reversible photocontrol of biological processes,^[23–25]^ mechanistic insights on how photoswitchable UAAs affect enzyme activity are sparse, which creates a major bottleneck towards effective strategies for photoxenase engineering. Previous engineering endeavors have primarily generated photoxenases with light-regulation factors (LRFs) of ~2.^[12,14,18]^ Only recently, we could show that the incorporation of AzoF close to flexible domains that undergo catalytically relevant conformational changes allows for LRFs up to ~10.^[26–28]^ In comparison, allosteric (de)activation factors of natural enzymes range between 2- and >4000 to adapt to the respective environmental requirements.^[29–32]^ Likewise, we expect that each biotechnological application has its individual demands on the photocontrol efficiency of photoxenases; for some doubling of yields may suffice, while others might benefit from LRFs of >100. Hence, to meet the individual needs, a strategy is required that customizes and particularly increases the photocontrol efficiency of photoxenases.

**Figure 1.**
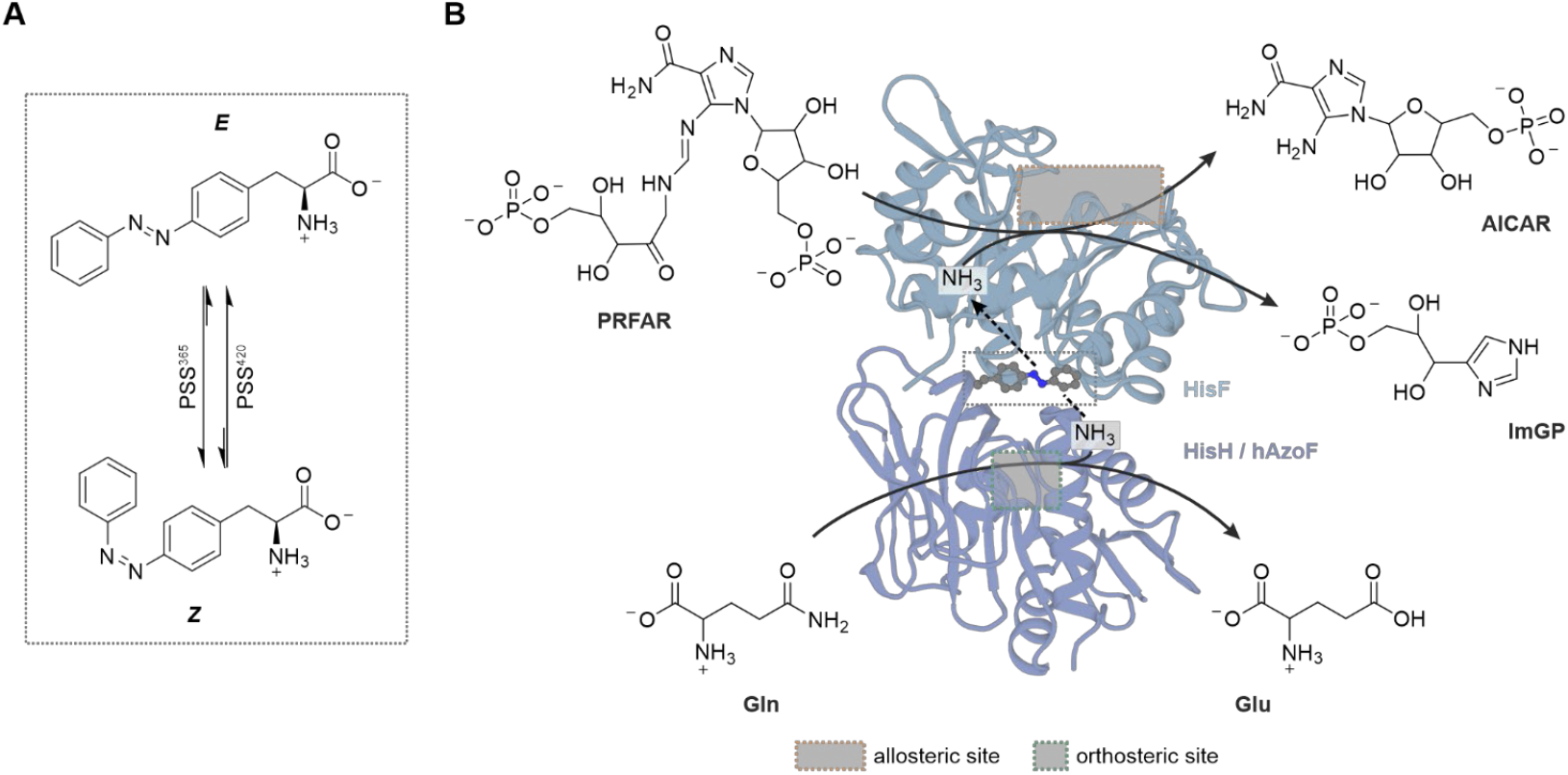
Photoxenase engineering model hAzoF_HisF. A) Light-induced reactions of AzoF. B) The heterodimer HisH_HisF couples glutamine (Gln) hydrolysis at the HisH active site with the production of imidazole glycerol phosphate (ImGP) and 5-aminoimidazol-4-carboxamidribotide (AICAR) at the HisF active site by ammonia-induced turnover of PRFAR. Incorporation of AzoF at position W123 generates hAzoF_HisF and facilitates effective photocontrol of glutamine hydrolysis. Glu: glutamate. PDB: 6YMU.

There are two potential ways to boost photocontrol in photoxenases. First, photoconversion yields of the UAA can be improved towards higher *Z* fractions in PSS^365^ and higher *E* fractions in PSS^420^ either by optimizing the wavelength of irradiation or using a different photoswitch. A second protein-based approach towards higher photocontrol efficiencies is presented by photoreceptor engineering in optogenetics.^[33]^ Here, the initially low LRFs of identified photoreceptor variants are further increased by directed evolution such as semi-rational design strategies.^[34]^

In this work, we set out to test both approaches, photochemical tuning and semi-rational design, on a photoxenase. To this end, we chose imidazole glycerol phosphate synthase (HisH_HisF), for which initial photoxenases are available,^[26,27]^ as a model system. HisH_HisF from *Thermotoga maritima* is a well-characterized allosteric bi-enzyme complex (**Figure 1B**). Its glutaminase subunit HisH hydrolyzes glutamine to generate ammonia via a catalytic E180-H178-C84 triad.^[35]^ Ammonia travels through an intermolecular channel to the active site of the second subunit HisF, where it reacts with N′-[(5′-phosphoribulosyl)formimino]-5-aminoimidazole-4-carboxamide ribonucleotide (PRFAR). HisH shows only residual activity as a monomer and is poorly activated by complexation with HisF. However, binding of PRFAR or its precursor N′-[(5′-phosphoribosyl)formimino]-5-aminoimidazole-4-carboxamide ribonucleotide (ProFAR) to the active site in HisF, termed allosteric site, stimulates glutamine hydrolysis at the active site in HisH, termed orthosteric site.^[36,37]^ Computational studies have postulated an allosteric network that connects the more than 25 Å apart active sites of HisH and HisF.^[38–44]^ This leads to a backbone flip at the HisH active site, which fully forms a catalytically relevant oxyanion hole.^[38,42–45]^ Notably, the highly flexible loop1 at the HisF active site appears to be part of this allosteric signaling network.^[30,38,40,46]^ Moreover, a hinge motion using the interaction between R249 in HisF and W123 in HisH at the subunit interface was confirmed to be involved in the allosteric activation.^[38,47]^ Remarkably, replacement of W123 with AzoF in one of the HisH_HisF-based photoxenases (here named hAzoF_HisF) established photocontrol of glutamine hydrolysis with an LRF of ~12 with however limited reversibility.^[27]^

To pave the way for highly effective photoxenase engineering strategies, we aimed to investigate i) whether we can improve the photocontrol efficiency of hAzoF_HisF, ii) by which method, and iii) how this improvement is achieved in mechanistic detail. While tuning of the photochemistry at position W123 was largely ineffective, further engineering of the enzyme scaffold in a semi-rational and combinatorial design resulted in significantly improved photocontrol efficiencies. Furthermore, combination of extensive steady-state kinetic evaluations and computational studies using Molecular Dynamics (MD) simulations coupled to correlation-based Shortest-Path-Map (SPM) analysis allowed us to illuminate the mechanism of photocontrol in the parental construct and two best hits.

## Results and Discussion

### Photochemical tuning of the photoswitchable UAA

Although azobenzenes are frequently used photoswitches, they are limited by non-quantitative photoconversion yields, particularly in the *E*→*Z* switch. Moreover, our recent studies on isolated AzoF (PSS^365^: 14*E*:86*Z*)^[22]^ and HisH containing AzoF at position W123 (PSS^365^: 27*E*:73*Z*)^[27]^ demonstrated that the *E*:*Z* ratio varies after incorporation. To optimize the ratio towards a higher Z content, we initially tried different irradiation wavelengths for the *E*→*Z* switch. For this, we excluded wavelengths <365 nm because of the high UV light sensitivity of ProFAR (*λ*_*max*_ = 270 nm), which we used as allosteric ligand throughout this study as it previously yielded higher LRFs.^[26,27]^ However, irradiation with >365 nm reduced the *Z* content even further (**Figure S1**). Interestingly, during the course of this work, the arylazopyrazole-based UAA AapF (**Figure 2A**) was introduced, which showed higher photoconversion yields (PSS^365^: 9*E*:91*Z*) than AzoF and which can be co-translationally incorporated in *Escherichia coli* via orthogonal aaRS/tRNA pairs.^[48,49]^ Hence, we compared recombinantly produced HisH containing either AzoF (hAzoF) or AapF (hAapF) at position W123. Initial characterizations revealed that hAzoF and hAapF were obtained with high purity (>95%), contained AzoF or AapF at position W123 and were properly folded (**Figure S2–S4A**). Interestingly, hAapF in complex with HisF (hAapF_HisF) appeared to be more stable during thermal denaturation (*T*_*m*_ = 106°C) than hAzoF_HisF (*T*_*m*_ = 96°C; **Figure S4B**) and comparably stable to wild-type HisH_HisF (*T*_*m*_ = 109°C^[27]^). Remarkably, UV/Vis spectra (**Figure 2B**) confirmed improved photoconversion yields for hAapF in both the *E*→*Z* and the *Z*→*E* switch (365 nm: 13*E*:87*Z*; 540 nm: 92*E*:8*Z*) compared to the previously determined values for hAzoF^[27]^ (365 nm: 27*E*:73*Z*; 420 nm: 83*E*:17*Z*). We then compared the rates of glutamine hydrolysis for hAzoF_HisF and hAapF_HisF in their TEQ, PSS^365^ and PSS^420/540^ (**Figure 2C**), for which we used a previously established colorimetric coupled enzymatic assay (**Figure S5**).^[26,27]^ Similar to our recent study, hAzoF_HisF showed an 11-fold activation in PSS^365^ with 24% reversibility in PSS^420^. In contrast, glutaminase activity of hAapF_HisF was unaffected by irradiation despite the better photoconversion of AapF. These results imply that the photocontrol efficiency is not only defined by the photoswitch but also by the enzyme itself and particularly suggest a decisive role of interactions between the photoswitchable UAA and residues of the enzyme in its direct proximity. This is in line with two current studies, in which we demonstrated that various photoswitchable UAAs^[22]^ as well as an alternative AzoF isomer induced upon dichromatic irradiation^[50]^ mediate different photocontrol efficiencies within the same positions in HisH_HisF. While we argue that the versatility of different photoswitches is beneficial for the initial identification of photoxenases, it appears to be largely ineffective to further boost the photocontrol efficiency of an already established photoxenase.

**Figure 2.**
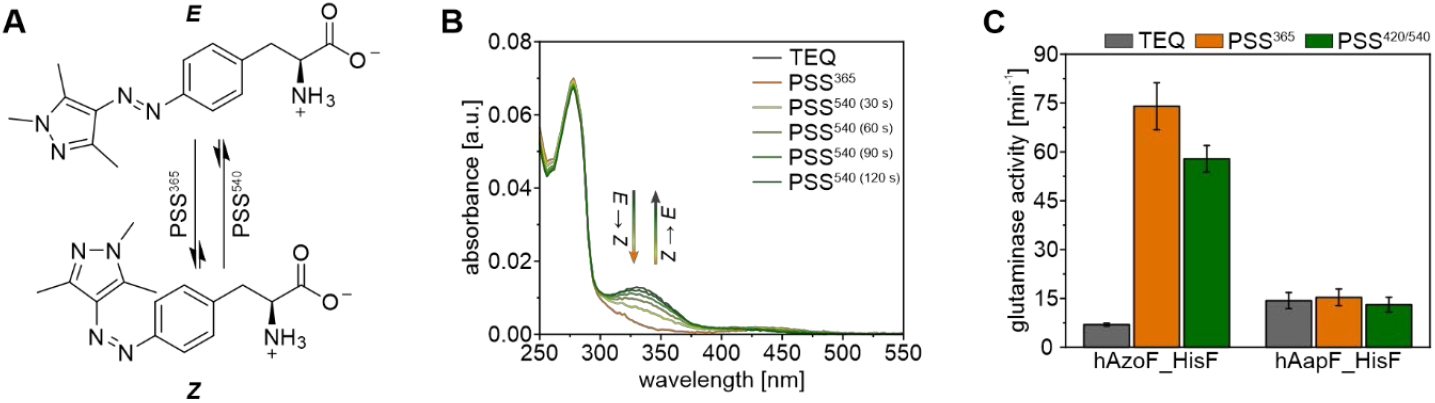
Photochemical tuning in position W123 of HisH. A) Light-induced reactions of AapF. B) UV/Vis spectroscopy shows good (365 nm) and near-quantitative (540 nm) photoconversion yields for hAapF. C) Replacement of AzoF with AapF results in complete loss of photocontrol in hAapF_HisF. Glutaminase activity relates to the rate of glutamine hydrolysis normalized to the enzyme concentration (*v/E*_*0*_). Bars and error bars represent the mean ± standard deviation (SD) of technical triplicates. *Irradiation:* 2 s per well with 365 nm for PSS^365^ and 4 s per well with 420 nm for PSS^420/540^ (individual setup, **Table S8**).

## Semi-rational and combinatorial design of hAzoF_HisF variants

In parallel, we set out to improve the photocontrol efficiency of hAzoF_HisF by protein engineering. For this, we chose a semi-rational design approach (**Figure 3A**). Crystal structure analysis in our previous study showed that AzoF in position 123 of hAzoF maintained the essential cation-π interaction with R249 in HisF and established an additional hydrophobic interaction with I75 in HisF.^[27]^ Moreover, our findings indicated that photocontrol is associated with a conformational change of the subunit interface, which is important for the allosteric activation of glutamine hydrolysis. Based on these results, we decided to randomize residues that are i) in direct proximity to AzoF or ii) located at the interface and relevant for the allosteric activation of HisH (**Figure 3B**). To this end, we first selected all residues within a 4 Å distance to the AzoF molecule from the crystal structure. For the second approach, we initially reduced the number of positions at the interface by searching for residues that changed their sidechain conformation in a structural comparison of the inactive and active HisH_HisF conformation.^[45,46]^ We then performed an alanine scan of the resulting five positions to determine which of these are allosterically connected to the orthosteric site (**Figure S6**). While three of the tested variants (hN12A_HisF, hR117A_HisF and HisH_fQ123A) showed only minor effects on glutamine hydrolysis, hR18A_HisF and HisH_fI73A exhibited >5-fold decreased glutaminase rates. Hence, we created gene libraries targeting the 16 positions close to AzoF and the two positions identified at the more distant interface. For this, we performed scanning mutagenesis obtaining 19 library members for each residue position.

**Figure 3.**
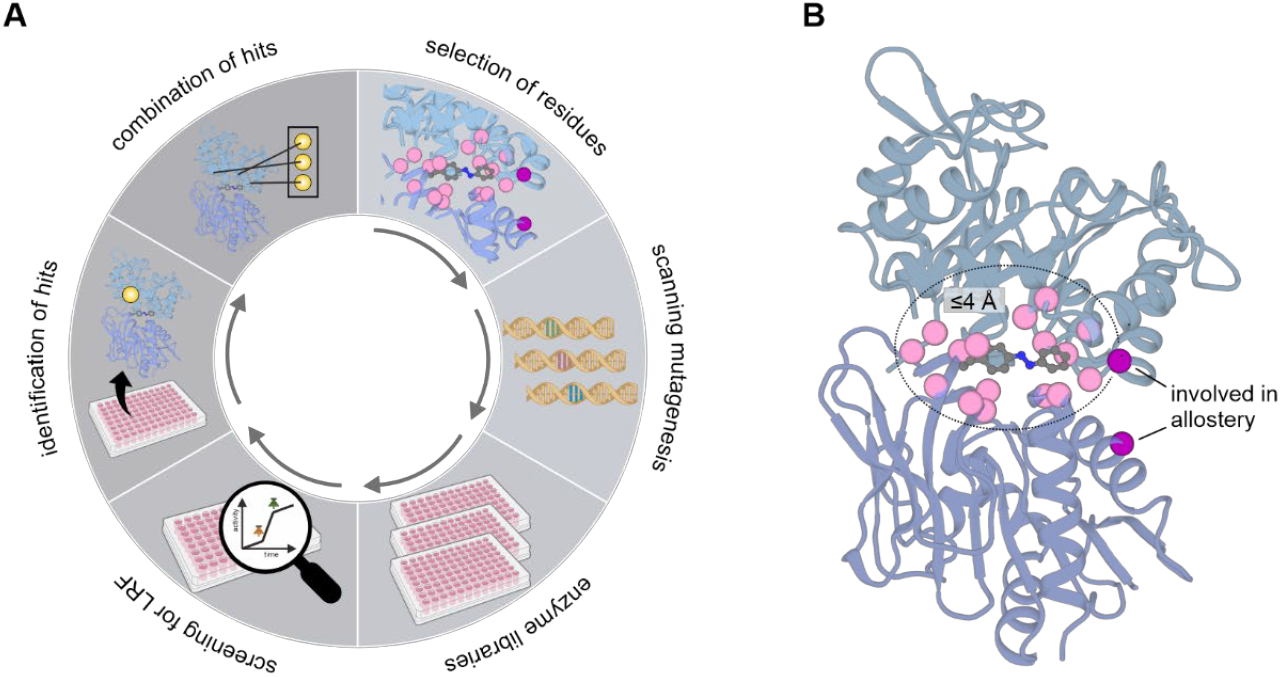
Semi-rational design approach for the improvement of photocontrol in hAzoF_HisF. A) Individual steps in the engineering process. B) Selected residue positions surrounding AzoF (pink spheres) or at the more distant interface (magenta spheres) for randomization and production of enzyme libraries. AzoF: grey. PDB: 6YMU.

We then established a screening assay for the evaluation of photocontrol efficiencies using 96-well microtiter plates (**Extended Text S1**; **Figures S7–S10**). Remarkably, we thereby found that the standard *E*.*coli* expression host BL21 Gold (DE3), which allows for high expression yields required for directed evolution experiments, outperformed the B95.ΔAΔfabR^[51]^ strain, which was previously engineered for UAA incorporation, since it showed only negligible misincorporation of natural amino acids. A standardized split-GFP assay^[52–54]^ thereby allowed us to track the enzyme content in the lysates. Furthermore, we chose our real-time photocontrol assay for the activity-based screening, in which we generally start the glutaminase reaction by the addition of purified enzyme (or lysate) in its TEQ, spectrophotometrically follow its progress using our colorimetric assay, and then determine the photocontrol effect upon 365 nm and subsequent 420 nm irradiation in real-time. To improve the signal-to-noise ratio in this assay, we removed most *E. coli* proteins from the cell lysates via a heat step. Quality control by determining Z′-factors confirmed the reliability of this optimized screening approach.

Remarkably, screening of all 342 hAzoF_HisF variants that were produced from the 18 gene libraries identified three variants, hAzoF_fL2D, hAzoF_fI75K and hAzoF_fI73P, with improved LRFs (1.3-fold, 1.5-fold, and 1.8-fold) and slightly enhanced reversibility (revs.) of photocontrol compared to hAzoF_HisF (**Figure 4A**; **Table S1**). While positions fL2 and fI75 are located in direct proximity to AzoF, fI73 is an allosterically relevant residue position. We produced and purified fL2D, fI73P and fI75K in large scale with purities >95% (**Figure S2**) for further evaluation of the three identified hits in comparison to the parental construct. For this, we recorded real-time (rt) photocontrol measurements of glutamine hydrolysis with glutamine in saturation and high auxiliary enzyme concentrations (**Figure 4B**). The latter reduced the lag phase of the coupled enzymatic assay, which improved its response time after the addition of enzyme as well as after irradiation. While we achieved an LRF(rt) of 15 and a revs.(rt) of ~42% for the parental construct, each hit showed a significantly increased LRF(rt) (46–98) and revs.(rt) of photocontrol (55–66%) (**Figure 4C–D**; **Table S2**). Notably, the improved photocontrol efficiencies mainly originated from a reduced activity in TEQ. Moreover, all hAzoF_HisF variants showed either similar or higher activities in PSS^365^ compared to the light-insensitive HisH_HisF as control.

**Figure 4.**
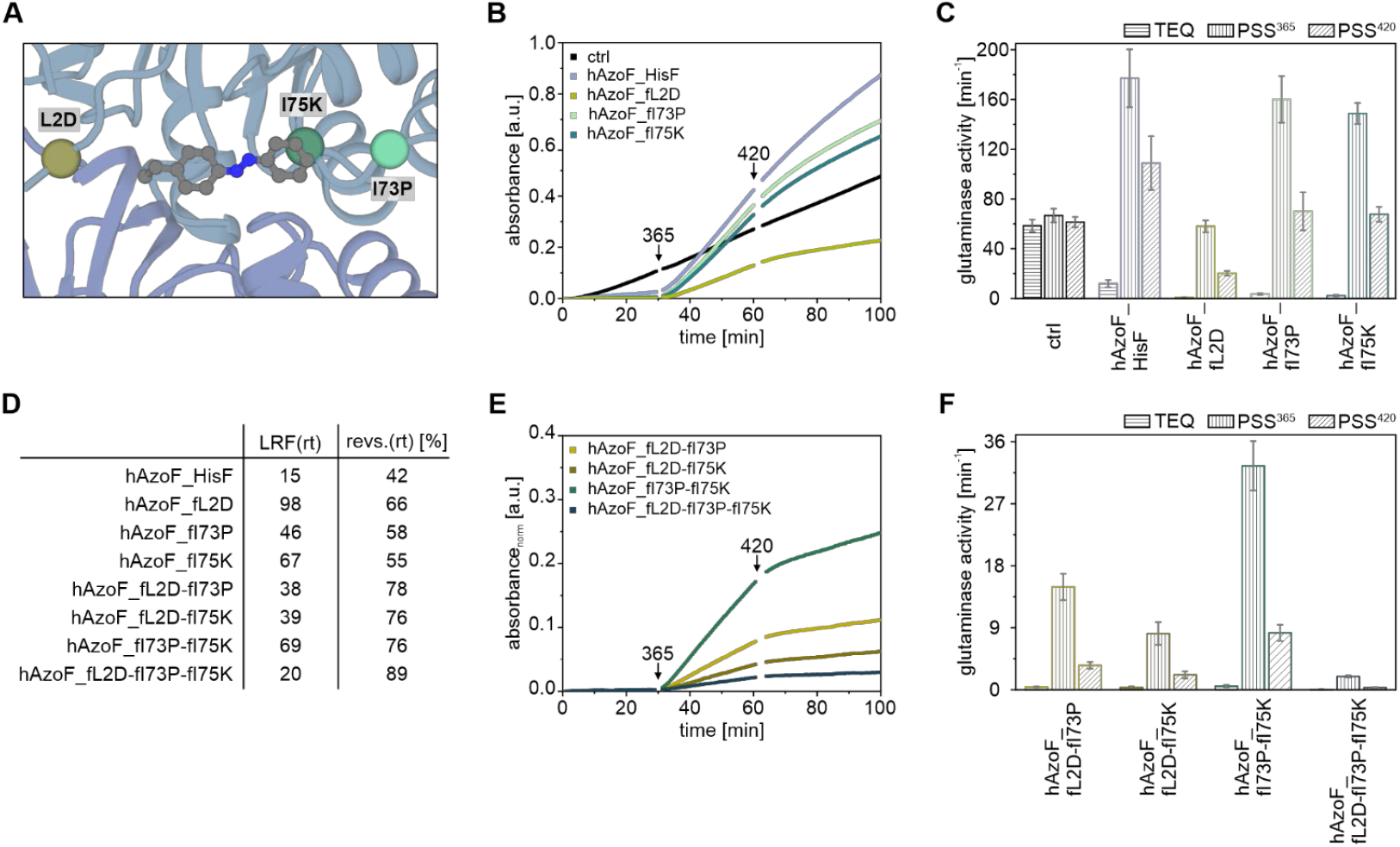
Real-time photocontrol of identified hAzoF_HisF hits. A) Position of each identified mutation at the hAzoF_HisF interface as determined by molecular dynamics simulations (vide infra). B) Mean progress curves, summarizing three technical replicates, from real-time photocontrol experiments for the three single variants in comparison to hAzoF_HisF and HisH_HisF as control (ctrl). C) Mean *v/E*_*0*_ ± SD of technical triplicates for each single variant. D) Summary of mean LRF(rt) and revs.(rt) values. E) Mean progress curves, summarizing three technical replicates and normalized to match the absorbance values in (B), from real-time photocontrol experiments for the combined variants. F) Mean *v/E*_*0*_ ± SD of technical triplicates for each combined variant. *Note:* For exact activity, LRF(rt) and revs.(rt) values of each replicate see **Table S2**. *Irradiation:* 2 s per well with 365 nm for PSS^365^ and 4 s per well with 420 nm for PSS^420^ (individual setup, **Table S8**).

Encouraged by these results, we further generated variants that combined two or all three of the identified mutations and recorded their real-time photocontrol (**Figure 4E–F**; **Table S2**). Although the revs.(rt) appeared to be slightly better (76–89%), the LRF(rt) values for most variants were generally less pronounced than for the single variants (20–69) and the overall glutaminase activity was largely hampered.

### Evaluation of improved photocontrol efficiencies and selection of top hits

To further break down the photocontrol effect in the hAzoF_HisF variants, we recorded glutaminase steady-state kinetics (**Figure S11**). Since our previous studies on HisH_HisF^[26,27]^ and asparaginase-glutaminase^[28]^ based photoxenases indicated that the photocontrol efficiency increases when the enzymes are irradiated in their substrate-bound state instead of their apo state, we established the PSS^365^ and PSS^420^ (successive 365 nm and 420 nm irradiation) of the variants directly after the addition to each reaction mixture. Moreover, we used the same enzyme concentration for PSS^365^ and PSS^420^ reactions to guarantee the successful establishment of each PSS, which is dependent on the photoxenase concentration. Finally, since the activity of TEQ was close to the detection limit in real-time photocontrol experiments, we employed higher enzyme concentrations for the determination of TEQ activity providing more precise LRF values for each variant.

In fact, all variants in TEQ showed reduced catalytic strength compared to hAzoF_HisF with ≤64-fold decreased *k*_*cat*_, ≤3-fold increased *K*_*m*_ and ≤299-fold decreased *k*_*cat*_/*K*_*m*_ values (**Table 1**; **Table S3**). Interestingly, the improvement of the variants results from up to 7-fold increased LRFs(*k*_*cat*_) compared to hAzoF_HisF. LRF(*K*_*m*_) values were throughout worse, and consequently LRF(*k*_*cat*_/*K*_*m*_) values largely comparable to hAzoF_HisF. Notably, while three of the variants showed similar revs.(*k*_*cat*_) values compared to hAzoF_HisF in the range of 50–54%, four variants demonstrated improved values between 59–72%. These results demonstrated that hAzoF_fL2D and hAzoF_fI73P-fI75K were the top two hits as they combine the best LRFs(*k*_*cat*_) of 48 and 99 with high revs.(*k*_*cat*_) of 72% and 66%, respectively.

**Table 1.**
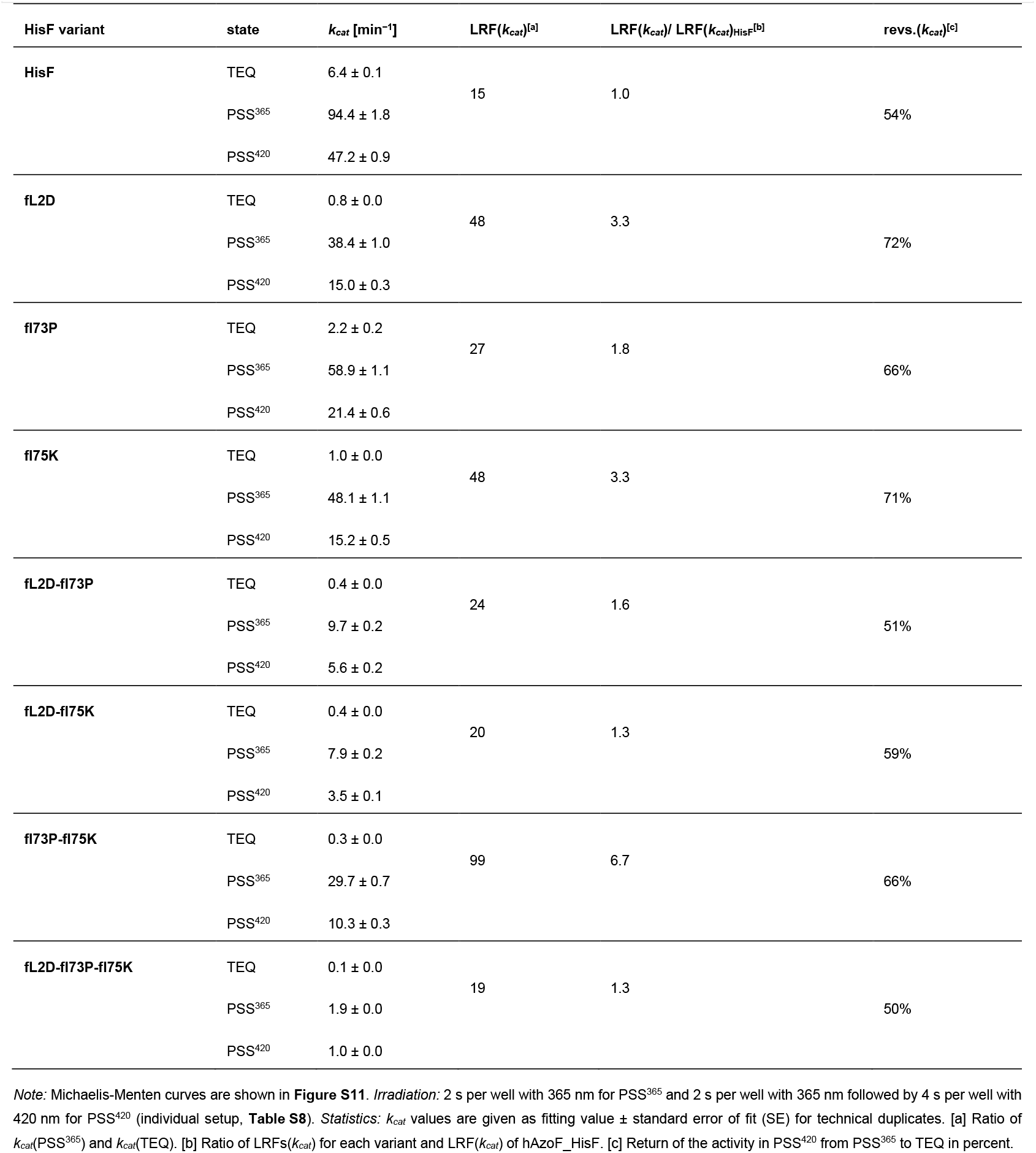
Glutaminase *k*_*cat*_ values and evaluation of photocontrol based on *k*_*cat*_ for hAzoF combined with different HisF variants.

To complete the evaluation of the photocontrol efficiency for the parental construct and our best two hits, we analyzed whether hAzoF_HisF, hAzoF_fL2D and hAzoF_fI73P-fI75K can undergo multiple switching cycles. To this end, we further increased the auxiliary enzyme concentration in our coupled enzymatic assay to facilitate an instant response time, which completely eliminated the lag phase resulting in a pseudo-linear reaction course. While HisH_HisF retained its activity despite repeated 365 nm and 420 nm irradiation cycles, the hAzoF_HisF variants demonstrated fast and efficient photocontrol of HisH activity in form of step-shaped progress curves (**Figure 5A**). We then plotted the glutaminase rate of each segment in these irradiated data sets as well as data sets monitored in the dark (**Figure 5B**) against the corresponding cycle number (**Figure 5C**; **Table S4**). This emphasized that each variant largely maintains its TEQ activity in the dark (light grey) as well as its PSS^365^ activity after each 365 nm irradiation step (orange) and returns to lower activity values after each 420 nm irradiation step (green) rendering photocontrol reversible over multiple cycles. Thereby, the first switching cycle yielded revs.(rt) of ~55% (hAzoF_HisF), ~73% (hAzoF_fL2D), and ~78% (hAzoF_fI73P-fI75K) as expected from steady-state kinetics. However, after the third switching cycle reversibility improved to ~70% (hAzoF_HisF), ~87% (hAzoF_fL2D) and ~95% (hAzoF_fI73P-fI75K). Although we cannot further explain this effect at this point, multiple irradiation steps appear to help enforce a conformational change between the light-induced enzymatic states, and hence the return to a less productive state in the case of hAzoF_HisF variants.

**Figure 5.**
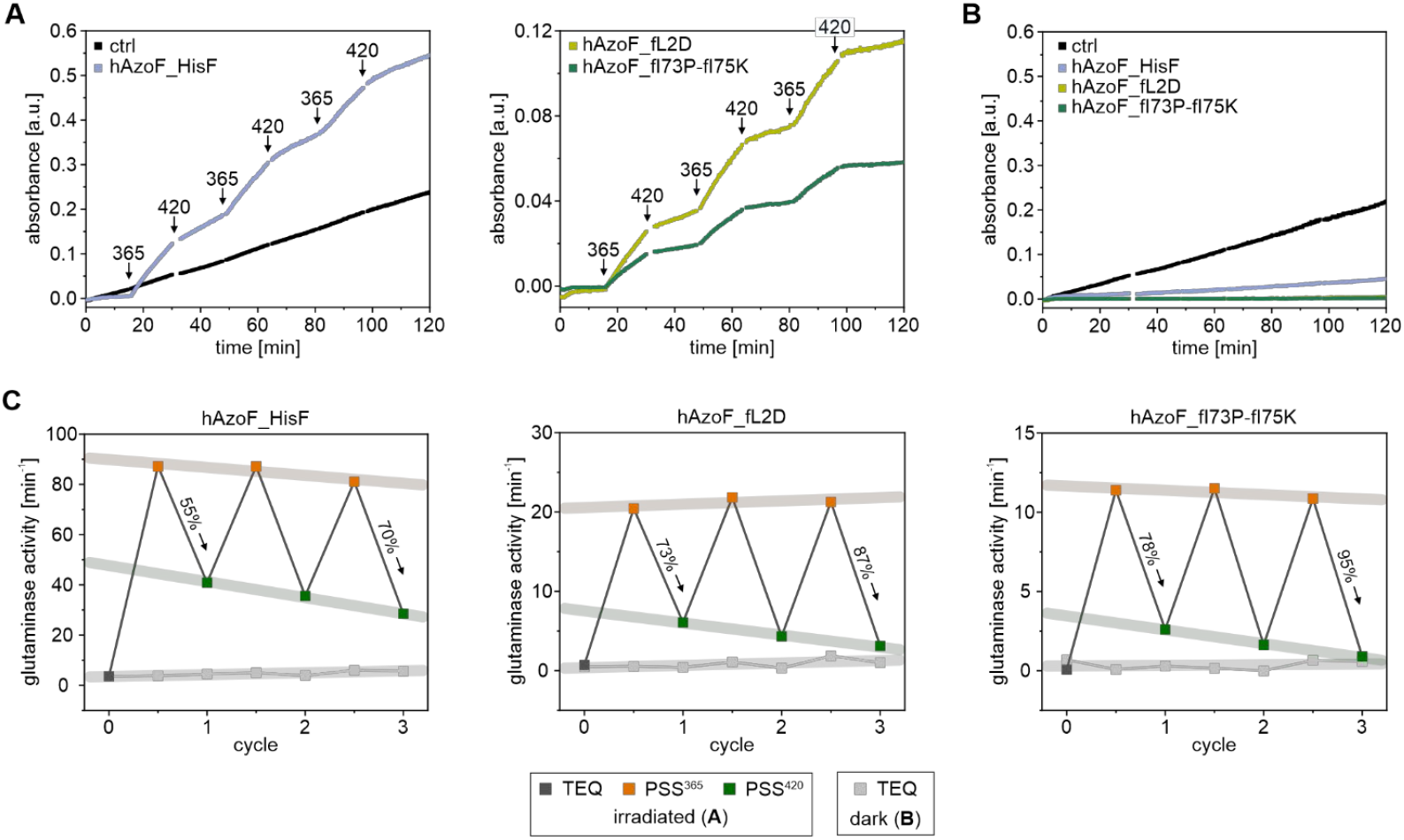
Cycle performance of real-time photocontrol experiments for hAzoF_HisF, hAzoF_fL2D, hAzoF_fI73P-fI75K, and HisH_HisF as control (ctrl). A) Mean progress curves of technical duplicates for real-time photocontrol measurements with three irradiation cycles (365 nm and 420 nm). B) Constant activity of each variant in the dark (TEQ) following a pseudo-linear reaction course over the entire measurement (single measurements). C) Processed real-time photocontrol measurements with mean glutaminase rates (from A and B; **Table S4**) plotted against each irradiation cycle. *Irradiation:* 2 s per well with 365 nm for PSS^365^ and 4 s per well with 420 nm for PSS^420^ (individual setup, **Table S8**).

Altogether, these findings identified two hits that are promising for future application-oriented studies because of their up to ~100-fold improved LRF(*k*_*cat*_) values and good to nearly quantitative reversibility of photocontrol compared to the parental construct.

### Independence of the improved photocontrol effect from photochemistry and complex formation

So far, our findings demonstrated that protein engineering is the method of choice for improving the photocontrol efficiency of photoxenases. However, at the current state-of-the-art these experiments are majorly limited to low or medium throughput screenings because of the low availability of the UAA. Although semi-rational design appears to be suited for this endeavor, residues for randomization need to be carefully selected. In addition to hAzoF_HisF, we set out to improve another HisH_HisF based photoxenase, in which AzoF was incorporated close to the HisF active site (HisH_fAzoF).^[26]^ Interestingly, mutagenesis of residues within an 8 Å radius to AzoF in HisH_fAzoF, similar to the approach for hAzoF_HisF, was ineffective for tuning the photocontrol at the HisH active site. This emphasizes the need of knowledge on the mechanism of photocontrol to establish effective engineering strategies to further boost photoxenases. For this reason, we performed a detailed mechanistic investigation to further clarify the molecular causes of the improved LRF(*k*_*cat*_) values in hAzoF_fL2D and hAzoF_fI73P-fI75K. We started with a biophysical characterization of the two hits in comparison to the parental construct. For this, we initially confirmed that all three hAzoF_HisF variants were properly folded and retained high thermal stability (**Figure S4C**,**D**), although the denaturation midpoint was reduced by 20°C for hAzoF_fI73P-fI75K (*T*_*m*_ = 75°C) compared to hAzoF_HisF and hAzoF_fL2D (*T*_*m*_ ~95°C). Photochemical characterization then demonstrated that AzoF reached only minimally diverging *E*:*Z* ratios in the PSS^365^ and PSS^420^ in all three variants with comparable half-lives *t*_*½*_ of irradiation (**Figure 6**). Moreover, the photoswitch reversibility values upon 420 nm irradiation only vary slightly for hAzoF_HisF (71%), hAzoF_fL2D (67%), and hAzoF_fI73P-fI75K (63%). This further substantiates that the enzyme environment instead of the photochemistry accounts for the better photocontrol efficiencies in the two hits.

**Figure 6.**
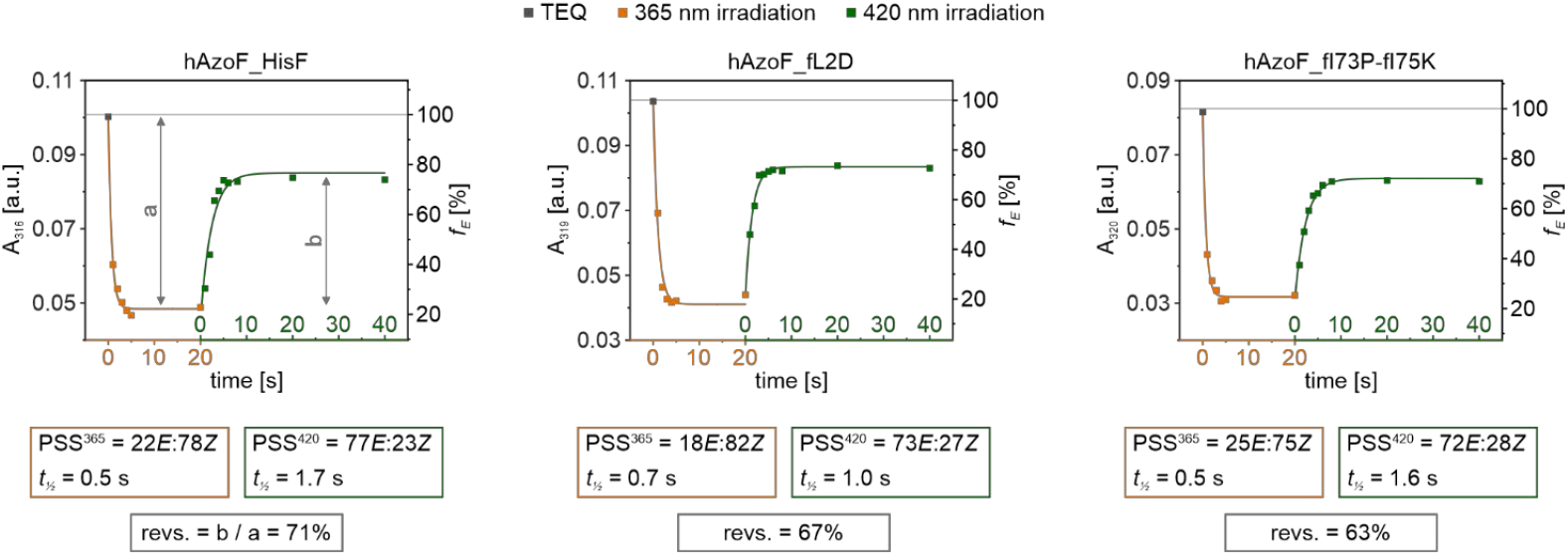
Progress curves following the timely change of the AzoF signal at its π→π* transition maximum upon irradiation (individual setup, **Table S8**), and evaluation of the *E*:*Z* content at each PSS, the half-live of isomerization as obtained by monoexponential fitting, and the reversibility of the photoswitch. Exemplary data of spectra and evaluation of *E*:*Z* ratios are shown in **Figure S12**.

We next analyzed whether heterodimer formation required for the activation of HisH by HisF is affected by the photoswitch of AzoF at the subunit interface. Recent studies indicated that the dissociation constant (*K*_*d*_) of HisH and HisF lies in the low nM range.^[55]^ Hence, we trialed different methods for the highly sensitive analysis of heterodimer formation. While surface plasmon resonance and micro scale thermophoresis experiments were problematic owing to unspecific sticking of the proteins to the matrix and capillaries, fluorescence titration by introduction of the UAA L-(7-hydroxycoumarin-4-yl)ethylglycine (CouA) into each respective HisF subunit showed the necessary sensitivity (**Figure S13**). CouA can be readily incorporated by amber suppression using an orthogonal aaRS/tRNA pair^[56]^ and has been frequently used as a protein-based fluorescent sensor for protein-ligand interactions.^[57–61]^ Incorporation of CouA at position K99 in HisF (HisF^CouA^) resulted in a sensitive signal change by complex formation with HisH. This makes HisF^CouA^ suitable for a comparative analysis of complex formation in TEQ and PSS^365^, although we expect an overall hampered *K*_*d*_ owing to the location of CouA in the interface. We determined CouA fluorescence in the absence or presence of various concentrations of hAzoF in TEQ or PSS^365^ for HisF^CouA^, fL2D^CouA^ and fI73P-fI75K^CouA^ (**Figure S14**). Interestingly, hAzoF_HisF^CouA^ exhibited the same apparent binding constant *K*_*d*_ in its TEQ (~0.19 µM) and PSS^365^ (~0.21 µM; **Table 2**). hAzoF_fL2D^CouA^ demonstrated similar behavior with an overall slightly reduced binding strength (~0.31 µM and ~0.29 µM) compared to hAzoF_HisF^CouA^. Due to a reduced signal change in the presence of hAzoF, we could not analyze hAzoF_fI73P-fI75K^CouA^. However, control measurements with HisH instead of hAzoF showed that while heterodimer formation in HisH_fL2D is hampered with a *K*_*d*_ in the range of ~0.075 µM compared to HisH_HisF (~0.009 µM), it retains high affinity in HisH_fI73P-fI75K (~0.005 µM). Hence, while AzoF incorporation and mutation of HisF minimally affected complex formation, AzoF isomerization instead appeared to maintain a stable complex.

**Table 2.**
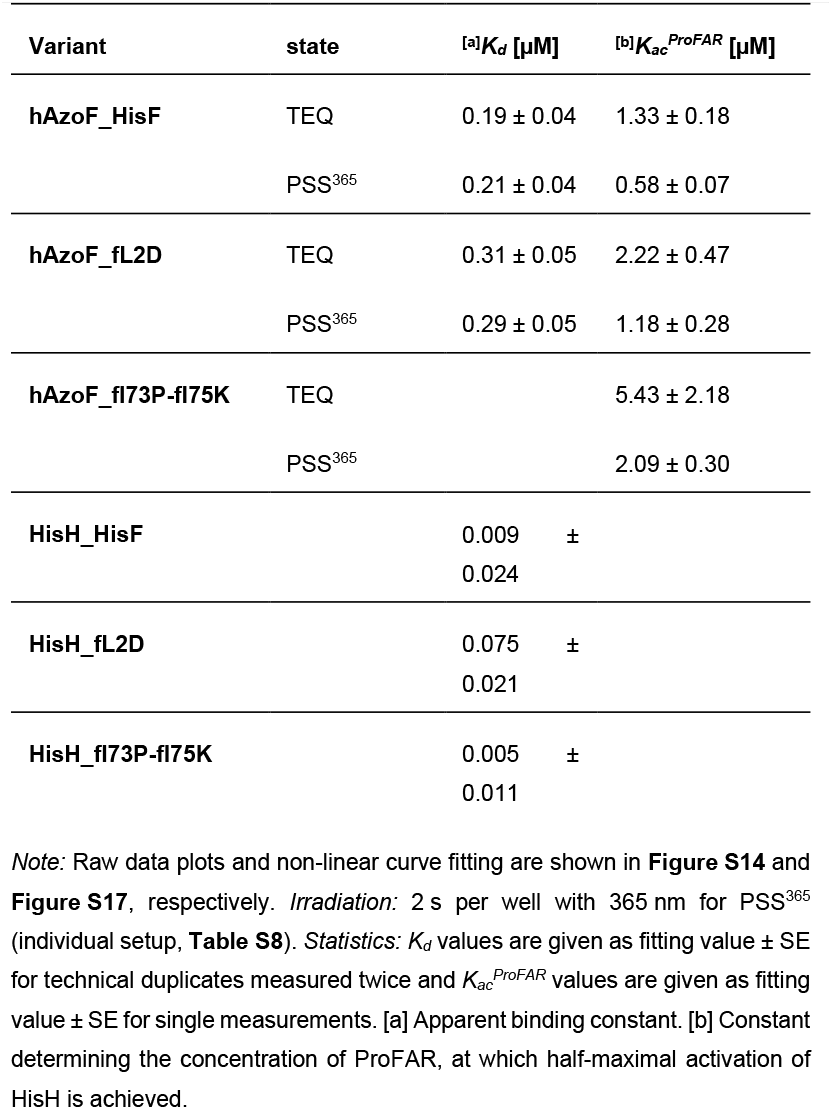
Biophysical analysis of hAzoF_HisF variants.

### Modification of allosteric components in the selected photoxenase hits

Steady-state kinetics of the two best hits revealed that the improvement of their photocontrol efficiency is primarily rooted in an increased LRF(*k*_*cat*_) (**Table 1**). Since HisH_HisF is regulated in nature by a strong V-type allosteric stimulation, we contemplated that increased LRFs(*k*_*cat*_) might hence correlate with a light-dependent modulation of the allosteric machinery in the hAzoF_HisF variants. As a first assessment thereof, we performed real-time photocontrol experiments of glutamine hydrolysis in the absence of an allosteric stimulator (**Figure S15**; **Table S5**). In hAzoF_HisF, TEQ is as dependent on allosteric activation as HisH_HisF showing a similar glutaminase activity (~0.024 min^‒1^), whereas PSS^365^ exhibits constitutive activity (~0.939 min^‒1^) with an LRF(rt) of ~39 and a revs.(rt) of ~47%, which coincides with our previous study.^[27]^ Remarkably, this photocontrol of basal activity was reinforced in hAzoF_fL2D with a 10-fold decrease in glutamine hydrolysis in TEQ (≤0.002 min^‒1^) and a 5-fold increase in PSS^365^ (~0.142 min^‒1^) compared to HisH_HisF, resulting in an LRF(rt) of ~71 and a revs.(rt) of ~69%. In contrast, the effect was reduced in hAzoF_fI73P-fI75K, which obtained a low glutaminase activity in TEQ (~0.004 min^‒1^) and an only slightly increased activity in PSS^365^ (~0.094 min^‒1^) compared to HisH_HisF with an LRF(rt) of ~24 and a revs.(rt) of ~70%. The deviation of activities as well as LRF(rt) values in the absence and presence of ProFAR already suggested that the allosteric mechanism might differ in the three hAzoF_HisF variants.

To further unravel the potential changes in allostery, we next focused on the effect of *E*→*Z* isomerization in the three hAzoF_HisF variants on elements that connect the allosteric and orthosteric site (**Figure 1B**): i) ProFAR binding and loop1 flexibility at the HisF active site, and ii) signal transmission along an allosteric path to the HisH active site. To this end, we employed a combined approach of kinetic investigations and computational evaluations via MD simulations coupled to correlation-based analysis with the SPM tool.^[62,63]^ All simulations were thereby performed starting from the ProFAR- and glutamine-bound catalytically active state (**Figure S16**) with AzoF incorporated either in its *E* or *Z* configuration at position W123 in HisH. The modelled hAzoF_HisF^E/Z^, hAzoF_fL2D^E/Z^, and hAzoF_fI73P-fI75K^E/Z^ complexes thus contain 100% *E*/*Z* isomer and correlate either with the *E*-enriched TEQ or the *Z*-enriched PSS^365^ that we used in our experimental analyses. In the interest of clarity, “h” delineates residues in the HisH subunit and “f” residues in the HisF subunit from now on.

First, we evaluated the binding of ProFAR and its influence on hAzoF by recording glutamine hydrolysis in the absence and presence of various ProFAR concentrations. As a result, we obtained *K*_*acProFAR*_ values, which constitute the ProFAR concentration required to induce 50% hAzoF activity. All variants showed only minimal LRF(*K*_*acProFAR*_) values ≤2 in their PSS^365^ compared to TEQ (**Table 2**; **Figure S17**), which coincides with the results for hAzoF_HisF in our previous study.^[27]^ However, *K*_*acProFAR*_ values were increased two- (hAzoF_fL2D) and four-fold (hAzoF_fI73P-fI75K) compared to hAzoF_HisF. To clarify these effects, we computationally analyzed the ProFAR binding pocket when the orthosteric site in HisH is in the closed catalytically active state for enhanced glutamine hydrolysis. Interestingly, hAzoF_fL2D and hAzoF_fI73P-fI75K showed alternative conformations of loop1 and loop5 compared to hAzoF_HisF (**Figure 7A**). Owing to this difference, fK19 (loop1) and fK179 (loop5) were unable to form a salt-bridge with the phosphate group of ProFAR in the two hits, which is in line with the experimentally increased *K* ^*ProFAR*^. Moreover, the comparison of the non-covalent interactions via NCI analysis^[64]^ established by ProFAR suggests a higher number of stronger interactions (purple-blue color; **Figure S18**) for all variants in their *Z* state, which is consistent with the lower *K* ^*ProFAR*^ in PSS^365^ compared to TEQ. Intrigued by the differences in loop1 and loop5 between the variants, we further evaluated the root mean square fluctuations (RMSF) indicative of changes in flexibility/rigidity (**Figure S19**) of the heterodimers. Loop5 showed an overall rigidification in the two hits but no distinction between the *E* and *Z* states. In comparison, loop1 retained a similar flexibility in the *E* state of all three variants. Interestingly, while loop1 gains flexibility through *E*→*Z* isomerization in hAzoF_HisF, it loses flexibility in hAzoF_fL2D and hAzoF_fI73P-fI75K. Since rigidification of loop1 correlates with higher activities in HisF,^[65]^ we further investigated this effect by determining PRFAR turnover rates (**Extended Text S2; Figure S20–22; Table S6**). In line with the RMSF values, the increase in flexibility of loop1 in hAzoF_HisF coincided with lower HisF activities in PSS^365^ compared to TEQ, whereas its rigidification in both variants resulted in higher HisF activities in PSS^365^. Hence, our findings indicate changes in ProFAR binding and loop1 flexibility upon *E*→*Z* isomerization in the allosteric site of each variant, the latter of which appears to be associated to changes in PRFAR turnover. However, they lack to explain the improvement of LRF(*k*_*cat*_) at the orthosteric site.

**Figure 7.**
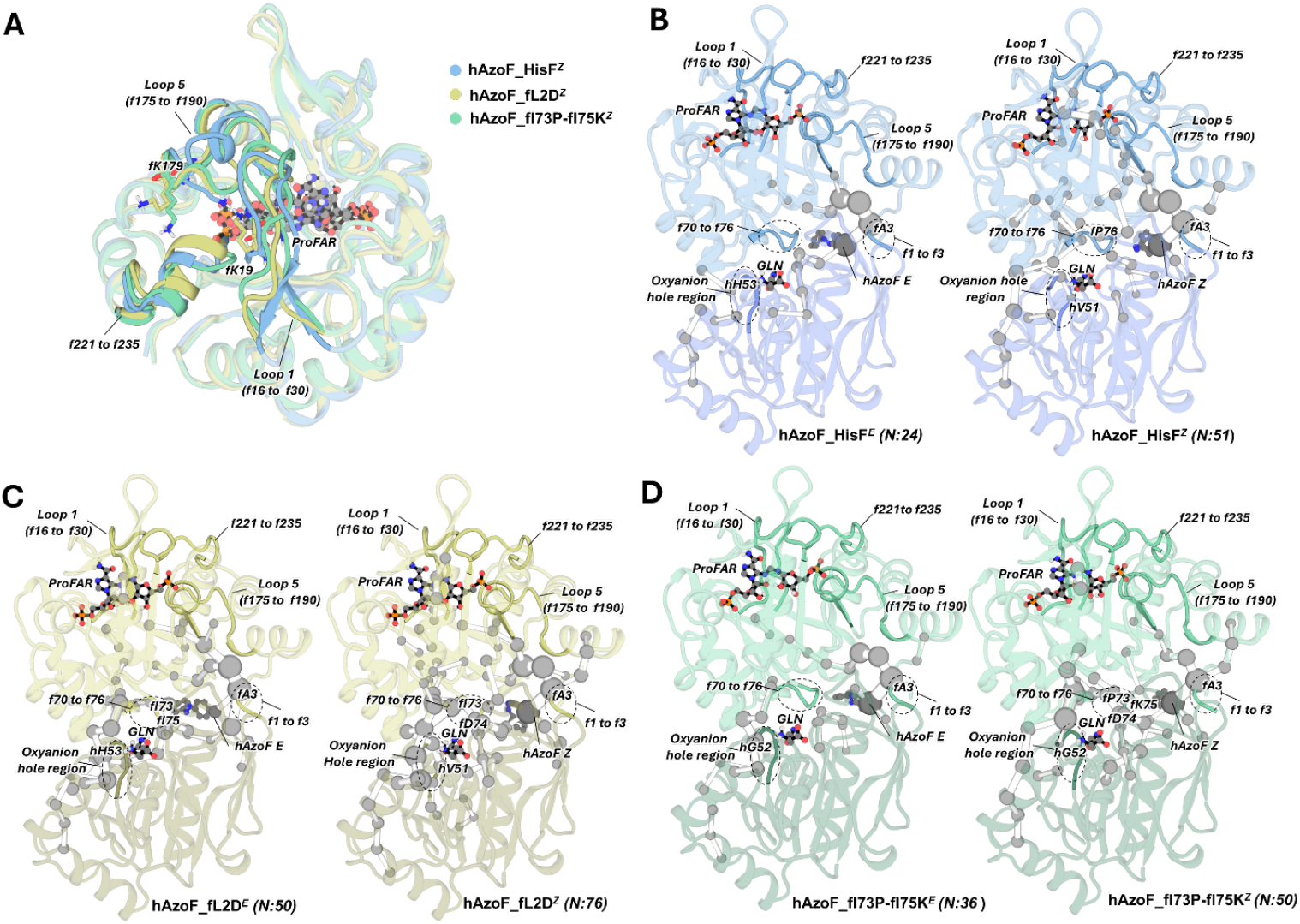
Analysis of the allosteric site and the allosteric signal transmission to the orthosteric site via MD simulations and the correlation-based SPM method. A) Overlay of a representative structure of hAzoF_HisF^Z^ (in blue), hAzoF_fL2D^Z^ (yellow) and hAzoF_fl73P-fl75K^Z^ (green). ProFAR is shown as ball and sticks, and the most important residues contributing to ProFAR stabilization are shown in sticks (cf. **Figure S18**). B–D) Representation of the allosteric communication pathways computed using the SPM tool in the three hAzoF_HisF variants. The number of residues (N) identified in each network is included. The most relevant regions involved in allosteric communication, activation and catalysis are highlighted.

Finally, we were interested in how signal transmission differs in the three hAzoF_HisF photoxenases and whether the catalytically relevant regions are similarly identified in the allosteric pathways of all variants. To this end, we evaluated the results of our correlation-based SPM analysis. For hAzoF_HisF we found a disconnected path between the allosteric site and the orthosteric site with only N = 24 when AzoF adopted the *E* configuration (N is the number of residues identified in the SPM network; **Figure 7B**). In line with an increased glutaminase activity in the *Z* state, the communication between the allosteric and orthosteric site was then enhanced with a continuous path of N = 51 starting at the ProFAR binding site, traversing loop1 and loop5, the interface and the catalytic site in hAzoF, and ending at the oxyanion strand harboring the oxyanion hole. In contrast, the ProFAR pocket is majorly and even entirely decoupled from the orthosteric site in hAzoF_fL2D and hAzoF_fI73P-fI75K, respectively (**Figure 7C,D**). In fact, the nearest SPM positions are located >7 Å from the center of mass of ProFAR. Moreover, while hAzoF_fI73P-fI75K maintains a similar number of conformationally relevant residues compared to hAzoF_HisF (N = 36/50 for *E*/*Z*), hAzoF_fL2D shows a larger number of connected residues for both AzoF configurations (N = 50/76 for *E*/*Z*) that are mostly located at the subunit interface. In conclusion, the observed degree of (de)coupling of the allosteric and orthosteric site in hAzoF_HisF, hAzoF_fL2D, and hAzoF_fI73P-fI75K correlated with the *k*_*cat*_ of each variant state, most likely because of the V-type allosteric connection. In this respect, PSS^365^ of hAzoF_HisF showed the highest turnover number owing to the intact allosteric path, and hAzoF_fL2D as well as hAzoF_fI73-fI75K exhibited increasingly hampered *k*_*cat*_ values owing to the increasingly intensified allosteric disconnection. This demonstrates a complex rewiring of allostery in all three hAzoF_HisF variants.

Although analysis of neither the allosteric site nor the allosteric signal transmission provided a reasonable explanation for the improved LRF(*k*_*cat*_) values at the orthosteric site in HisH, SPM analysis indicated an enhanced conformational network at the orthosteric site and connected subunit interface in both hits compared to the parental construct. Hence, we next performed a detailed analysis of conformational changes between the *E* and *Z* configurations of AzoF at the active site of HisH as well as the subunit interface.

### Correlation of photocontrol improvement in selected photoxenases with altered conformational shifts at the orthosteric site

Glutamine hydrolysis in HisH involves the formation of a covalent thioester intermediate by the release of ammonia and its attack by water to generate glutamate.^[66]^ Recent calculations of reaction energy profiles thereby indicated that conformational transitions associated to the allosteric activation of HisH primarily influence the first part of the reaction (**Figure 8**).^[67]^ This includes the closure of the subunit interface as defined by the angle α between fF120_Cγ_ and hW/AzoF123_Cγ_ as well as hG52_Cα_ (**Figure S23A**).^[27,38,68]^ The closed state facilitates a productive active site through the repositioning of the substrate glutamine. Moreover, the backbone rotation of hV51 in the oxyanion strand from an inactive conformation (dihedral angle: ‒180°<φ<‒90°) to an active conformation (30°<φ<90°) establishes the catalytically decisive oxyanion hole.^[45,69]^ Consequently, we focused our analysis on these two conformational changes as well as the two critical catalytic residues hH178 and hC84 during thioester formation. In step1, hH178 deprotonates hC84 along distance d1 to increase its nucleophilicity for the subsequent nucleophilic attack (step2) on the carboxamide group of the glutamine substrate along distance d2. This generates a negatively charged tetrahedral intermediate, which is stabilized by the oxyanion hole. Acid-catalysis by hH178 along d3 finally initiates the release of the amide moiety of the tetrahedral intermediate as ammonia to create the thioester intermediate (step3).

**Figure 8.**
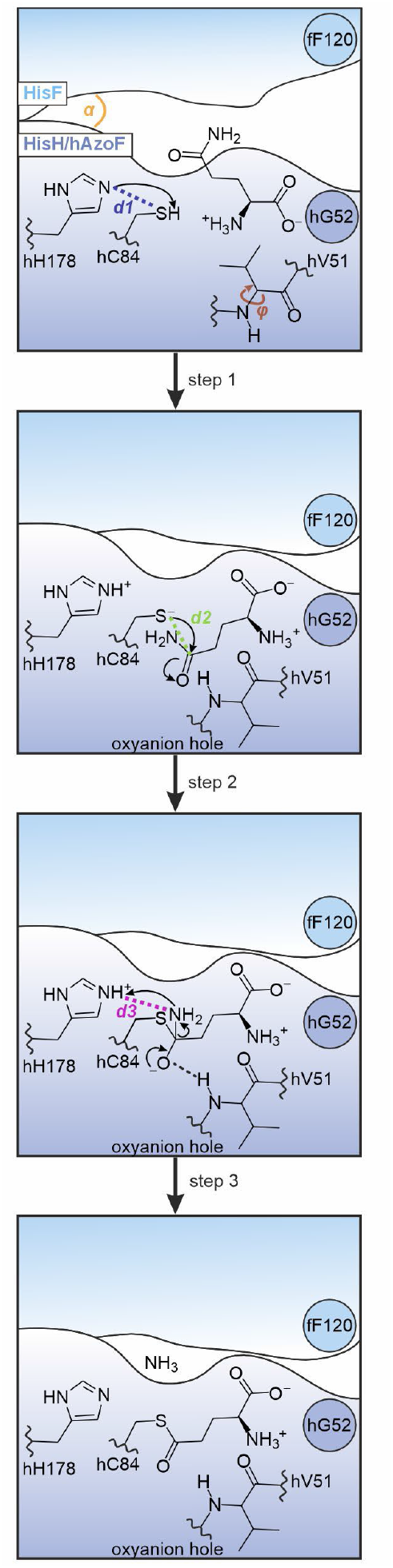
The first three chemical steps of glutamine hydrolysis in HisH. A thioester intermediate is generated and covalently attached to the catalytic residue hC84. The subunit interface angle α, the dihedral angle φ of residue hV51, distance d1 (hH178-hC84), distance d2 (hC84-Gln) and distance d3 (hH178-Gln) that are critical for photocontrol as determined from MD simulations are highlighted.

We started our analysis by reconstructing a conformational landscape based on two (d1 and d3) of the five critical angles/distances defined in **Figure 8**, from our MD simulations. Notably, we observed two distinct conformational populations in the parental construct with AzoF in its *E* configuration (**Figure 9A**). Closer investigation revealed that conformation1 is highly productive with a closed subunit interface (α ~15°), an active oxyanion hole (φ ~77°), a weak^[70]^ hH178-hC84 hydrogen bond (d1 ~3.7 Å), an effective nucleophilic attack distance hC84-Gln (d2 ~3.2 Å), and a moderate^[70]^ hH178-Gln hydrogen bond (d3

**Figure 9.**
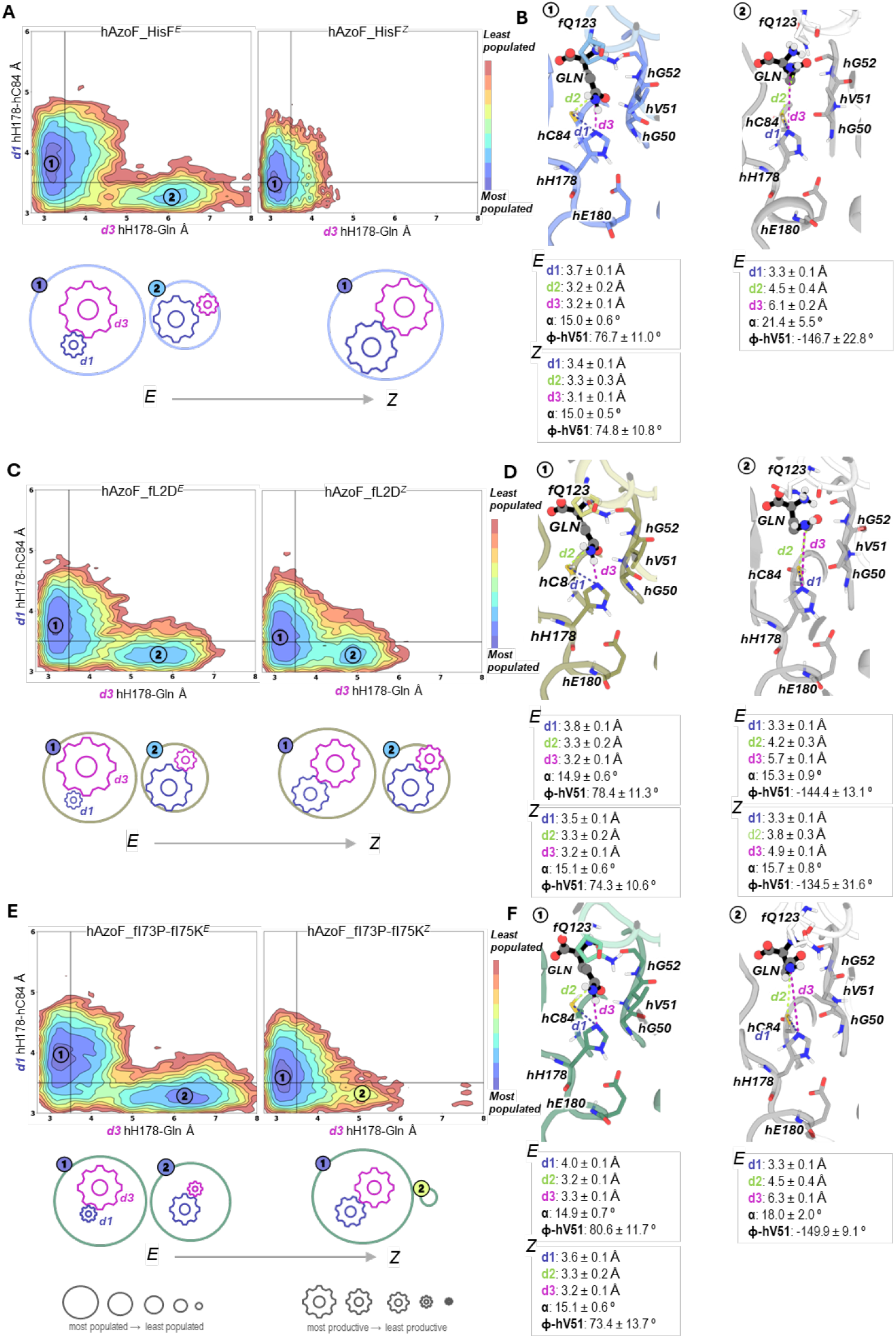
Reconstructed conformational landscapes for selected photoxenases. A, C, E) Conformational landscapes based on d1 and d3 distances for hAzoF_HisF, hAzoF_fL2D and hAzoF_fI73P-fI75K in both *E* and *Z* configurations. Most populated conformations are colored in blue, whereas red is used for the least populated ones. In most cases, two conformations are identified: the catalytically competent 1 presenting short d1 and d3 and the unproductive 2. Below each conformational landscape a qualitative representation with spheres and gears is included: the sphere size denotes the population of the state, whereas the gear sizes indicate the productivity of the state as indicated by d1 and d3. B, D, F) A representative structure of each conformation1 and 2 is shown together with the average values of d1–d3 (in A), intersubunit closure angle (in degrees), and dihedral of the oxyanion residue hV51 (in degrees). The most important residues are shown as sticks, and glutamine substrate is represented in spheres and black sticks.

~3.2 Å; **Figure 9B**; **Figures S23–S27**). In contrast, conformation2 appeared to be unproductive with an open subunit interface (α ~21°), an inactive oxyanion hole (φ ~‒147°), and similar or weakened catalytic distances d1–d3. The long d2 (~4.5 Å) and d3 (~6.1 Å) distances explored were mostly related to the repositioning of the substrate glutamine due to both conformational changes at the subunit interface and the active-to-inactive transition of the oxyanion strand. This favored the establishment of a hydrogen bond between the carbonyl oxygen of glutamine and hG52 (3.0 ± 0.2 Å), which has been described in a previously reported inactive crystal structure of HisH_HisF (PDB-ID: 3zr4).^[37]^ Remarkably, isomerization of AzoF to *Z* led to a slightly improved productivity of conformation1 with a reduced d1 (~3.4 Å; **Figure S25**) and d3 (~3.1 Å; **Figure S26**) distances, and to the complete loss of conformation2 (**Figure 9A,B**). This change between hAzoF_HisF^E^ and hAzoF_HisF^Z^ is comparable to the observed factors that define photocontrol in optogenetic photoreceptors.^[33,71,72]^ Hence, similar to photoreceptors, LRF(*k*_*cat*_) of hAzoF_HisF can be described by the altered difference in population of conformation1 and conformation2 between the *E* and *Z* state, meaning the size of the resulting conformational shift, and the productivity of the more productive conformation1 (shorter d1 and d3 distances). In **Figure 9**, a qualitative representation of the relative populations of both conformations is shown with weighted spheres, whereas productivity of each conformation is represented with gears (one for each d1/d3 distance and scaled based on the distance value).

Next, we compared hAzoF_fL2D to the parental construct using the conformational shift and the productivity of conformation1. The difference to hAzoF_HisF appeared to be mainly based on a changed equilibrium towards higher proportions of the unproductive conformation2, and thus towards higher conformational heterogeneity, in both the *E* and *Z* state (**Figure 9C**). This is in line with the largely disconnected allosteric paths (**Figure 7**) and the overall reduced *k*_*cat*_ (**Table 1**) of hAzoF_fL2D compared to hAzoF_HisF. Moreover, the size of the conformational shift between the *E* and *Z* state was significantly reduced, since conformation2 is strongly populated in hAzoF_fL2D^Z^. Interestingly, the degree of productivity of conformation1 in hAzoF_fL2D^Z^ is also lower as compared to hAzoF_HisF^Z^ due to the displacement of the minimum towards longer d1 distances (**Figure 9D**). This is again in line with the reduced *k*_*cat*_ for hAzoF_fL2D^Z^ (**Table 1**). Hence, both factors, the size of the conformational shift and the productivity of conformation1 are actually reduced compared to hAzoF_HisF, which appears to be contradictory to an improved LRF(*k*_*cat*_). However, the photoreceptor model in optogenetics states that an increase in heterogeneity is a third factor for the improvement of photocontrol (**Figure S28**), which appears to be the case for hAzoF_fL2D. Importantly, we have to note that the N-terminal position of the mutation fL2D makes the interpretation of the MD data and therefore the correlation to experimental data more challenging.

Similarly, hAzoF_fI73P-fI75K showed a changed equilibrium towards even higher proportions of the unproductive conformation2 in its *E* state compared to hAzoF_HisF and hAzoF_fL2D (equally sized spheres; **Figure 9E**) increasing its conformational heterogeneity and hence LRF(*k*_*cat*_). Again, this correlates with the completely disconnected allosteric paths (**Figure 7**) and the overall decreased *k*_*cat*_ (**Table 1**). Although, conformation2 is highly populated in hAzoF_fI73P-fI75K^E^, only remnants of it were detected in hAzoF_fI73P-fI75K^Z^. As a result, the size of the conformational shift between the *E* and *Z* state was increased compared to hAzoF_HisF, which further explains the large LRF(*k*_*cat*_). Furthermore, productivity of both conformations as a measure of the interface configuration, the oxyanion hole conformation and the three catalytic distances d1–d3 remained similar to those in hAzoF_HisF. However, the lower *k*_*cat*_ value observed for hAzoF_fI73P-fI75K^Z^ is explained by the displacement of the minimum towards slightly longer d1 distances. As an additional measure, we found that fK75 establishes stable cation-π and CH-π interactions with AzoF primarily in conformation1, while longer cation-π and CH-π distances are explored in conformation2 (**Figure S29**). This additional interaction might be key for the changes in the allosteric path and the conformation shift, and hence for the improved LRF(*k*_*cat*_) in hAzoF_fI73P-fI75K.

In conclusion, the two best hits demonstrated a change in the size of the conformational shift, the productivity of conformation1 and the conformational heterogeneity upon *E*→*Z* isomerization. This not only clarifies the molecular causes of the improved LRF(*k*_*cat*_) values but also suggests that future semi-rational design approaches of photoxenases might be most effective by targeting catalytically relevant domains or directly the active site.

## Conclusion

In this study, we have demonstrated that the photocontrol efficiency of hAzoF_HisF can be improved by introducing mutations at the heterodimer interface close to AzoF, which led to significantly increased LRF values with near-quantitative reversibility. By combining experimental steady-state kinetic data with the correlation-based SPM method and computational reconstruction of conformational landscapes we elucidated the photocontrol mechanism of hAzoF_HisF and how this is altered in the best two hits. The analysis of the allosteric networks indicates a more decoupled communication between both subunits leading to an overall lower catalytic activity in the hits. We further observed that conformational changes at the orthosteric site are directly correlated with the photocontrol improvement established in our semi-rational design experiments. This particularly involves the size of the conformational shift induced upon AzoF isomerization, the productivity of the active conformation and the conformational heterogeneity. In sum, these findings suggest that amino acid exchanges close to the orthosteric site are highly effective to customize photocontrol in photoxenases. Finally, while the parental hAzoF_HisF and the best two hits already achieved pronounced LRF(*k*_*cat*_) values of 15, 48 and 99, the changes in the conformational shifts appeared to be rather small. This observation implies that further semi-rational design, and directed evolution in general, may optimize the conformational shift of these hAzoF_HisF photoxenases and hence maximize the LRF values beyond 100, which significantly emphasizes the power of photoxenase engineering.

## Acknowledgements

The authors thank Reinhard Sterner for his valuable support throughout the project as well as Volker Sieber for mentoring S.M. Furthermore, the authors highly appreciate the excellent technical assistance of Sabine Laberer, Jeannette Ueckert, Klaus Tiefenbach, Elena Adlmanninger, and Ann-Kathrin Wolff. A special thanks to the team of the electronics workshop at the University of Regensburg for the construction of the irradiation setups. Moreover, A.B. thanks Patricia Luckner and Eduard Hochmuth for their help with mass spectrometry. This work was funded by the Deutsche Forschungsgemeinschaft (STE 891/12– 2). We thank the Generalitat de Catalunya for the consolidated group TCBioSys (SGR 2021 00487), Spanish MICIN for grant projects PID2021-129034NB-I00 and PDC2022-133950-I00. S.O. is grateful to the funding from the European Research Council (ERC) under the European Union’s Horizon 2020 research and innovation program (ERC-2015-StG-679001, ERC-2022-POC-101112805, ERC-2023-POC-101158166, and ERC-2022-CoG-101088032), and the Human Frontier Science Program (HFSP) for project grant RGP0054/2020. J.S. was supported by the Spanish MINECO for a PhD fellowship (PRE2022-105114), and M. E by the ERC projects ERC-2015-StG-679001, ERC-2022-POC-101112805, ERC-2022-CoG-101088032.

## Notes

### Competing Interest Statement

The authors have declared no competing interest.

