## Supporting Information for "Boosting reversible photocontrol of a photoxenase by an engineered conformational shift"

Table S9. 33
Experimental section 34
Chemicals and other resources 34
Irradiation devices 34
Synthesis of AzoF 35
Biosynthesis of ProFAR 36
Production of auxiliary enzymes HisG/IE and HisA 36
Site-directed mutagenesis and Golden Gate cloning 37
Subcloning of pEVOL-AapF 37
Subcloning for the split-GFP assay 38
Scanning mutagenesis of HisF and hAzoF 38
Production and purification of HisF and HisH variants without UAA 38
Production and purification of hAzoF and hAapF 39
Production and purification of $HisF^{CouA}$ variants 40
Production and purification of GFP1–10 40
Screening of DNA libraries 41
Real-time photocontrol activity assay 42
Z'-factor determination 43
Alanine screen of ProFAR-stimulated glutaminase activity 44
Cycle performance of enzymatic photocontrol 44
Steady-state kinetics of ProFAR-stimulated HisH-activity 45
UV/Vis analysis 46
Estimation of *E*:*Z* ratios from UV/Vis spectra 46
Fluorescence analysis of complex formation with $HisF^{CouA}$ 47
Determination of $K_{ac}^{ProFAR}$ values 48
Determination of $K_{ac}^{hAzoF}$ values and $k_{cat}^{app}$ of HisF 48
Determination of $K_{ac}^{HisF}$ values and $k_{cat}^{app}$ of ProFAR stimulated hAzoF 49
Circular dichroism analysis 49
Differential scanning calorimetry 49
Mass spectrometry 50
Molecular modelling system preparation 51
MD simulation details 51
Conformational landscape (CL) reconstruction 52
References 54

#### Extended Texts

**Extended Text S1.** Establishment and implementation of an activity-based screening assay

For evaluation of photocontrol efficiencies, we established a reliable screening assay. To this end, we employed a standardized split-GFP assay[1–3] that allowed us to quantify the amount of produced enzyme via a C-terminally attached s11-tag on either hAzoF (hAzoF$^{s11}$) or HisF (HisF$^{s11}$) and our colorimetric detection assay for glutamine turnover (**Figure 5**). To optimize the signal-to-noise ratio when using enzyme-containing cell lysates in these assays, we applied a heat step at 60°C, which separates the thermostable hAzoF or HisF from most *E. coli* proteins. Spiking an *E. coli* lysate with purified hAzoF_HisF particularly showed that the background glutaminase signals, which impact the apparent photocontrol strength after irradiation, could be significantly reduced by the heat step (**Figure S7**). We then optimized expression conditions, for which we only employed hAzoF$^{s11}$, since expression of hAzoF mutants, which involves UAA incorporation, is more critical regarding misincorporation and yields than expression of HisF variants. Recently, various *E. coli* strains were engineered specifically for the production of UAA-containing proteins.[4–7] Strain B95.ΔAΔfabR was particularly appealing for our screening as it is based on the commonly used *E. coli* expression strain BL21 (DE3) and therefore compatible with pET plasmids.[8] In B95.ΔAΔfabR, 95 of the 273 UAG codons, which are located at the end of genes and which are important for bacterial growth, are replaced by other stop codons. Additionally, release factor 1 (RF1), which recognizes UAG at the ribosome, is eliminated to facilitate an improved UAA incorporation without compromising the reproductive strength of the host cells. We followed the production of hAzoF$^{s11}$ in B95.ΔAΔfabR 95 in the presence and absence of AzoF in LB, TB, and M9$^{-}$ medium via SDS-PAGE and split-GFP detection (**Figure S8A–B**). Signals for hAzoF$^{s11}$ were only detectable after expression in LB and TB, but also appeared after expression in the absence of AzoF indicating high amounts of misincorporated natural amino acids. Since native MS experiments in our previous studies could not determine misincorporations after expression of UAA-containing enzymes in BL21 Gold (DE3) cells,[9] we subsequently switched to this strain. Again, hAzoF$^{s11}$ only appeared in measurable amounts after expression in LB and TB medium (**Figure S8C**). Moreover, in the absence of AzoF we only detected signals on the level of our background control implying negligible amounts of misincorporation (**Figure S8D**). Doubling of the AzoF concentration in the medium could not further improve the yields (**Figure S9A–B**). With this, we chose expression in TB medium supplemented with 0.4 mM AzoF with subsequent heat step purification at 60°C for our screening in microtiter plates. The same conditions also facilitated the production of high amounts of $HisF^{s11}$ (**Figure S9C**).

In the next step, we set up the activity-based screening of LRFs. For this, we used a standardized approach in our lab, in which we start the glutaminase reaction by the addition of purified enzyme (or lysate) in its TEQ, spectrophotometrically follow its progress using our colorimetric assay, and then determine the photocontrol effect upon 365 nm and subsequent 420 nm irradiation in real-time ("real-time photocontrol"; **Figure S9D**). The previous optimization of expression yields and signal-to-noise ratio by introducing a heat step resulted in only negligible glutaminase background whilst obtaining an LRF of ~4 with a reversibility of ~45% (**Figure S9D**). The unexpectedly low LRF could be traced back to the low concentration of glutamate oxidase as auxiliary enzyme, which mitigated the response time of the coupled enzymatic assay. Statistic evaluation determining excellent Z′-factors (0.52 and 0.62 for the measurement of LRF values after 365 nm and 420 nm irradiation) further confirmed the reliability of this assay for the screening of improved photocontrol efficiencies. Furthermore, steady-state kinetics of hAzoF_HisF and $hAzoF\_HisF^{s11}$ showed that glutaminase activity was largely retained in $hAzoF\_HisF^{s11}$, although the s11-tag is positioned at the subunit interface (**Figure S9E**). In contrast, the s11-tag in hAzoF is far from the interface and not expected to interfere with the allosteric activation of glutamine hydrolysis.

Finally, we screened all 342 hAzoF_HisF variants in 18 enzyme libraries. For this, we equally distributed the 19 $hAzoF^{s11}$ or $HisF^{s11}$ variants from one library in four biological replicates throughout the 96-well microtiter plate for heterologous gene expression (**Figure S10**). Moreover, to compensate for the inhomogeneous light-intensity in our in-house developed irradiation chamber (for details see: Experimental Section in the Supporting Information), we prepared a second plate with the same clones inversely positioned. Glutamine hydrolysis was then initiated by addition of the harvested lysates, which were supplemented with either HisF or hAzoF to form the $hAzoF^{s11}\_HisF$ or $hAzoF\_HisF^{s11}$ variants, to the coupled enzymatic assay following the same microtiter plate patterns.

**Extended Text S2.** Experimental evaluation of loop1 flexibility influence on activities in hAzoF_HisF variants.

The result that loop1 in hAzoF_HisF is more rigid in the *E* state than in the *Z* state, whereas it is more flexible for hAzoF_fL2D and hAzoF_fI73P-fI75K, puzzled us (**Figure S19**). Loop1 is decisive for the cyclase reaction in HisF fixating PRFAR in the active site.[10,11] Hence, higher rigidity of loop1 should correlate with higher PRFAR turnover. However, our previous steady-state kinetics of hAzoF_HisF indicated that PRFAR turnover is higher in the *Z*-enriched $PSS^{365}$ than in the *E*-enriched TEQ.[12,13] To decipher this discrepancy we chose an experimental setup, in which we followed PRFAR turnover of HisF, fL2D, and fI73P-fI75K spectrophotometrically (**Figure S20**) in the absence and presence of various hAzoF concentrations (**Figure S21**). As a result, we obtained two values, an apparent $k_{cat}$ ($k_{cat}^{app}$) and the hAzoF concentration required to induce 50% of HisF activity ($K_{ac}^{hAzoF}$; **Table S6**). While $k_{cat}^{app}$ increases up to 3-fold in $PSS^{365}$ and decreases in $PSS^{420}$ for both evolved variants, it instead declines 1.3-fold in $PSS^{365}$ and increases again in $PSS^{420}$ for the parental construct. This trend is consistent with the observed flexibility/rigidity of loop1 in the *E* and *Z* state (**Figure S19**). The overall lower LRF values are also in line with our previously determined LRF values for PRFAR turnover in hAzoF_HisF.[12,13] Remarkably, the activity behavior of hAzoF_HisF is reversed at lower hAzoF concentrations matching the findings of our previous study, which were performed at 0.1 µM.[12] This effect is explained by a light-induced change (3-fold) in $K_{ac}^{hAzoF}$, which was only observed for the parental construct but not for the two hits. This response to irradiation of $K_{ac}^{hAzoF}$ appears to contradict the results of the $K_d$ of heterodimer formation (**Table 2**), which indicated no change in complexation upon irradiation. However, $K_{ac}^{hAzoF}$ involves various other equilibrium constants in addition to the $K_d$ of heterodimer formation including the $K_d$ of glutamine binding and conformational equilibria of allosteric signal transduction.

To check whether the inverted flexibility behavior of loop1 in hAzoF_HisF might also affect the photocontrol effect of the ProFAR-stimulated glutaminase reaction, we followed glutamine hydrolysis of hAzoF in the absence and presence of various concentrations of HisF (**Figure S22**). As a result, we again obtained an apparent $k_{cat}$ ($k_{cat}^{app}$) and the HisF concentration required to induce 50% of HisF activity ($K_{ac}^{HisF}$). Importantly, the $k_{cat}^{app}$ values confirmed that hAzoF_HisF exhibits higher glutaminase turnover in $PSS^{365}$ (~86.5 $min^{-1}$) than in TEQ (~5.7 $min^{-1}$). Consistent with the $K_{ac}^{hAzoF}$ measurements, we also found a decrease of $K_{ac}^{HisF}$ in $PSS^{365}$ (~0.003 µM) compared to TEQ (~0.021 µM) with an LRF($K_{ac}^{HisF}$) of 7. Notably, both $K_{ac}^{HisF}$ values were up to 23-fold reduced compared to the

$K_{ac}^{hAzoF}$ values. The complexity of equilibria that are involved in the definition of $K_{ac}^{hAzoF}$ and $K_{ac}^{HisF}$ makes it difficult to pinpoint the exact cause of this discrepancy. However, the clear differences in size and LRF suggests that $K_{ac}^{hAzoF}$, $K_{ac}^{HisF}$, and $K_d$ of complexation are unequal to each other.

In conclusion, the changes in loop1 flexibility result in differences in PRFAR turnover at the HisF active site, but appear to not influence glutamine hydrolysis at the orthosteric site in HisH upon stimulation with ProFAR.

### Supplementary Figures

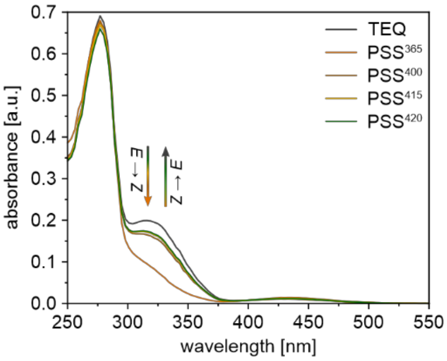

**Figure S1.** UV/Vis analysis of hAzoF_HisF in TEQ and after irradiation with 365 nm for 30 s and with 400 nm, 415 nm, and 420 nm for 2 min (individual setup, **Table S8**) to establish the respective PSSs.

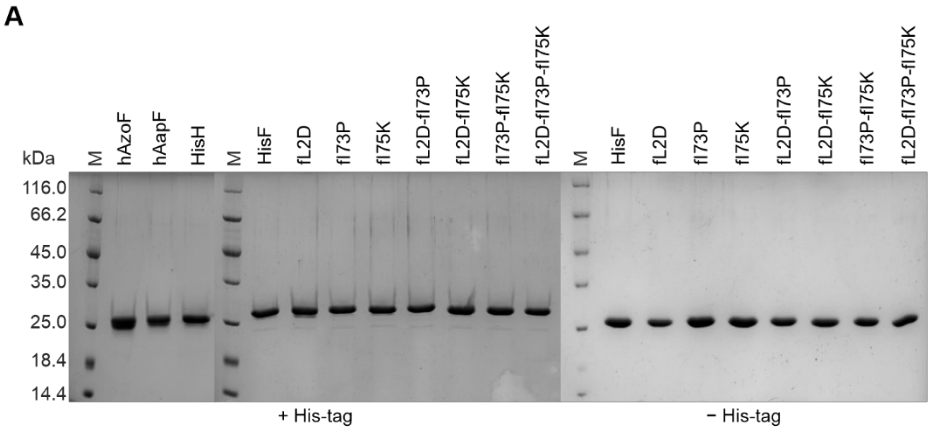

**B**

| | + His mg/L | − His mg/L |
|---|---|---|
| hAzoF | 17 | |
| hAapF | 18 | |
| HisH | 14 | |
| HisF | 28 | 18 |
| fL2D | 32 | 40 |
| fI73P | 15 | 23 |
| fI75K | 20 | 28 |
| fL2D-fI73P | 23 | 13 |
| fL2D-fI75K | 22 | 25 |
| fI73P-fI75K | 24 | 15 |
| fL2D-fI73P-fI75K | 23 | 21 |

**Figure S2.** Purities (A) and yields (B) of enzymes after heterologous gene expression in *E.coli*. A) SDS-PAGE of 3 µg enzyme confirming >95% purity of all produced variants. HisH variants and HisF variants with (+) N-terminal $His_6$-tag are all at the expected heights of ~25 kDa and ~30 kDa, respectively. HisF variants without (−) $His_6$-tag are all at the expected height of ~28 kDa. B) Production yields of enzymes in mg per liter expression medium.

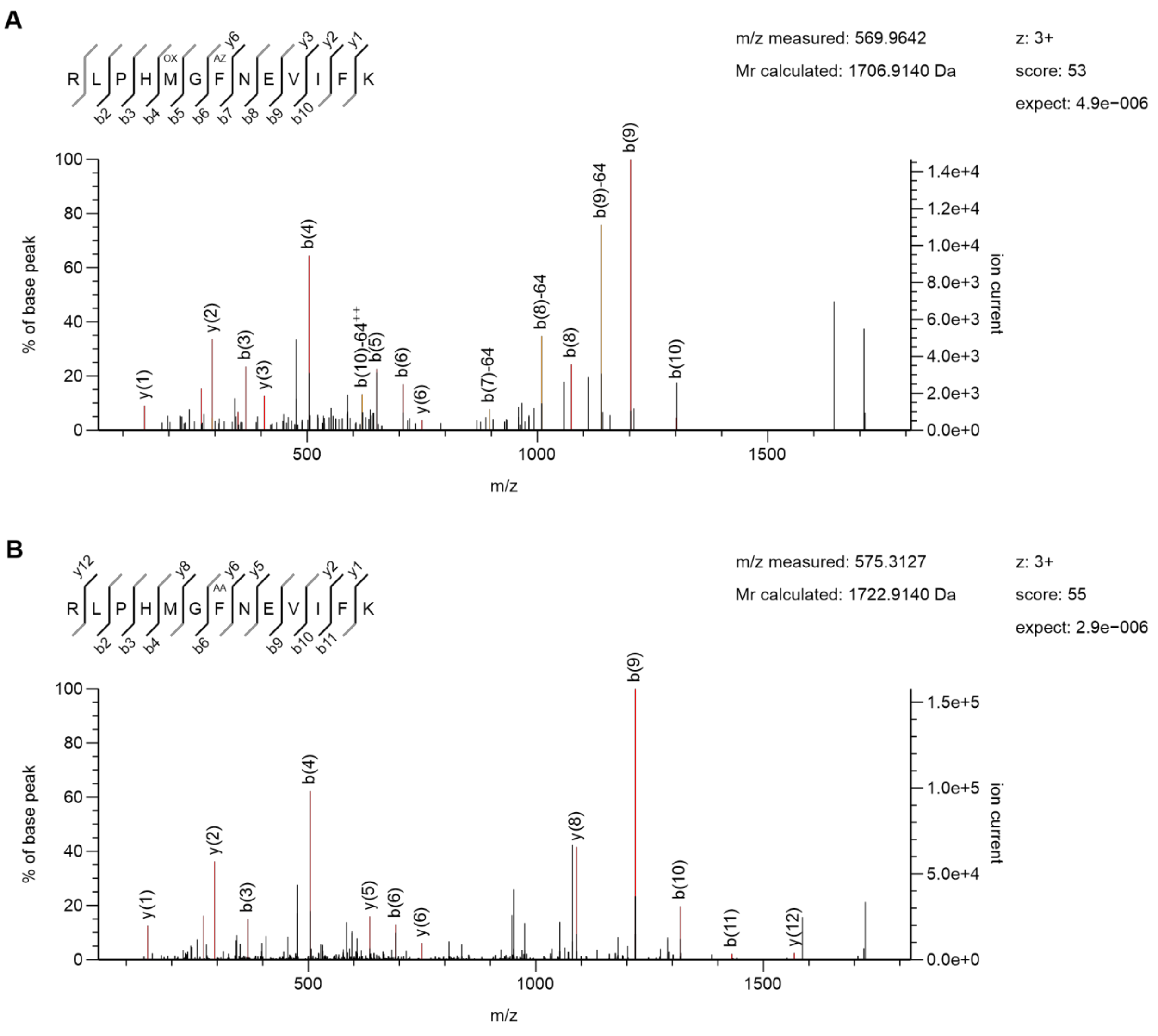

**Figure S3.** Confirmed incorporation of AzoF (A) and AapF (B) at position W123 in HisH by tryptic digest and analysis via liquid chromatography coupled to mass spectrometry (LC-ESI-MS/MS). Both panels show the fragment spectrum and an overview of all measured fragments (highlighted in red) of the semi tryptic peptide harboring AzoF (AZ) or AapF (AA) at position 123. Fragments highlighted in orange indicate a loss of $CH_3SOH$ of oxidized methionine during sample preparation.

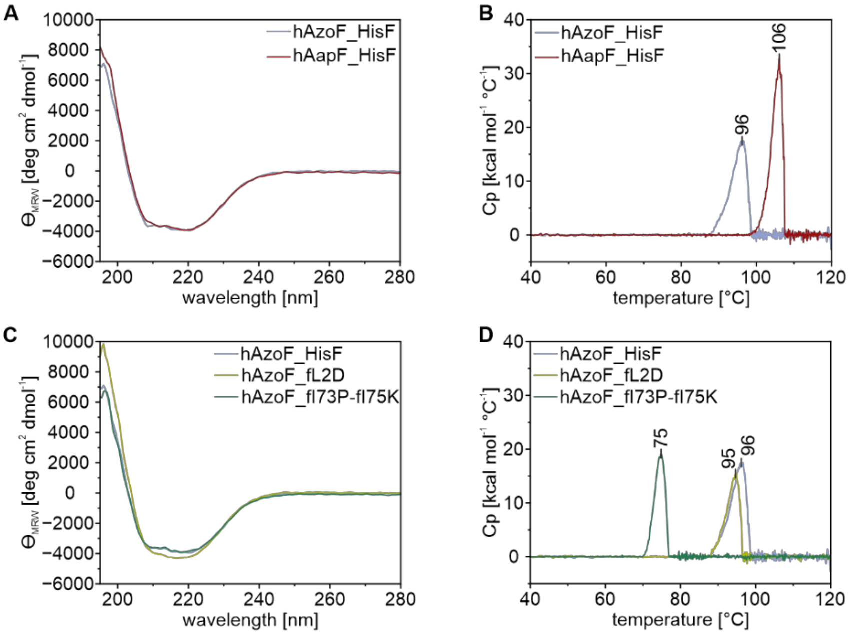

**Figure S4.** Structural integrity and thermal stability of hAzoF and hAapF in complex with different HisF variants. A) Far-UV circular dichroism (CD) spectra of 30 μM hAzoF_HisF and 30 μM hAapF-HisF in 50 mM KP (pH 7.5) indicate an intact overall fold. B) Differential scanning calorimetry (DSC) measurements of 20 μM hAzoF_HisF and 20 μM hAapF_HisF in 50 mM KP (pH 7.5), 100 mM NaCl demonstrate high thermal stability of the complexes with denaturation midpoints $T_m$ of 96°C for hAzoF_HisF and 106°C for hAapF_HisF. C) Far-UV CD spectra indicate an intact overall fold for all hAzoF complexes identified as hits [30 μM enzyme complex in 50 μM KP (pH 7.5)]. D) DSC measurements demonstrate an unaffected high $T_m$ of 95°C for hAzoF_fL2D and a reduced $T_m$ of 75°C for hAzoF_fI73P-fI75K [20 μM complex in 50 mM KP (pH 7.5), 100 mM NaCl].

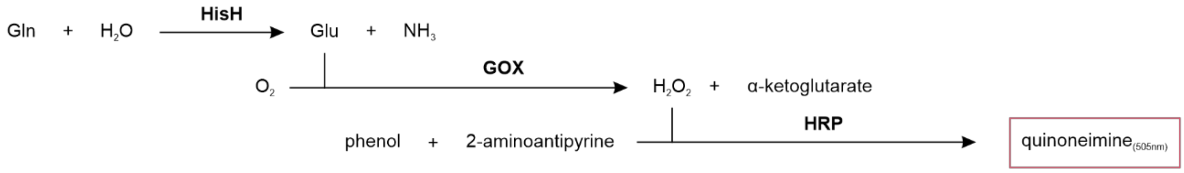

**Figure S5.** Coupled enzymatic assay for the detection of the ProFAR stimulated glutaminase activity.

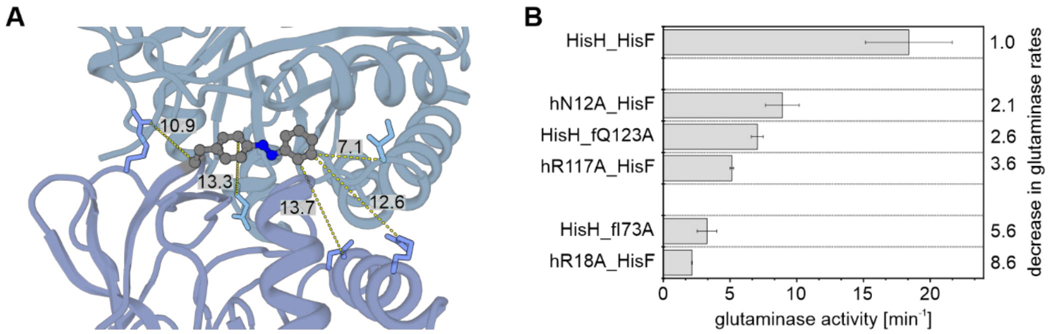

**Figure S6.** Identification of residue positions involved in allostery for randomization. A) Residues (shown as sticks) that are located at the interface and that change their sidechain conformation in a structural comparison of the inactive (PDB: 1GPW) and active HisH_HisF conformation (PDB: 7AC8). Their minimal distances to AzoF (in Å) are shown. B) Alanine scan to experimentally determine the involvement of the selected residues in allostery. The mean glutaminase activities ($v/E_0$) ± standard deviations (SD) and their decrease in glutaminase rates relative to HisH_HisF are shown.

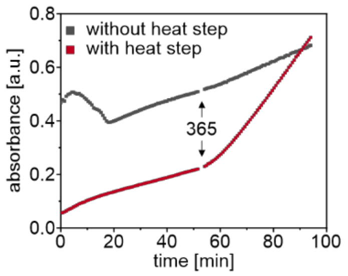

**Figure S7.** Glutaminase activity initiated by the addition of an *E. coli* lysate spiked with 0.25 µM hAzoF_HisF. The progress curves show that a heat step at 60°C for 15 min (red curve) significantly reduces background signals that are present without heat treatment (grey curve). *Irradiation:* 2 min per plate with 365 nm (screening setup, **Table S8**).

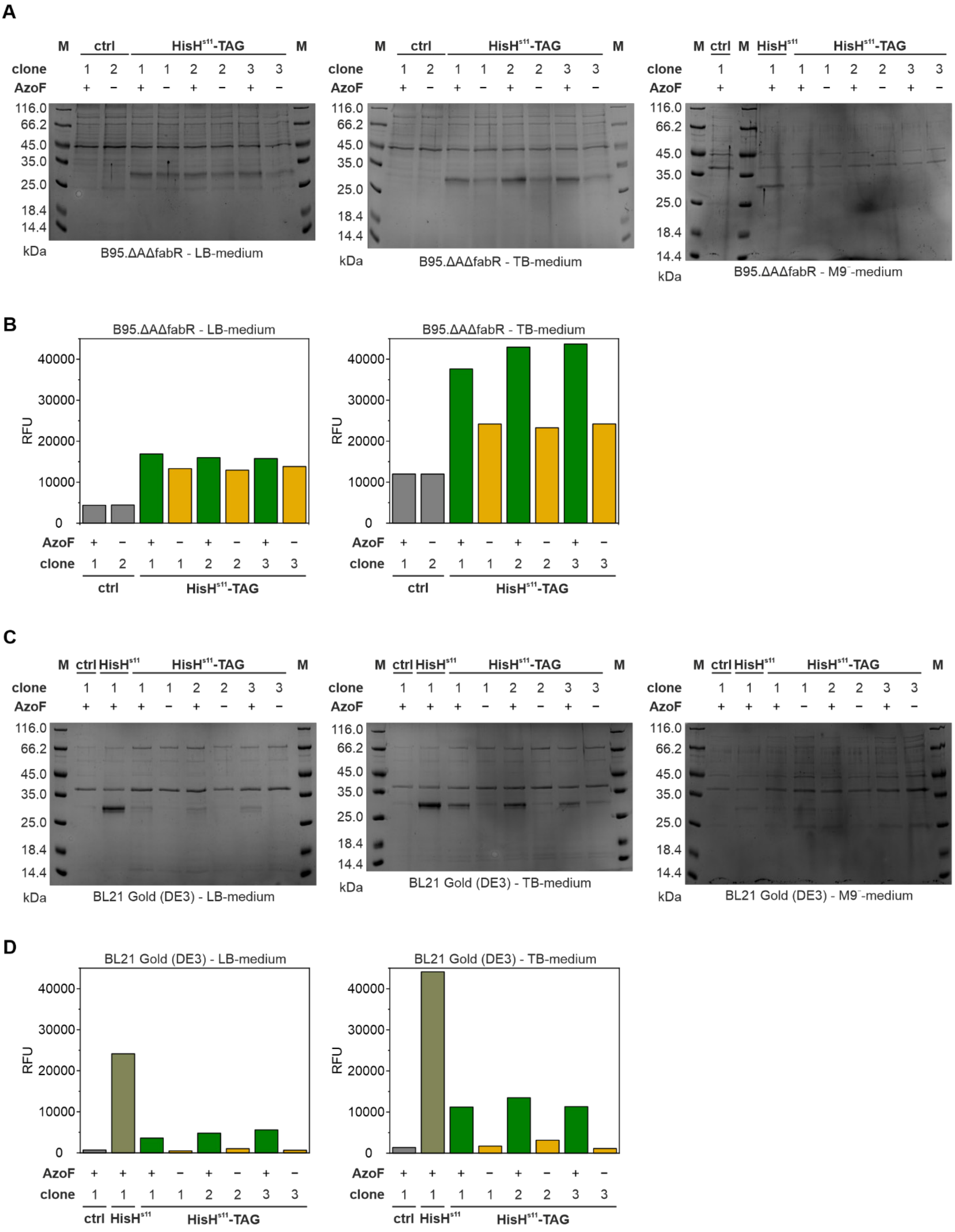

**Figure S8.** Optimization of expression conditions in microtiter scale. A) Lysates after heterologous gene expression in *E.coli*. B95.ΔAΔfabR[8] performed either in lysogeny broth (LB), terrific broth (TB), or M9⁻ medium and heat step purification (60°C, 15 min). For the control (ctrl) consisting of an empty vector two clones were tested. For the incorporation of AzoF at position W123 in HisH$^{s11}$ three clones were tested (HisH$^{s11}$-TAG), each with 0.4 mM (+) or without (−) addition of AzoF at the induction timepoint. Bands of HisH$^{s11}$ potentially containing hAzoF are visible at ~28 kDa. M: low molecular weight marker. B) Split-GFP measurements after heterologous gene expression in *E.coli*. B95.ΔAΔfabR performed in LB or TB medium. The same lysates as in A) were used, which apparently contain HisH$^{s11}$ even in the absence of AzoF indicating significant misincorporation of natural amino acids. C) Lysates after heterologous gene expression in *E.coli*. BL21 Gold (DE3) performed either in LB, TB, or M9$^{-}$ medium and heat step purification (60°C, 15 min). In addition to one control (ctrl) clone consisting of an empty vector we tested the expression of HisH$^{s11}$. For the incorporation of AzoF at position W123 in HisH$^{s11}$ three clones were tested (HisH$^{s11}$-TAG), each with 0.4 mM (+) or without (−) addition of AzoF at the induction timepoint. Bands of HisH$^{s11}$ potentially containing AzoF are visible at ~28 kDa. M: low molecular weight marker. D) Split-GFP measurements after heterologous gene expression in *E.coli*. BL21 Gold (DE3) performed in LB or TB medium. The same lysates as in C) were used, which showed only residual HisH$^{s11}$ signals in the absence of AzoF indicating only negligible misincorporation of natural amino acids. *Note:* to simplify the labeling, only the term HisH$^{s11}$ is used in the panels; the combination of the label HisH$^{s11}$-TAG and a "+" for AzoF indicates the occurrence of hAzoF$^{s11}$.

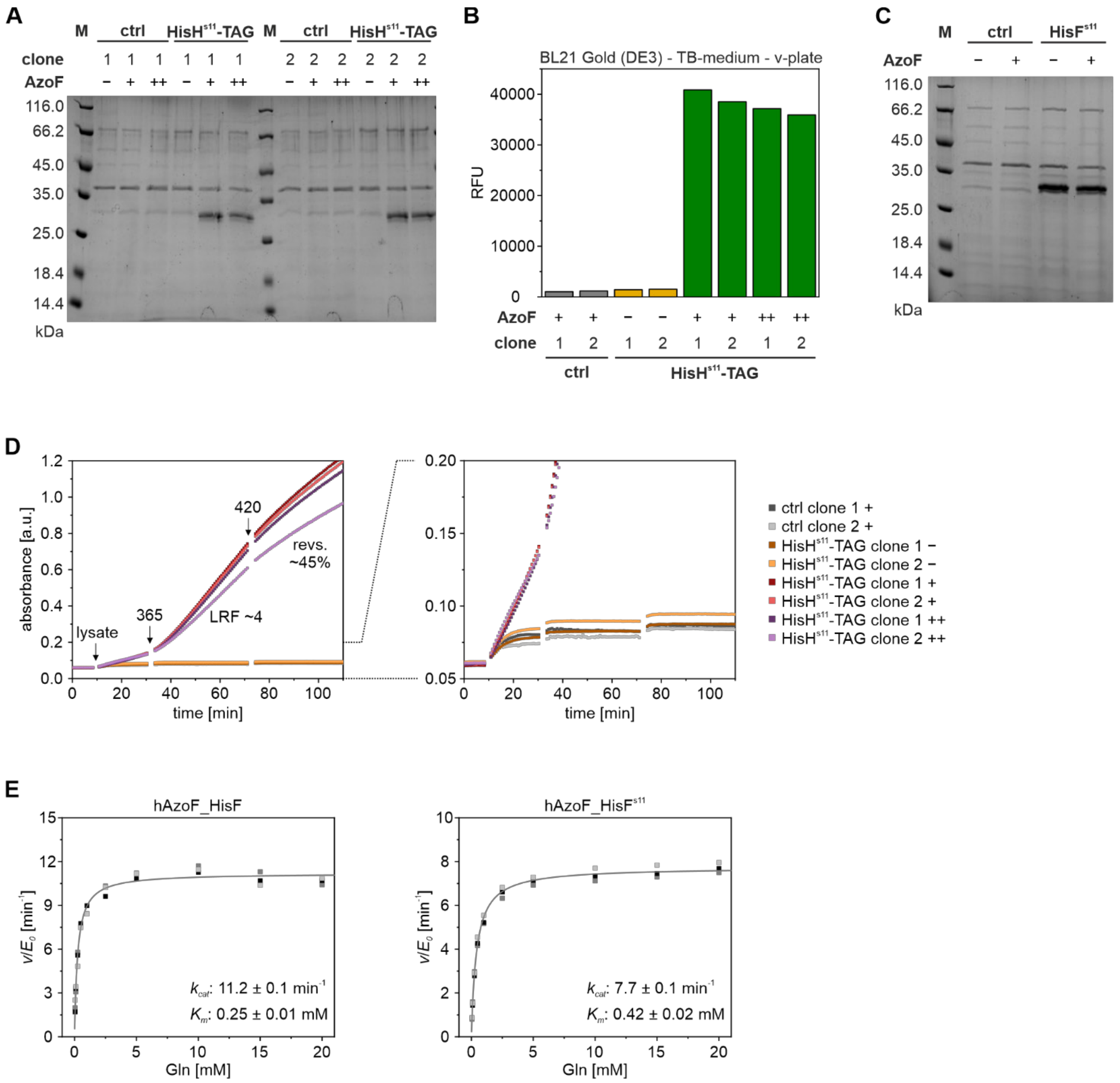

**Figure S9.** Optimized expression conditions and signal-to-noise ratios as well as steady-state kinetics of hAzoF_HisF$^{s11}$. A) Lysates after heterologous gene expression in *E.coli*. BL21 Gold (DE3) performed in TB medium and heat step purification (60°C, 15 min). Two control (ctrl) clones consisting of an empty vector and two clones of HisH$^{s11}$-TAG were used to test the incorporation of AzoF at position W123 in HisH$^{s11}$ in terms of different UAA concentrations: 0.4 mM (+), 0.8 mM (++) or no (−) AzoF was added at the induction timepoint. Bands of HisH$^{s11}$ potentially containing AzoF are visible at ~28 kDa. M: low molecular weight marker. B) Split-GFP measurements after heterologous gene expression in *E.coli*. BL21 Gold (DE3) using the same lysates as in A). C) Lysates after heterologous gene expression in *E.coli*. BL21 Gold (DE3) performed in TB medium and heat step purification (60°C, 15 min). In addition to a control (ctrl) clone consisting of an empty vector we tested the expression of HisF$^{s11}$ with 0.4 mM (+) or without (−) addition of AzoF at the induction timepoint. Bands of HisF$^{s11}$ are visible at ~33 kDa. M: low molecular weight marker. D) Progress curves of real-time photocontrol measurements performed with the same lysates as in A) indicate that optimizing expression conditions and performing a heat step facilitate the determination of LRFs and reversibility values with optimal signal-to-noise ratios. Addition of higher AzoF concentrations during expression did not affect the photocontrol efficiency. Shown is the mean LRF (~4) and reversibility value (~13%) among all four hAzoF lysates. *Irradiation:* 2 min per plate with 365 nm for PSS$^{365}$ and 2 min per plate with 420 nm for PSS$^{420}$ (screening setup, **Table S8**). E) Steady-state kinetics performed with hAzoF_HisF and hAzoF_HisF$^{s11}$ demonstrate that glutaminase activity is largely retained with the s11-tag. *Note:* to simplify the labeling, only the term HisH$^{s11}$ is used in the panels; the combination of the label HisH$^{s11}$-TAG and a "+" for AzoF indicates the occurrence of hAzoF$^{s11}$.

|  | 1 | 2 | 3 | 4 | 5 | 6 | 7 | 8 | 9 | 10 | 11 | 12 |
|---|---|---|---|---|---|---|---|---|---|---|---|---|
| A | D | S | M | G | F | N | W | P | H | I | neg ctrl |  |
| B | T | Y | C | E | K | A | R | V | pos ctrl | Q | neg ctrl |  |
| C | H | I | D | S | M | G | F | N | W | P | TB |  |
| D | pos ctrl | Q | T | Y | C | E | K | A | R | V | TB |  |
| E | W | P | H | I | D | S | M | G | F | N | TB |  |
| F | R | V | pos ctrl | Q | T | Y | C | E | K | A | TB |  |
| G | neg ctrl | N | W | P | H | I | D | S | M | G | F |  |
| H | neg ctrl | A | R | V | pos ctrl | Q | T | Y | C | E | K |  |

**Figure S10.** Exemplary representation how four biological replicates of each library member were equally distributed in a 96-well plate. A–Y: Variants containing the respective amino acid replacement at the chosen position (e.g. L2 in HisF); pos ctrl: the parental construct hAzoF_HisF; neg ctrl: empty expression vector; TB: TB medium instead of lysate.

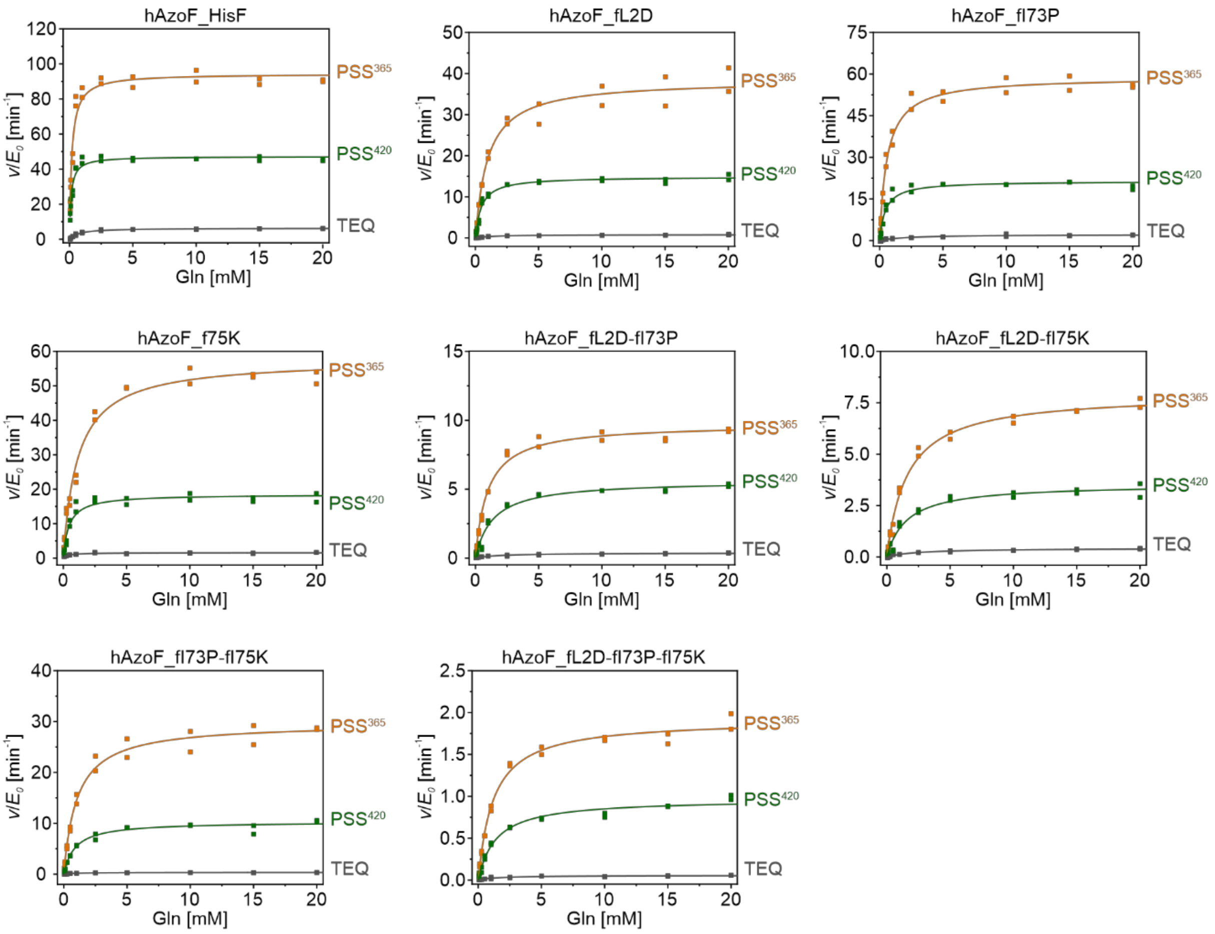

**Figure S11.** Michaelis-Menten curves for the glutaminase activity of all single, double and triple hAzoF_HisF variants. For each variant technical duplicates before (TEQ) and after irradiation ($PSS^{365}$ and $PSS^{420}$) were measured. Data of the duplicates were fitted with **Equation S5** to determine $k_{cat}$ and $K_m$ and **Equation S6** to determine $k_{cat}/K_m$ values ± standard errors (SE). *Irradiation:* 2 s per well with 365 nm for $PSS^{365}$ and 2 s per well with 365 nm followed by 4 s per well with 420 nm for $PSS^{420}$ (individual setup, **Table S8**).

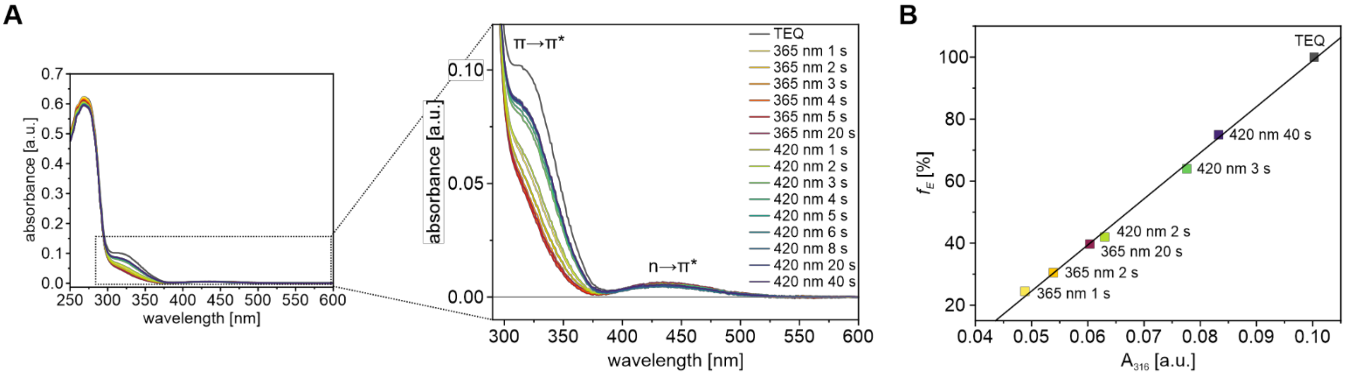

**Figure S12.** Exemplary estimation of *E*:*Z* ratios based on UV/Vis analysis. A) UV/Vis spectra of 25 µM hAzoF_HisF in 20 mM Tris-HCl (pH 7.0) in TEQ and after irradiation with 365 nm and 420 nm for various time-points (individual setup, **Table S8**). Complete spectra and views zoomed-in on the π→π* and n→π* signals of AzoF are shown. B) Obtained $f_E$ values of selected spectra (a detailed description of the analysis is given in the Experimental Section) were plotted against the measured absorbance at 316 nm and fitted with a linear regression model. The resulting linear fit equation was used to estimate $f_E$ of any spectrum within the experiment. The same procedure was performed for hAzoF_fL2D and hAzoF_fI73P-fI75K.

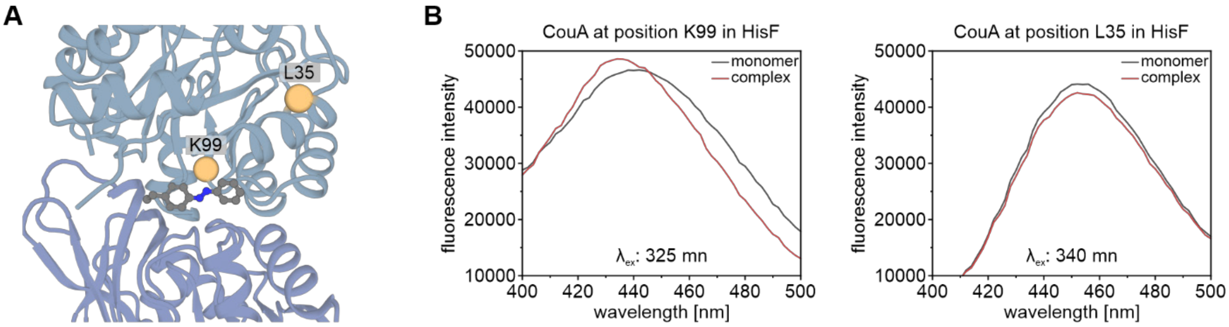

**Figure S13.** Identification of a suitable incorporation position of CouA for sensitive evaluation of heterodimer formation in hAzoF_HisF variants. A) Selection of two positions close to the interface, which tolerate UAA incorporation as previously confirmed.[14] While position L35 is located outside the interface, which could reduce the signal strength, position K99 is part of the interface, which might facilitate a good signal. Although the introduction of CouA directly in the interface might also impact the $K_d$, we anticipate that it can be used for a comparative analysis of TEQ and PSS$^{365}$ as well as different hAzoF_HisF variants. B) Emission spectra of both recombinantly produced and purified CouA-containing HisF variants (500 nM) as monomer (black) and in complex with saturated concentrations of HisH (5 µM; red). Incorporation of CouA in position K99 in HisF (HisF$^{CouA}$) led to a change in fluorescence intensity as well as a wavelength shift of the emission spectrum resulting in a higher signal change induced by complex formation compared to incorporation of CouA at position L35, which only shows a change in fluorescence intensity.

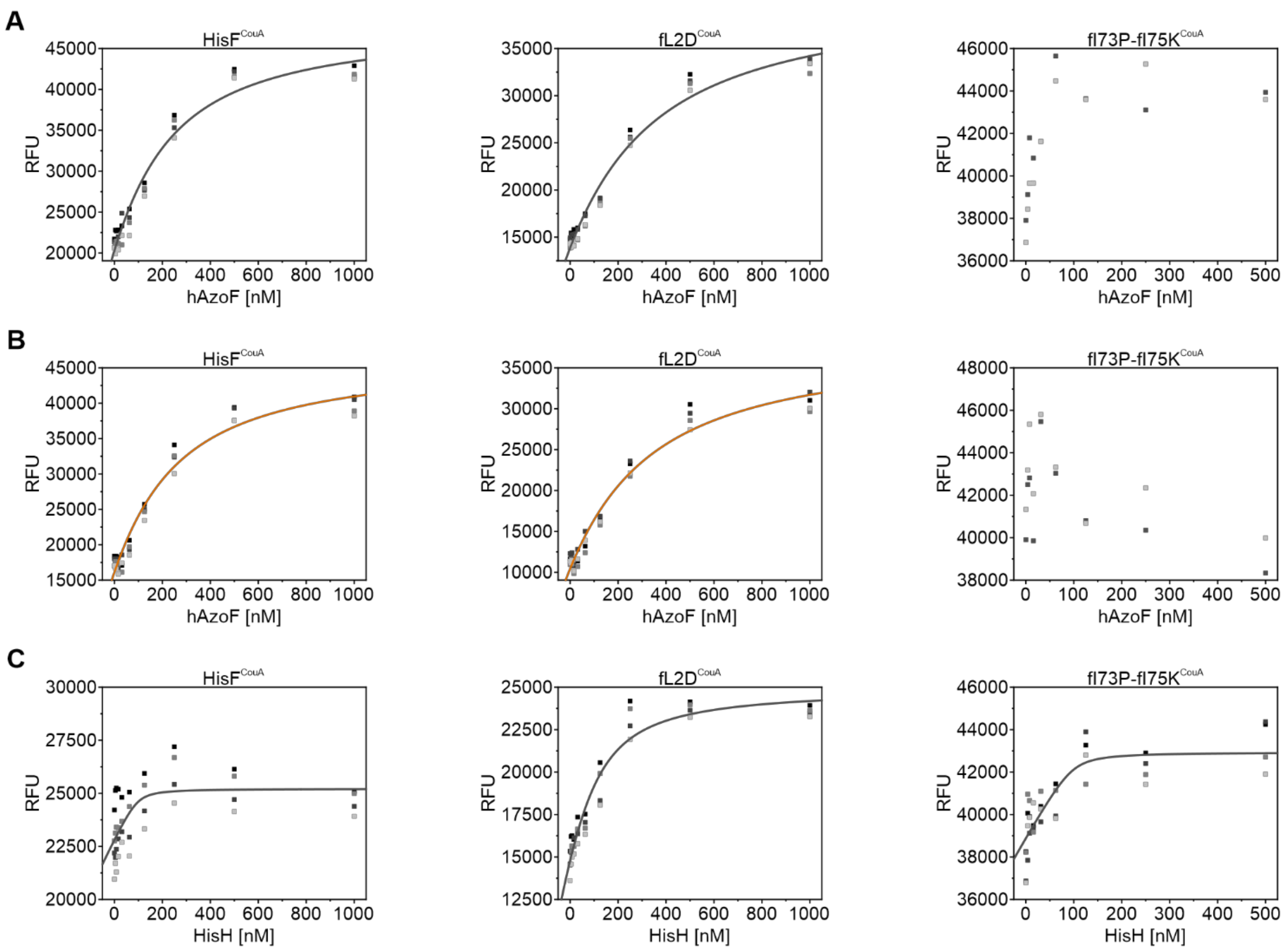

**Figure S14.** Fluorescence titration curves with HisF$^{CouA}$, fL2D$^{CouA}$ and fI73P-fI75K$^{CouA}$. A) Measurements performed with different hAzoF concentrations in TEQ. B) Measurements performed with different hAzoF concentrations in PSS$^{365}$ (*irradiation:* 2 s per well with 365 nm; individual setup, **Table S8**). C) Measurements performed with different HisH concentrations. Data were obtained from technical duplicates measured twice (different grey shades) and fitted with **Equation S12** to determine $K_D$ values ± SE.

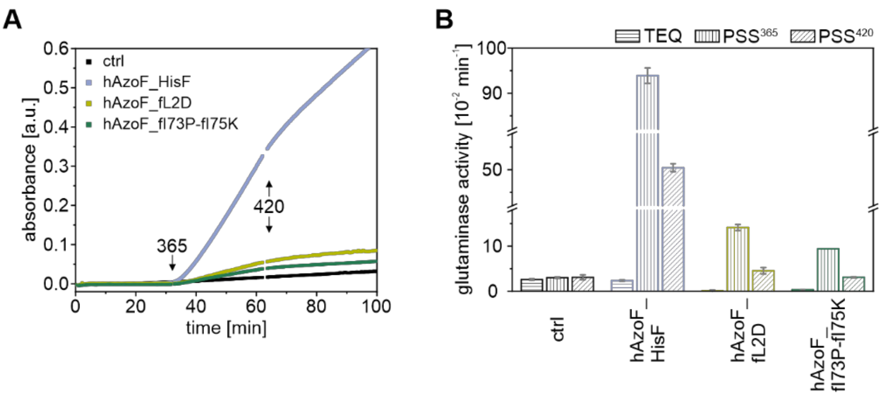

C

| | LRF(rt) | revs.(rt) [%] |
|---|---|---|
| hAzoF_HisF | 39 | 47 |
| hAzoF_fL2D | 71 | 69 |
| hAzoF_fI73P-fI75K | 24 | 70 |

**Figure S15.** Real-time photocontrol in the absence of an allosteric stimulator for hAzoF_HisF, hAzoF_fL2D, and hAzoF_fI73P-fI75K. A) Exemplary raw data from real-time photocontrol experiments of hAzoF_HisF, the two selected variants, and HisH_HisF as control (ctrl). B) Mean ± SD of technical triplicates for each single variant before (TEQ) and after irradiation (PSS$^{365}$, PSS$^{420}$). C) Summary of mean LRF(rt) and revs.(rt) values. *Irradiation:* 2 s per well with 365 nm for PSS$^{365}$ and 4 s per well with 420 nm for PSS$^{420}$ (individual setup, **Table S8**).

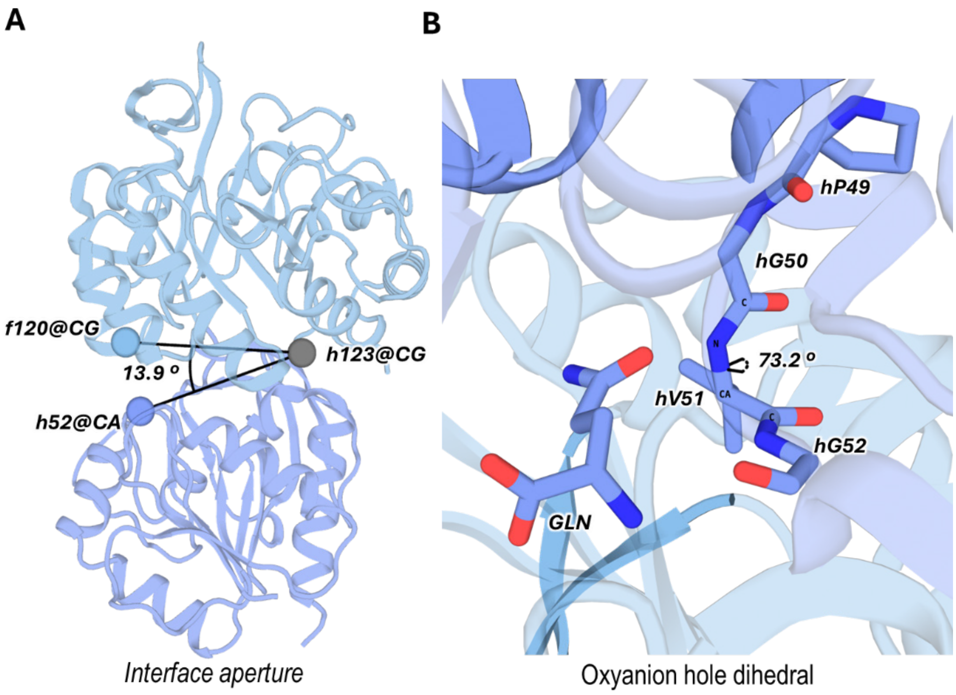

**Figure S16.** Definition of the catalytically active closed state of HisH_HisF (PDB-ID 7AC8, chains E+F). A) Definition of the subunit interface closure angle based on residues f120, h123 and h52. The crystallographic closed state displays an angle of 13.9º. B) Representation of the active state of the oxyanion strand h49PGVG52: the amide backbone of residue hV51 is pointing towards the carbonyl group of the glutamine substrate with a dihedral angle of 72.3°.

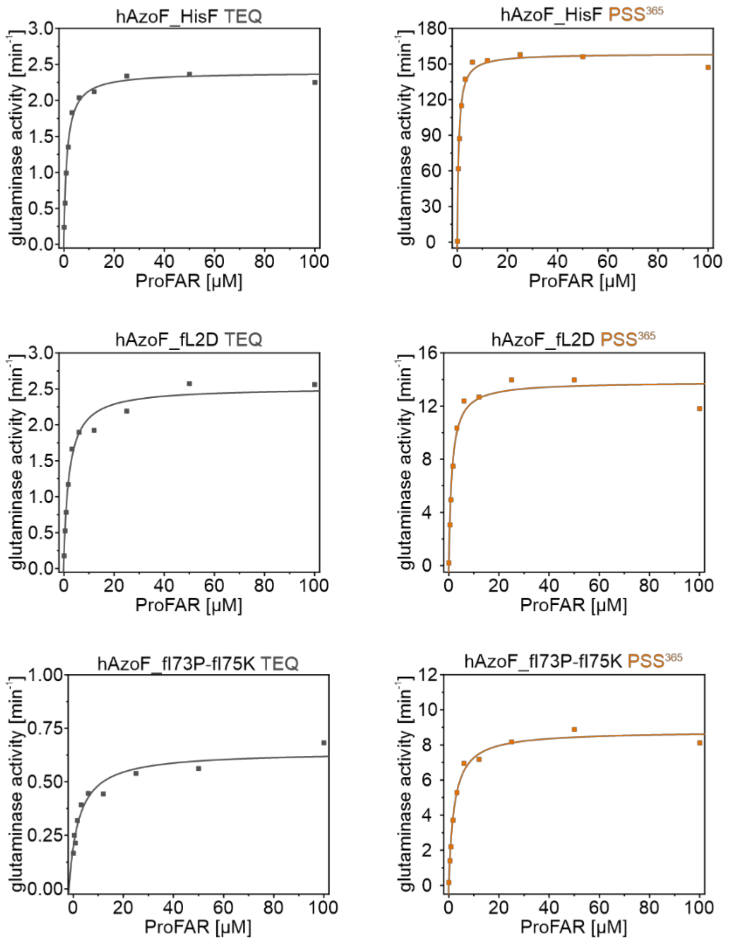

**Figure S17.** Glutaminase activities of hAzoF_HisF variants in dependence on different ProFAR concentrations. Each variant was measured in TEQ and $PSS^{365}$ (*irradiation:* 10 s per well with 365 nm; individual setup, **Table S8**). Data were fitted with **Equation S12** to determine $K_{ac}^{ProFAR}$ values ± SE.

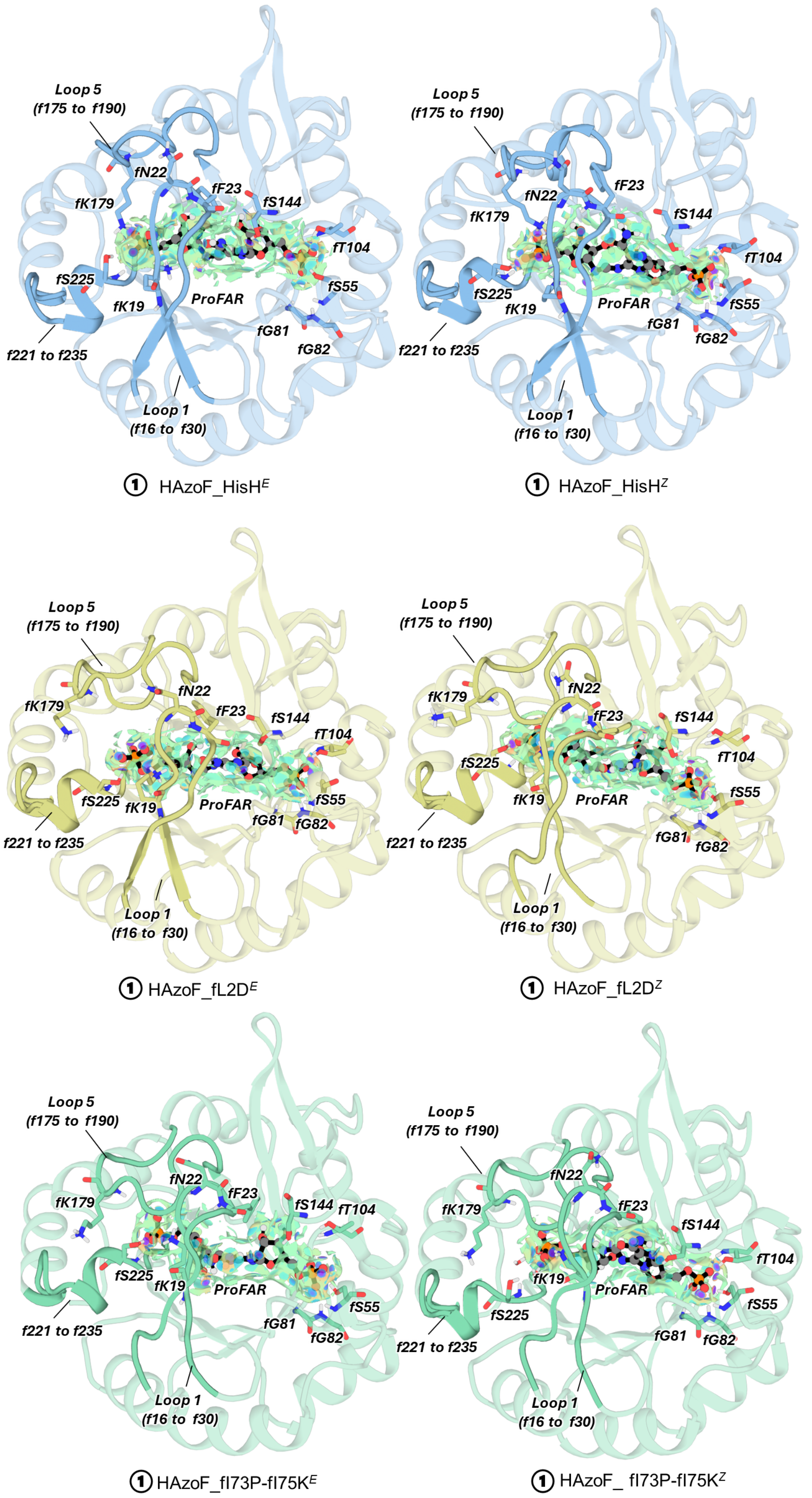

**Figure S18.** ProFAR interactions at the ProFAR binding site in HisF for the different analyzed systems: hAzoF_HisF (upper panel), hAzoF_fL2D (middle panel), and hAzoF_fI73P-fI75K (lower panel). NCI analysis suggests a higher number of stronger interactions (shown as purple-blue regions) in all hAzoF$^Z$ states. However, major differences are observed in the left phosphate group of ProFAR where the interactions for hAzoF_fL2D$^Z$ and hAzoF_fI73P-fI75K variants are weaker (green-yellow color) than for the hAzoF_HisF.

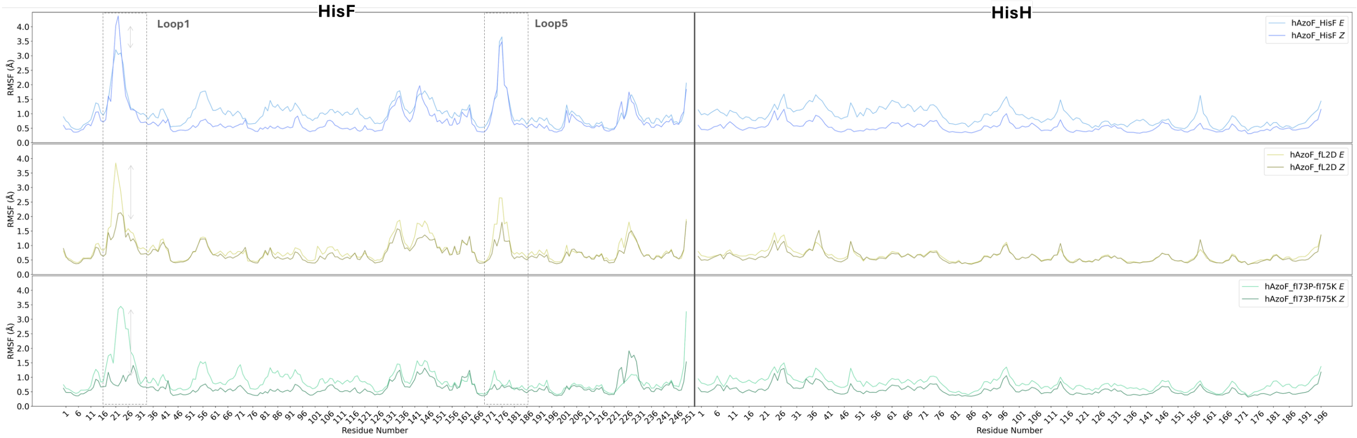

**Figure S19.**RMSF analysis of the three hAzoF_HisF variants. Major differences were found in the HisF subunit where loop1 flexibility exhibited a small shift between *E*/*Z* configurations for hAzoF_HisF, while this shift increased for hAzoF_fL2D and is even more pronounced in hAzoF_fl73P-fl75K. Regarding loop5, the flexibility has been drastically reduced for hAzoF_fl73P-fl75K variant.

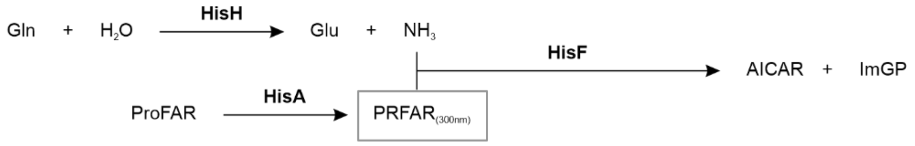

**Figure S20.** Assay for the detection of the coupled HisH_HisF activity in the presence of glutamine as a source of ammonia (supplied by HisH).

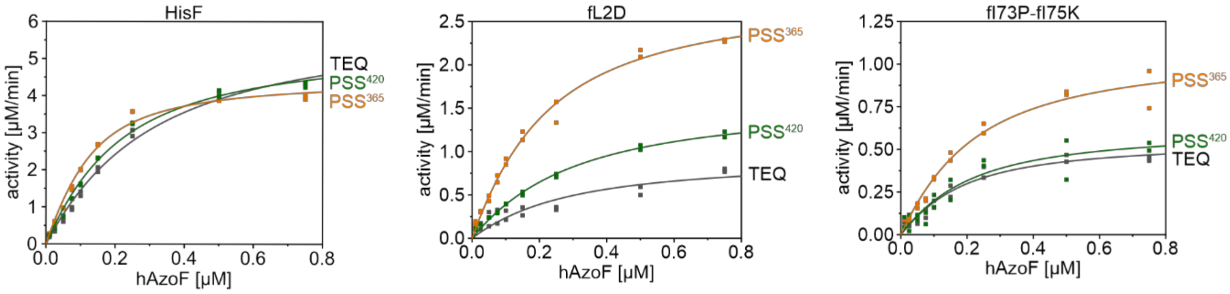

**Figure S21.** HisF activity following PRFAR turnover of HisF, fL2D, and fl73P-fl75K in dependence on different hAzoF concentrations. For each variant technical duplicates before (TEQ) and after irradiation ($PSS^{365}$ and $PSS^{420}$) were measured. Data were fitted with **Equation S12** to determine $K_{ac}^{hAzoF}$ and $k_{cat}^{app}$ values ± SE. *Irradiation:* 2 s per well with 365 nm for $PSS^{365}$, and 2 s per well with 365 nm followed by 4 s per well with 420 nm for $PSS^{420}$ (individual setup; **Table S8**).

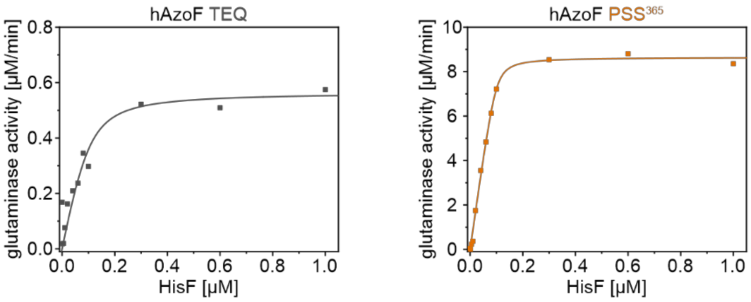

**Figure S22.** ProFAR-stimulated glutaminase activity of hAzoF in dependence on different HisF concentrations. Single measurements were performed in TEQ and PSS365. Data were fitted with **Equation S12** to determine $K_{ac}^{HisF}$ and $k_{cat}^{app}$ values ± SE. *Irradiation:* 10 s per well with 365 nm; (individual setup, **Table S8**).

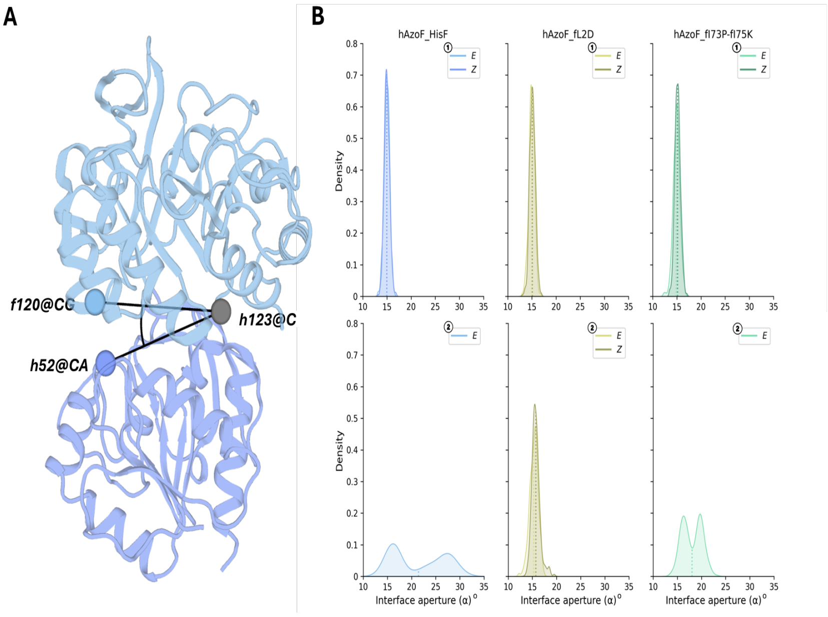

**Figure S23.** Analysis of the subunit interface closure. A) Definition of the subunit interface angle α. B) Distribution of angles α in hAzoF_HisF variants. Minimum 1 (top panels in B) for all variants shows a closed interface while for minimum 2 (bottom panel), hAzoF_HisF$^{E}$ and hAzoF_fI73P-fI75K$^{E}$ present a wider distribution of the angle α, thus indicating that open subunit interface states are visited.

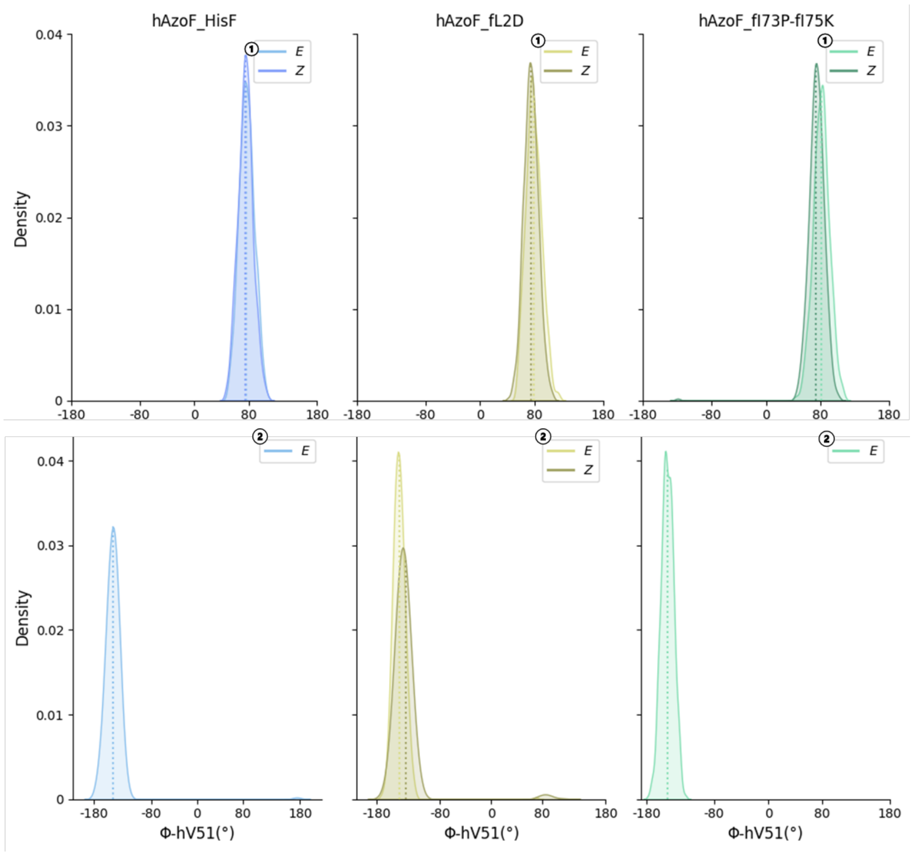

**Figure S24.** Distribution of dihedral angle φ of the oxyanion hole residue hV51 in HisH for hAzoF_HisF variants. Minimum 1 (top panel) for all variants shows an oxyanion hole formed (active form) while for minimum 2 (bottom panel) the oxyanion hole is not in its active form.

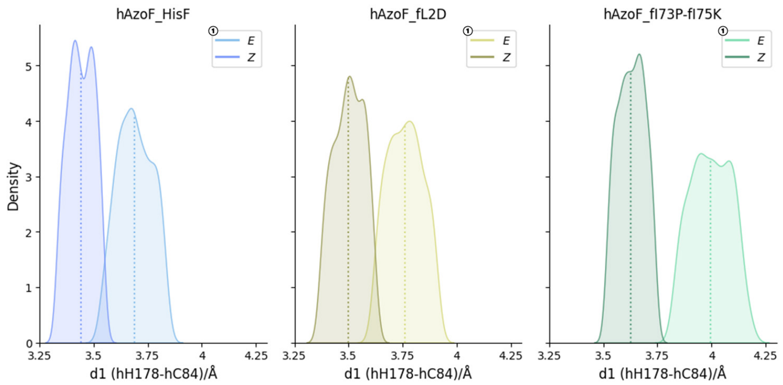

**Figure S25.** Distribution of the hydrogen bond distance between the catalytic residues hH178 and hC84 (d1) in HisH for hAzoF_HisF variants. In minimum 1, hAzoF_HisF$^{Z}$ exhibits the shortest distance followed by hAzoF_fL2D$^{Z}$ and hAzoF_fI73P-fI75K$^{Z}$ consecutively. The same trend is observed with *E* configuration but in all cases the distances are longer than those observed for the *Z* configuration.

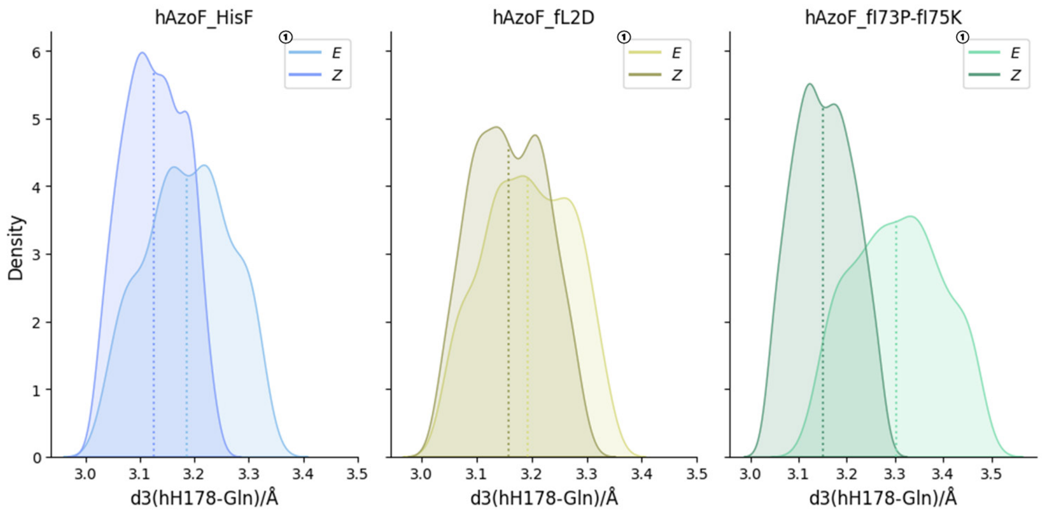

**Figure S26.** Distribution of the distance between the catalytic residue hH178 and the substrate glutamine (d3) in HisH for hAzoF_HisF variants. In minimum 1, hAzoF_HisF$^{Z}$ exhibits the shortest distance followed by hAzoF_fL2D$^{Z}$ and hAzoF_fI73P-fI75K$^{Z}$. The same trend is observed with *E* configuration but in all cases the distances are longer than those observed for the *Z* configuration.

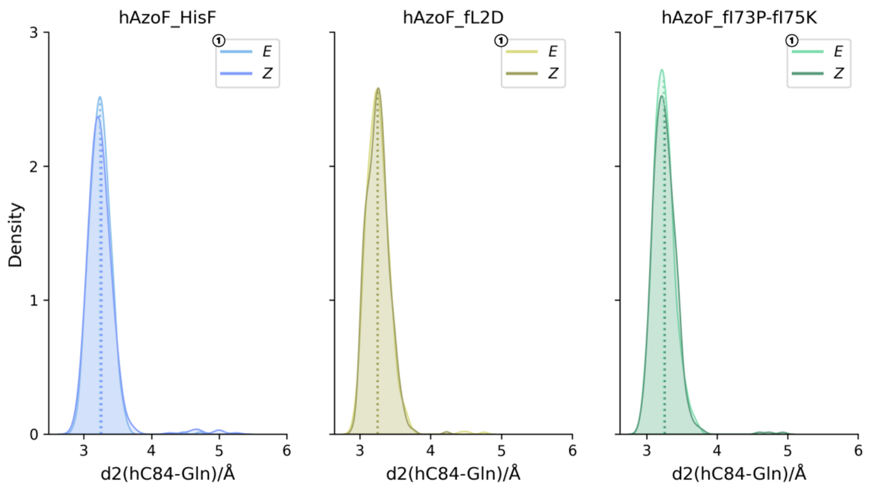

**Figure S27.**Distribution of the nucleophilic attack distance between the catalytic residue hC84 and the substrate glutamine (d2) in HisH for hAzoF_HisF variants. For minimum 1, there are no differences between *E*/*Z* conformations and variants.

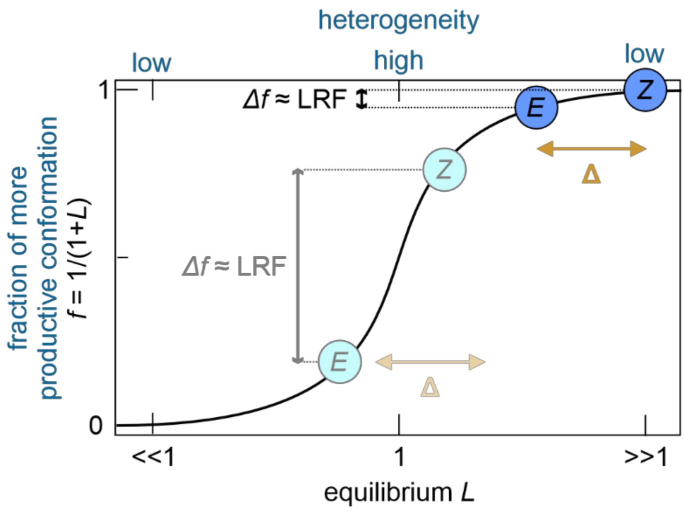

**Figure S28.** Photoreceptor model explaining how a change in conformational heterogeneity increases the LRF.[15–17] An equilibrium *L* is established between two conformations, one more productive than the other. The fraction *f* of the more productive conformation is thereby a function of *L*, and the difference in *f* (*Δf*) between the dark and lit state, corresponding to *E* and *Z* in photoxenases, influences the LRF. *E* and *Z* states adopt different equilibria *L* resulting in a conformational shift of a defined size (gold Δ). To improve the LRF, both states can be moved towards equilibria of higher heterogeneity (*L* ~1).

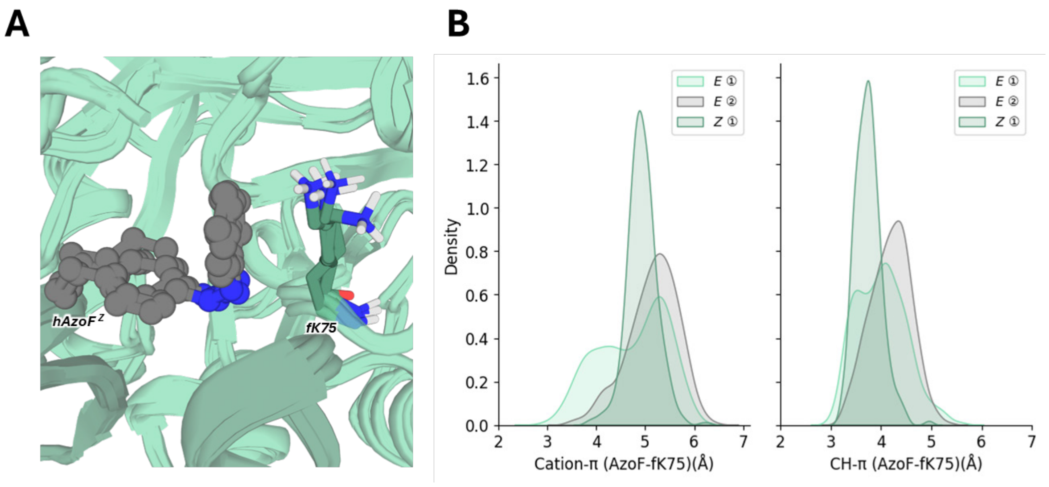

**Figure S29.** fK75 establishes a cation-π interaction with the incorporated AzoF. A) Overlay of several MD snapshots showing the cation-π interaction of AzoF and the mutation fK75 in hAzoF_fI73P-fI75K. B) Distribution of the cation-π and CH-π interaction for minima 1 and 2 when AzoF is in the *E* and *Z* configurations.

#### Supplementary Tables

**Table S1.** Glutaminase activities in the presence of ProFAR for TEQ, PSS$^{365}$, and PSS$^{420}$ as well as mean LRFs and mean reversibility (revs.) values of the parental construct and the three identified hits determined during the screening of enzyme libraries. Activities of all four replicates in the two measured plates are shown.

| | TEQ [min$^{-1}$] | PSS$^{365}$ [min$^{-1}$] | PSS$^{422}$ [min$^{-1}$] | mean LRF[a] | mean revs.[b] |
|---|---|---|---|---|---|
| **screening of position L2** | | | | | |
| **plate 1 hAzoF_HisF** | 14.0 | 44.2 | 31.8 | 3.3 | 41% |
| | 14.4 | 45.5 | 33.2 | | |
| | 12.7 | 42.0 | 30.1 | | |
| | 11.6 | 42.1 | 29.3 | | |
| **plate 2 hAzoF_HisF** | 12.9 | 39.8 | 28.6 | 3.1 | 44% |
| | 9.7 | 32.1 | 21.3 | | |
| | 15.6 | 44.0 | 32.5 | | |
| | 11.0 | 35.7 | 24.5 | | |
| **plate 1 hAzoF_fL2D** | 4.9 | 21.9 | 9.7 | 4.1 | 72% |
| | 3.4 | 14.4 | 6.9 | | |
| | 3.1 | 9.6 | 4.6 | | |
| | 3.3 | 14.4 | 6.7 | | |
| **plate 2 hAzoF_fL2D** | 2.6 | 11.1 | 4.4 | 4.4 | 75% |
| | 2.0 | 11.2 | 5.0 | | |
| | 2.2 | 9.4 | 3.9 | | |
| | 3.6 | 14.7 | 6.1 | | |
| **screening of position I73** | | | | | |
| **plate 1 hAzoF_HisF** | 3.9 | 19.6 | 17.8 | 4.9 | 11% |
| | 3.9 | 19.6 | 17.7 | | |
| | 4.2 | 18.5 | 17.1 | | |
| | 3.0 | 16.6 | 15.4 | | |
| **plate 2 hAzoF_HisF** | 2.5 | 16.7 | 15.1 | 5.8 | 12% |
| | 4.5 | 22.1 | 20.3 | | |
| | 3.8 | 20.3 | 18.1 | | |
| | 1.0 | 9.3 | 8.3 | | |
| **plate 1 hAzoF_fI73P** | 1.5 | 12.9 | 11.5 | 14.1 | 12% |
| | 0.7 | 11.5 | 9.5 | | |
| | 1.6 | 10.1 | 10.0 | | |
| | 0.7 | 9.5 | 8.4 | | |

| | | | | | |
|---|---|---|---|---|---|
| **plate 2 hAzoF_fI73P** | 1.4 | 13.0 | 11.3 | 9.5 | 17% |
| | 1.2 | 11.5 | 9.4 | | |
| | 0.7[c] | 1.3[c] | 1.1[c] | | |
| | 1.3 | 12.4 | 10.6 | | |
| **screening of position I75** | | | | | |
| **plate 1 hAzoF_HisF** | 1.1 | 15.8 | 12.0 | 10.0 | 30% |
| | 0.5 | 3.2 | 2.2 | | |
| | 0.5 | 5.1 | 3.6 | | |
| | 0.6 | 3.8 | 2.6 | | |
| **plate 2 hAzoF_HisF** | 0.8 | 4.8 | 3.3 | 9.5 | 33% |
| | 1.0 | 4.7 | 3.3 | | |
| | 0.4 | 2.3 | 1.5 | | |
| | 0.1 | 9.7 | 7.0 | | |
| **plate 1 hAzoF_fI75K** | 0.9 | 10.7 | 6.8 | 15.1 | 35% |
| | 0.7 | 9.1 | 6.0 | | |
| | 1.0 | 18.9 | 13.3 | | |
| | 0.6[c] | 0.6[c] | 0.3[c] | | |
| **plate 2 hAzoF_fI75K** | 0.1[c] | n.d.[c] | n.d.[c] | 13.9 | 35% |
| | 0.0[c] | 0.1[c] | n.d.[c] | | |
| | 0.8 | 9.5 | 6.3 | | |
| | 0.6 | 9.7 | 6.6 | | |

*Irradiation:* 2 min per plate with 365 nm for $PSS^{365}$ and 2 min per plate with 420 nm for $PSS^{420}$ (screening setup, **Table S8**). [a] Comparing the activities of TEQ and $PSS^{365}$. [b] Reversibility (revs.) values indicate the return of the activity in $PSS^{420}$ from $PSS^{365}$ to TEQ in percent. [c] Values were not included in the calculation of mean LRF and mean revs. due to their low activity and lack of response after irradiation.

**Table S2.** Glutaminase activities in the presence of ProFAR for TEQ, PSS$^{365}$, and PSS$^{420}$ as well as LRF(rt) and revs.(rt) values for hAzoF_HisF variants and wild-type HisH_HisF as determined from real-time photocontrol experiments in three technical replicates.

| variant | TEQ [min$^{-1}$] | PSS$^{365}$ [min$^{-1}$] | PSS$^{420}$ [min$^{-1}$] | LRF(rt)[a] | revs.(rt) [%][b] |
|---|---|---|---|---|---|
| **HisH_HisF** | 61.6 | 69.8 | 64.0 | | |
| | 61.0 | 69.9 | 63.5 | | |
| | 52.8 | 60.3 | 56.5 | | |
| **hAzoF_HisF** | 11.9 | 177.3 | 108.5 | 14.9 | 41.6 |
| | 14.8 | 200.2 | 130.7 | 13.5 | 37.5 |
| | 9.0 | 153.7 | 87.4 | 17.1 | 45.8 |
| **hAzoF_fL2D** | 0.8 | 61.6 | 21.4 | 77.0 | 66.1 |
| | 0.4 | 59.8 | 21.2 | 149.5 | 65.0 |
| | 0.8 | 52.6 | 18.0 | 65.8 | 66.8 |
| **hAzoF_fI73P** | 3.5 | 159.2 | 68.8 | 45.5 | 58.1 |
| | 4.2 | 179.2 | 85.9 | 42.7 | 53.3 |
| | 2.9 | 141.8 | 55.4 | 48.9 | 62.2 |
| **hAzoF_fI75K** | 2.0 | 149.1 | 67.9 | 74.6 | 55.2 |
| | 2.9 | 157.0 | 73.5 | 54.1 | 54.2 |
| | 1.9 | 140.0 | 61.6 | 73.7 | 56.8 |
| **hAzoF_fL2D-fI73P** | 0.5 | 16.9 | 3.9 | 33.8 | 79.3 |
| | 0.3 | 13.1 | 3.0 | 43.7 | 78.9 |
| | 0.4 | 14.7 | 3.7 | 36.8 | 76.9 |
| **hAzoF_fL2D-fI75K** | 0.3 | 10.0 | 2.7 | 33.3 | 75.3 |
| | 0.1 | 6.8 | 1.7 | 68.0 | 76.1 |
| | 0.5 | 7.7 | 2.1 | 15.4 | 77.8 |
| **hAzoF_fI73P-fI75K** | 0.4 | 34.9 | 8.6 | 87.3 | 76.2 |
| | 0.4 | 28.4 | 7.0 | 71.0 | 76.4 |
| | 0.7 | 34.1 | 9.2 | 48.7 | 74.6 |
| **hAzoF_fL2D-fI73P-fI75K** | 0.1 | 2.1 | 0.2 | 21.0 | 95.0 |
| | n.d. | 1.9 | 0.3 | n.d. | n.d. |
| | 0.1 | 1.8 | 0.4 | 18.0 | 82.4 |

*Note:* Exemplary raw data and bar graph of mean ± SE are shown in **Figure 4**. *Irradiation:* 2 s per well with 365 nm for PSS$^{365}$ and 4 s per well with 420 nm for PSS$^{420}$ (individual setup, **Table S8**). [a] Comparing the activities of TEQ and PSS$^{365}$. [b] Reversibility [revs.(rt)] values indicate the return of the activity in PSS$^{420}$ from PSS$^{365}$ to TEQ in percent.

**Table S3.** Glutaminase steady-state kinetics for hAzoF combined with different HisF variants.

| HisF variant | state | $K_m$ [mM] | LRF($K_m$)[a] | LRF($K_m$)/ LRF($K_m$)$_{HisF}$[b] | $k_{cat}/K_m$ [$s^{-1}$ $M^{-1}$] | LRF($k_{cat}/K_m$)[c] | LRF($k_{cat}/K_m$)/ LRF($k_{cat}/K_m$)$_{HisF}$[d] |
|---|---|---|---|---|---|---|---|
| **HisF** | TEQ | 0.71 ± 0.06 | 3.9 | 1.0 | 149.3 ± 11.5 | 57 | 1.0 |
| | PSS$^{365}$ | 0.18 ± 0.02 | | | 8544.1 ± 857.3 | | |
| | PSS$^{420}$ | 0.13 ± 0.01 | | | 5958.2 ± 609.2 | | |
| **fL2D** | TEQ | 1.25 ± 0.15 | 1.3 | 0.3 | 11.2 ± 1.2[c] | 60 | 1.0 |
| | PSS$^{365}$ | 0.96 ± 0.11 | | | 669.4 ± 68.7[c] | | |
| | PSS$^{420}$ | 0.45 ± 0.05 | | | 547.9 ± 50.5 | | |
| **fI73P** | TEQ | 2.21 ± 0.59 | 3.7 | 0.9 | 16.5 ± 3.4 | 101 | 1.8 |
| | PSS$^{365}$ | 0.59 ± 0.05 | | | 1668.8 ± 121.1 | | |
| | PSS$^{420}$ | 0.45 ± 0.06 | | | 786.4 ± 91.5 | | |
| **fI75K** | TEQ | 0.58 ± 0.13 | 0.5 | 0.1 | 28.9 ± 5.6[c] | 24 | 0.4 |
| | PSS$^{365}$ | 1.16 ± 0.11 | | | 690.0 ± 52.8 | | |
| | PSS$^{420}$ | 0.47 ± 0.08 | | | 534.9 ± 75.4 | | |
| **fL2D-fI73P** | TEQ | 2.04 ± 0.43 | 2.1 | 0.5 | 3.0 ± 0.5[c] | 55 | 1.0 |
| | PSS$^{365}$ | 0.98 ± 0.08 | | | 165.9 ± 11.5[c] | | |
| | PSS$^{420}$ | 1.39 ± 0.19 | | | 67.3 ± 7.1 | | |
| **fL2D-fI75K** | TEQ | 2.44 ± 0.42 | 1.5 | 0.4 | 2.9 ± 0.4 | 28 | 0.5 |
| | PSS$^{365}$ | 1.62 ± 0.13 | | | 81.7 ± 5.3 | | |
| | PSS$^{420}$ | 1.57 ± 0.25 | | | 37.7 ± 4.8 | | |
| **fI73P-fI75K** | TEQ | 0.88 ± 0.13 | 0.8 | 0.2 | 6.6 ± 0.9[c] | 71 | 1.2 |
| | PSS$^{365}$ | 1.06 ± 0.10 | | | 467.8 ± 37.3 | | |
| | PSS$^{420}$ | 0.89 ± 0.10 | | | 193.3 ± 17.8 | | |
| **fL2D-fI73P-fI75K** | TEQ | 1.74 ± 0.47 | 1.5 | 0.4 | 0.5 ± 0.1[c] | 54 | 0.9 |
| | PSS$^{365}$ | 1.19 ± 0.09 | | | 26.8 ± 1.7 | | |
| | PSS$^{420}$ | 1.39 ± 0.15 | | | 11.6 ± 1.0 | | |

*Note:* Michaelis-Menten curves are shown in **Figure S11**. *Irradiation:* 2 s per well with 365 nm for PSS$^{365}$ and 2 s per well with 365 nm followed by 4 s per well with 420 nm for PSS$^{420}$ (individual setup, **Table S8**). *Statistics:* $k_{cat}/K_m$ and $K_m$ values are given as fitting value ± SE for two technical replicates. [a] Ratio of $K_m$(PSS$^{365}$) and $K_m$(TEQ). [b] Ratio of LRFs($K_m$) for each variant and LRF($K_m$) of hAzoF_HisF. [c] Ratio of $k_{cat}/K_m$(PSS$^{365}$) and $k_{cat}/K_m$(TEQ). [d] Ratio of LRFs($k_{cat}/K_m$) for each variant and LRF($k_{cat}/K_m$) of hAzoF_HisF.

**Table S4.** Glutaminase activities in the presence of ProFAR for hAzoF_HisF, hAzoF_fL2D, hAzoF_fI73P-fI75K, and HisH_HisF as control (ctrl) as determined from the cycle performance of real-time photocontrol experiments (**Figure 5A,B**). Mean activity of technical duplicates for complexes with hAzoF and activity of single measurements for HisH_HisF (ctrl).

| **cycle No.** | | **0 (TEQ)** | **0.5 ($PSS^{365}$)** | **1 ($PSS^{420}$)** | **1.5 ($PSS^{365}$)** | **2 ($PSS^{420}$)** | **2.5 ($PSS^{365}$)** | **3 ($PSS^{420}$)** |
|---|---|---|---|---|---|---|---|---|
| **hAzoF_HisF** | activity “irradiated” [$min^{-1}$] | 3.6 | 87.2 | 40.9 | 87.2 | 35.6 | 81.1 | 28.5 |
| | activity “dark” [$min^{-1}$] | 3.6 | 3.8 | 4.5 | 5.0 | 4.0 | 6.0 | 5.7 |
| **hAzoF_fL2D** | activity “irradiated” [$min^{-1}$] | 0.7 | 20.5 | 6.1 | 21.9 | 4.3 | 21.3 | 3.1 |
| | acitivity “dark” [$min^{-1}$] | 0.5 | 0.5 | 0.4 | 1.1 | 0.3 | 1.9 | 1.0 |
| **hAzoF_fLI73P-fI75K** | activity “irradiated” [$min^{-1}$] | 0.1 | 11.4 | 2.6 | 11.5 | 1.6 | 10.9 | 0.9 |
| | acitivity “dark” [$min^{-1}$] | 0.7 | 0.1 | 0.3 | 0.2 | 0.0 | 0.7 | 0.6 |
| **ctrl** | activity “dark” [$min^{-1}$] | 19.6 | 25.2 | 23.1 | 27.0 | 24.1 | 27.9 | 23.1 |

*Note:* Exemplary raw data and graphical representation are shown in **Figure 5**. *Irradiation:* 2 s per well with 365 nm for $PSS^{365}$ and 4 s per well with 420 nm for $PSS^{420}$ (individual setup, **Table S8**).

**Table S5.** Glutaminase activities in the absence of ProFAR as allosteric stimulator for hAzoF_HisF variants and HisH_HisF as determined from real-time photocontrol experiments in three technical replicates.

| **variant** | **TEQ[$10^{-2}$ $min^{-1}$]** | **$PSS^{365}$ [$10^{-2}$ $min^{-1}$]** | **$PSS^{420}$ [$10^{-2}$ $min^{-1}$]** |
|---|---|---|---|
| HisH_HisF | 2.7 | 3.1 | 3.5 |
| | 2.5 | 2.9 | 2.7 |
| hAzoF_HisF | 2.3 | 95.1 | 51.1 |
| | 2.5 | 92.7 | 49.8 |
| hAzoF_fL2D | 0.02 | 13.7 | 4.0 |
| | 0.2 | 14.6 | 5.0 |
| hAzoF_fI73P-fI75K | 0.4 | 9.4 | 3.2 |
| | 0.3 | 9.4 | 3.0 |

*Note:* Exemplary raw data, bar graph of mean ± SE, LRF(rt), and reversibility [revs.(rt)] values are shown in **Figure S15**. *Irradiation:* 2 s per well with 365 nm for $PSS^{365}$ and 4 s per well with 420 nm for $PSS^{420}$ (individual setup, **Table S8**).

**Table S6.** hAzoF titration experiments resolving the effects of *E*→*Z* isomerization on PRFAR turnover at the HisF active site.

| variant | state | $k_{cat}^{app}$ [min$^{-1}$][a] | $K_{ac}^{hAzoF}$ [µM][b] |
|---|---|---|---|
| **hAzoF_HisF** | TEQ | 60.8 ± 2.8 | 0.24 ± 0.03 |
| | PSS$^{365}$ | 44.9 ± 1.3 | 0.07 ± 0.01 |
| | PSS$^{420}$ | 54.1 ± 1.8 | 0.15 ± 0.02 |
| **hAzoF_fL2D** | TEQ | 9.3 ± 1.6 | 0.22 ± 0.10 |
| | PSS$^{365}$ | 28.6 ± 1.1 | 0.16 ± 0.02 |
| | PSS$^{420}$ | 16.2 ± 0.6 | 0.24 ± 0.02 |
| **hAzoF_fI73P-fI75K** | TEQ | 5.6 ± 0.4 | 0.13 ± 0.03 |
| | PSS$^{365}$ | 10.9 ± 0.7 | 0.16 ± 0.03 |
| | PSS$^{420}$ | 6.3 ± 0.9 | 0.15 ± 0.06 |

*Note:* Raw data plots and non-linear curve fitting are shown in **Figure S21**. *Irradiation:* 2 s per well with 365 nm for PSS$^{365}$ and 2 s per well with 365 nm followed by 4 s per well with 420 nm for PSS$^{420}$ (individual setup; **Table S8**). *Statistics:* $k_{cat}^{app}$ and $K_{ac}^{hAzoF}$ values are given as fitting value ± SE for technical duplicates. [a] Apparent maximal turnover rate of glutamine-dependent HisF activity. [b] Constant determining the concentration of HisH, at which half-maximal activation of HisF is achieved.

**Table S7.** Key reagents and resources.

| Reagent or resource | Source | Identifier |
|---|---|---|
| **Chemicals, cells, and recombinant proteins** | | |
| AzoF | This paper | N/A |
| AapF | ChiroBlock | N/A |
| CouA | Sigma Aldrich | #792551 |
| ProFAR | This paper | N/A |
| HisG/IE | This paper | UniProt IDs: Q9X0D2 and Q9X0C5 |
| HisA | This paper | UniProt ID: Q9X0C7 |
| GOX | Sigma Aldrich | #G1924-5UN |
| HRP | Sigma Aldrich | #P8250-50KU |
| *E. coli* BL21 Gold (DE3) | Agilent Technologies | #230132 |
| *E. coli* NEB Turbo | New England Biolabs | #C2984H |
| *E. coli* B95.ΔAΔfabR | RIKEN BRC[8] | #RDB13712 |
| HisF variants | This paper | UniProt ID: Q9X0C6 |
| HisH variants | This paper | UniProt ID: Q9X0C8 |
| GFP1-10 | This paper | N/A |
| **Recombinant DNA** | | |
| pET28a_HisF | This paper | N/A |
| pET28a_HisF$^{s11}$ | This paper | N/A |
| pET28a_HisH | ref[12] | N/A |
| pET28a_HisH-W123TAG | ref[12] | N/A |
| PET28a_HisH$^{s11}$ | This paper | N/A |
| pET21a_GFP1-10 | This paper | N/A |
| pEVOL_AzoF-RS | Peter Schultz (Scripps Research Institute, La Jolla, CA)[18] | N/A |
| pEVOL_AapF-RS | This Paper | N/A |
| pEVOL_CouA-RS | ref[11] | N/A |
| **Software and Algorithms** | | |
| Origin 2024 | OriginLab | https://www.originlab.com |
| Pymol 3.1 | Schrödinger 2015 | https://www.pymol.org |
| Amber22 | Amber | https://ambermd.org |

**Table S8.** Specifications of LEDs used for irradiation.

| LEDs | $\lambda_{max}$ [nm] | $I_{max}$ [mA][a] | $P_{max}$ [mW] | $P_{max}$ [mW cm$^{-2}$] |
|---|---|---|---|---|
| **Screening setup** | | | | |
| 45 × Nichia NCSU275 | 365 | 500 | 150[b] | 19.4–24.9[b] |
| 45 × Avonec 1W | 420 | 350 | 500 [b] | 49.6–63.8[b] |
| **Individual setup** | | | | |
| LED ENGIN LZ4-44UV00-0000 | 365 | 850 | 2100[c] | 740[c] |
| Luxeon SZ-01-S2LHUV-0395-A065[d] | 400 | 1000 | 150[c] | 55[c] |
| Luxeon SZ-01-S4[d] | 415 | 500 | 210[c] | 75[c] |
| Avonec 1W410420m | 420 | 350 | 220[c] | 80[c] |
| Lumixtar WL-P5EP4545YG140-540 | 540 | 700 | 250[c] | 90[c] |

[a] Taken from the manufacturer specifications. [b] Determined with a RM-12 radiometer and UVA+ sensor (Optysec; range: 0–2000 mW cm$^{-2}$). [c]Determined with a Field Max$_{II}$-TO high sensitivity power sensor (Coherent, #1098579, Range 0–10 W) directly in front of the LED.

**Table S9.** Glutaminase activities in the presence of ProFAR, LRF(rt), and reversibility [revs.(rt)] values for hAzoF_HisF variants and HisH_HisF without a His$_6$-tag on HisF as determined from real-time photocontrol experiments in three technical replicates.

| variant | TEQ [min$^{-1}$] | PSS$^{365}$ [min$^{-1}$] | PSS$^{420}$ [min$^{-1}$] | LRF(rt)[a] | revs.(rt) [%][b] |
|---|---|---|---|---|---|
| HisH_HisF | 15.5 | 20.5 | 19.9 | 1.3 | |
| | 20.2 | 25.2 | 24.1 | 1.2 | |
| | 25.0 | 30.5 | 29.0 | 1.2 | |
| hAzoF_HisF | 34.0 | 266.9 | 233.6 | 7.8 | 14.3 |
| | 35.1 | 274.7 | 244.6 | 7.8 | 12.5 |
| | 34.6 | 326.4 | 283.5 | 9.4 | 14.7 |
| hAzoF_fL2D | 1.5 | 89.2 | 43.2 | 60.6 | 52.4 |
| | 1.3 | 91.2 | 43.2 | 67.8 | 53.4 |
| | 1.8 | 92.3 | 41.4 | 52.5 | 56.3 |
| hAzoF_fI73P-fI75K | 1.2 | 39.5 | 10.2 | 31.6 | 76.7 |
| | 0.4 | 36.9 | 8.7 | 92.3 | 77.2 |
| | 0.6 | 38.0 | 9.4 | 68.2 | 76.5 |

*Irradiation:* 2 s per well with 365 nm for PSS$^{365}$ and 4 s per well with 420 nm for PSS$^{420}$ (individual setup, **Table S8**). [a] Comparing the activities of TEQ and PSS$^{365}$. [b] Reversibility [revs(rt)] values indicate the return of the activity in PSS$^{420}$ from PSS$^{365}$ to TEQ in percent.

#### Experimental section

##### Chemicals and other resources

All reagents and solvents were purchased in analytical grade or higher from commercial sources and were used without further purification. *E. coli* BL21 Gold (DE3) and *E. coli* NEB Turbo were further maintained as chemically competent cells by following the manufacturer's guidelines. Key reagents and resources are listed in **Table S7**.

##### Irradiation devices

Irradiation was performed with two different setups, one for the screening assay ("screening setup") and one for all other experiments ("Individual setup"). Both setups were developed in-house. The screening setup consisted of a discarded gel documentation chamber that was equipped with exchangeable LED modules, which contained either 45 LEDs of 365 nm or 45 LEDs of 420 nm (**Table S8**). The LEDs are placed in a series connection, so that every single LED irradiates with the same intensity. The integrated power supply is run by constant electricity, meaning the voltage is automatically set to the maximum possible LED module voltage. The Ampere settings are changed depending on the LEDs used (compare **Table S8**). Furthermore, a protection circuit is integrated, which protects from excess voltage of >200 V, and automatically shuts down the LEDs when reaching a temperature of 50°C, turning them on again at approximately 40°C after cooling using an installed fan. As the protection circuit itself is supplied by electricity, it needs a permanent supply of 10 mA, which has to be added to the Ampere settings of the LED module. An integrated Reed-switch ensures that the LED module can only be turned on if the module is placed on top of the illumination chamber. Moreover, the LEDs are protected by a 3 mm thick UV-transparent acryl plate cover. The chamber itself contains a safety switch that turns out the LEDs when opening the chamber. Samples can be positioned at three different levels of varying distances to the lamp using rails installed in the chamber. For the screening assay, the closest level was used so that the plate was positioned 4 cm from the LED module. The irradiation intensities in mW or mW $cm^{-2}$ for this level were measured using different light sensors that were positioned on a tray at the respective distance (**Table S8**). Owing to the overlap of light rays originating from each single LED, the intensities vary throughout the plate (indicated by the range of values). The individual setup used to irradiate all other reactions, contained either a 365 nm, 400 nm, 415 nm, 420 nm or 540 nm LED attached to a power supply. For monochromatic irradiation, a microtiter plate with the respective samples was placed directly under the respective LED and each well was irradiated separately. The specifications of each LED are listed in **Table S8**.

##### Synthesis of AzoF

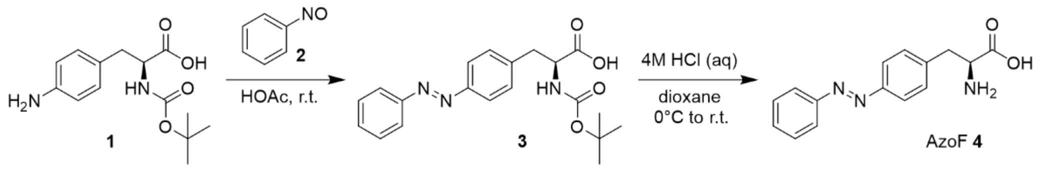

For the synthesis of AzoF, a previously reported protocol[18] was adapted.[19] In the first step, 4-amino-*N*-Boc-L-phenylalanine **1** (2.5 g, 8.95 mmol, 1.0 eq) was dissolved in glacial acetic acid (50 mL). The reaction flask was covered with aluminum foil, and nitrosobenzene **2** (955 mg, 8.93 mmol, 1.0 eq) was added. The reaction mixture was stirred at room temperature for 24 h before being quenched by pouring it onto crushed ice/water (500 mL). The product was extracted three times with ethyl acetate (3 × 100 mL). The combined organic layers were washed with water (2 × 50 mL) and brine (1 × 50 mL), followed by drying over $Na_2SO_4$. After removing the volatiles under reduced pressure, the crude product was purified using automated flash column chromatography (Biotage® Selekt, Biotage) in DCM with a linear methanol gradient (0–10%), yielding intermediate **3**, which was used directly in the next step. To obtain AzoF (**4**), intermediate **3** (1.8 g, 6.44 mmol) was dissolved in dioxane (20 mL) at 0°C. Then, 4 M HCl (6.6 mL) was added, and the reaction mixture was stirred at room temperature for 24 h. To facilitate the removal of 1,4-dioxane, toluene (50 mL) was added, and the solvent was evaporated under reduced pressure. The crude product was triturated four times with diethyl ether (4 × 50 mL) and subsequently lyophilized overnight, affording AzoF with an overall yield of 54–63%. The identity and purity of the final product were confirmed by $^1$H-NMR analysis. $^1$H-NMR (300 MHz, chloroform-*d*): δ = 7.97–7.83 (m, 4H), 7.72–7.55 (m, 3H), 7.54–7.43 (m, 2H), 4.22 (t, J = 6.4 Hz, 1H), 3.25 (t, J = 6.8 Hz, 2H).

#### Biosynthesis of ProFAR

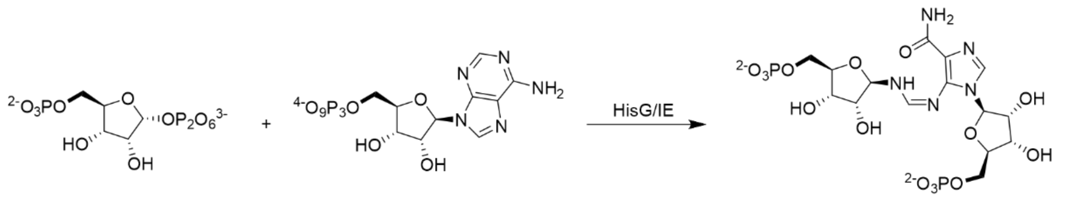

5 6 ProFAR **7**

ProFAR was synthesized following a standard protocol developed by Davisson et al.[20] from 5-phospho-D-ribosyl α-1-pyrophosphate **5** and adenosine triphosphate **6** in 50 mM $NH_4COOCH_3$ (pH 7.8) using the enzyme HisG/IE. The success of the reaction was tracked spectrophotometrically and after total conversion the products were purified with ion-exchange chromatography using a POROS column (HQ 20, 10 mL, Applied Biosystems) and a linear gradient of $NH_4COOCH_3$ (50 mM–1 M). Concentration of ProFAR **7** was determined at 300 nm ($\varepsilon_{300}$ = 6069 $M^{-1}cm^{-1}$)[21] and purity was checked through the absorbance ratio $A_{290}/A_{260}$. Highly concentrated and >90% pure fractions ($A_{290}/A_{260}$ = 1.1–1.2 accounts for >95% purity)[22] were lyophilized and stored at −70°C.

##### Production of auxiliary enzymes HisG/IE and HisA

HisG/IE was produced by heterologous gene expression in *E. coli* BL21 Gold (DE3) (Agilent Technologies) using the p_hisGIE_tac plasmid as described previously.[20] After transformation, the cells were grown in 2 L lysogeny broth (LB) medium at 37°C until reaching an $OD_{600}$ of 0.7, at which point protein expression was induced with 1 mM IPTG and incubated overnight at 30°C. After harvesting by centrifugation, the bacterial pellets were resuspended in 10 mM KP (pH 7.5), 2.5 mM EDTA, and 1 mM DTT and lysed by sonication. After repeated centrifugation, the protein-containing supernatant was further purified using ion-exchange chromatography (MonoQ HR 16/10, 20 mL, Pharmacia). Proteins were eluted using a linear gradient of KP (10–500 mM), and fractions containing HisG or HisIE or both were pooled and dialyzed against 50 mM KP (pH 7.5), 2.5 mM EDTA, and 1 mM DTT. Proteins were identified and checked for >90% purity using SDS-PAGE analysis. Concentrated proteins were flash-frozen in liquid nitrogen and stored at −70°C.

HisA was produced by heterologous gene expression in *E. coli* BL21 Gold (DE3) (Agilent Technologies) using the pET21_HisA plasmid as described previously.[23] The recombinant protein was purified from the soluble fraction of the crude extract through heat precipitation (15 min at 73°C) to remove host proteins, followed by nickel-affinity chromatography. The extract was loaded onto a HisTrap FF crude (5 mL, GE Healthcare) column, pre-equilibrated with 50 mM Tris-HCl (pH 7.5), 300 mM NaCl, and 10 mM imidazole. After washing with equilibration buffer, bound protein was eluted using a linear gradient of 10–750 mM imidazole. Pure fractions were pooled and dialyzed against 50 mM Tris-HCl (pH 7.8). A purity of at least 95% was confirmed by SDS-PAGE analysis. Proteins were flash-frozen in liquid nitrogen and stored at −70°C.

**Site-directed mutagenesis and Golden Gate cloning**

To introduce site-directed substitutions or deletions, particularly for the preparation of HisF and hAzoF gene libraries via scanning mutagenesis, or to prepare plasmids and gene fragments for Golden Gate cloning[24] by preparing them with BsaI or BbsI restriction sites, site directed mutagenesis was performed based on a protocol of the Phusion™ site directed mutagenesis kit from Finnzymes (Thermo Fisher Scientific) with HPLC purified primers (Metabion). For substitutions, forward and reverse primers were designed to start elongation in opposite directions, with their 5' ends directly next to each other. One primer carried the desired mutation. For deletions, primers starting elongation in opposite directions and skipping the desired sequence seamlessly were designed. PCR resulted in linear amplicons that were re-circularized in a coupled phosphorylation-ligation reaction using T4 Polynucleotide Kinase and T4 DNA Ligase (Thermo Fisher Scientific) according to the manufacturer's instructions. Integrity of the plasmids and correct mutagenesis was confirmed by Sanger Sequencing (Microsynth Seqlab).

For subcloning synthetic genes or amplified sequences into plasmid vectors, Golden Gate cloning[24] was performed. This highly efficient method uses type IIS restriction enzymes (e.g., BsaI, BbsI) to assemble multiple DNA fragments. These enzymes cut outside their recognition sites, creating specific overhangs. After ligation, the recognition sites are removed, preventing further cleavage. Therefore, one-pot restriction-ligation reaction as described by Rohweder *et al.*[24] was carried out with the fragments or plasmids containing suitable recognition sites. Correct sequence of the final plasmid was confirmed by Sanger Sequencing (Microsynth Seqlab).

**Subcloning of pEVOL-AapF**

A pEVOL plasmid with two copies of a *Methanocaldococcus janaschii* TyrRS and its respective tRNA provided by Peter Schultz (Scripps Research Institute, La Jolla, CA)[18] was used as backbone. Both copies of the TyrRS were removed and replaced with BbsI restriction sites for Golden Gate cloning via site-directed mutagenesis. A previously designed *Mj*TyrRS gene for the incorporation of AapF[25] was codon optimized for *E. coli*, synthesized with BbsI restriction sites (GeneArt, Thermo Fisher Scientific), and cloned into the prepared pEVOL plasmid via Golden Gate cloning. The correct sequence of the entire plasmid was confirmed by Sanger Sequencing (Microsynth Seqlab).

**Subcloning for the split-GFP assay**

The gene encoding GFP1-10 was codon optimized for *E. coli*, synthesized with BsaI restriction sites (GeneArt, Thermo Fisher Scientific), and cloned into a pET21a_BsaI vector via Golden Gate cloning.

The gene encoding HisF$^{s11}$ was codon optimized for *E. coli*, synthesized with BsaI restriction sites (GeneArt, Thermo Fisher Scientific), and cloned into a pET28aTEV_BsaI vector[14] via Golden Gate cloning. Thereby, the s11 tag was fused to HisF C-terminally via a linker to avoid interference with the N-terminal His-tag.

To create a pET28a_s11 vector that can be used to attach a C-terminal s11 tag to any gene via Golden Gate cloning, the *hisF* gene was removed from the plasmid pET28a_HisF$^{s11}$ and replaced with BsaI restriction sites via site-directed mutagenesis.

For the production of HisH$^{s11}$ variants, the *hisH* gene including a N-terminal His-tag was amplified with BsaI restriction sites from pET28a_HisH[14] and cloned into the prepared plasmid via GoldenGate cloning. Finally, a TAG point mutation at position W123 was introduced into pET28a_HisH$^{s11}$ via site-directed mutagenesis to obtain pET28a_HisH-W123TAG$^{s11}$. The correct sequences of the new pET21a_GFP1-10, pET28a_HisF$^{s11}$, pET28a_s11, and pET28a_HisF$^{s11}$ plasmids was confirmed by Sanger Sequencing (Microsynth Seqlab).

**Scanning mutagenesis of HisF and hAzoF**

Selected point mutations were introduced in pET28a_HisH-W123TAG$^{s11}$ and pET28a_HisF$^{s11}$ via site-directed mutagenesis to generate DNA libraries for screening purposes (vide infra). Mutations identified from screening hits were further cloned into pET28a_HisH-W123TAG[12] and pET28a_HisF, which was generated by removing the s11-tag via site-directed mutagenesis. Correct mutagenesis of the new plasmids was confirmed by Sanger Sequencing (Microsynth Seqlab) starting from the T7 promotor or terminator.

**Production and purification of HisF and HisH variants without UAA**

Production of wildtype proteins and variants without UAA was performed in *E. coli* BL21 Gold (DE3) cells (Agilent Technologies) by heterologous gene expression. The cells were transformed with the respective plasmids and grown overnight in a preculture of LB medium supplemented with 50 mg/L kanamycin. The next day, 6 L of LB medium (supplemented with 50 mg/L kanamycin) were inoculated with the preculture to an $OD_{600}$ of ~0.1 and grown at 37°C. As soon as an $OD_{600}$ of ~0.6 was reached, protein expression was induced with 0.5 mM IPTG and proceeded at 30°C overnight. Cells were harvested by centrifugation, resuspended in either 50 mM Tris-HCl (pH 7.5), 100 mM NaCl, and 10 mM imidazole (HisF variants) or 50 mM KP (pH 7.5), 100 mM NaCl, and 10 mM imidazole (HisH variants) and lysed by sonication. *E. coli* proteins were precipitated by a heat step (15 min, 60°C) and the proteins of interest (POI) were obtained from the supernatant after subsequent centrifugation. The supernatant was subjected to nickel-affinity chromatography (HisTrap_FF Crude column, 5 mL, GE Healthcare) and bound protein was eluted with a linear gradient of 10–750 mM imidazole. Fractions containing the POI were identified by SDS-PAGE, pooled, and concentrated. HisH variants were further purified via preparative size-exclusion chromatography (Superdex 75 HiLoad 26/600, GE Healthcare) using 50 mM KP (pH 7.5), 100 mM NaCl as running buffer. Fractions containing the POI with a purity of >90% were identified by means of SDS-PAGE, pooled, concentrated, flash-frozen in liquid nitrogen and stored at −70 °C.

HisF samples were split after nickel-affinity chromatography. One part was further purified via preparative size-exclusion chromatography (Superdex 75 HiLoad 26/600, GE Healthcare) analogous to HisH using 50 mM Tris-HCl (pH 7.5), 100 mM NaCl (HisF variants) or 50 mM KP (pH 7.5), 100 mM NaCl (HisH variants) as running buffer. The other part was dialyzed in 50 mM Tris-HCl (pH 7.5), 100 mM NaCl supplemented with 2 mg of TEV protease per ~10 mL of dialysate, overnight at room temperature. The remaining TEV protease was then removed by reverse nickel-affinity chromatography (HisTrap FF Crude column, 5 mL, GE Healthcare). Unbound, TEV digested POI was eluted in 50 mM Tris-HCl (pH 7.5), 100 mM NaCl and the rest with a linear gradient of 0–750 mM imidazole. Fractions containing the POI with a purity of >90% were identified by means of SDS-PAGE, pooled, concentrated, flash-frozen in liquid nitrogen and stored at −70 °C.

**Production and purification of hAzoF and hAapF**

For the production of hAzoF and hAapF a slightly adjusted protocol was used. Preparation of the expression strain thereby included the co-transformation of the respective pET28a_HisH-W123TAG plasmid and either pEVOL_AzoF-RS[18] or pEVOL_AapF-RS,[25] harboring the orthogonal aminoacyl-tRNA synthetase adapted for AzoF or AapF binding as well as the respective orthogonal tRNA required for AzoF incorporation. After inoculating 6 L of LB medium supplemented with 30 mg/L chloramphenicol and 50 mg/L kanamycin with a preculture, cells were grown at 37°C to an $OD_{600}$ of ~0.6. The cells were then harvested by centrifugation and resuspended in 600 mL terrific broth (TB) medium. Bacterial growth was resumed at 37°C until an $OD_{600}$ of ~10. Then, protein expression was induced by adding 0.5 mM IPTG, 0.02% L-arabinose, and 0.8 mM AzoF. The culture was incubated overnight at 25°C. Protein purification was performed in the dark, following the same procedure as described above for HisH variants without UAA.

**Production and purification of HisF$^{CouA}$ variants**

For production of HisF variants containing CouA, the protocol for expression of hAzoF and hAapF was used with small adjustments: Co-transformation included the appropriate plasmid encoding the gene of interest and pEVOL-CouA-RS, harboring the orthogonal aminoacyl-tRNA synthetase adapted for CouA binding as well as the respective orthogonal tRNA required for CouA incorporation.[11] Protein expression was induced by adding 0.5 mM IPTG, 0.02% L-arabinose, and 0.45 mM CouA and was proceeded at 30°C overnight. The cells were harvested, lysed, and further purified as described above for HisF variants without TEV digestion.

**Production and purification of GFP1–10**

GFP1–10 reporter solution was prepared following a standard protocol described previously.[3] Briefly, *E. coli* BL21 Gold (DE3) cells (Agilent Technologies) were transformed with the pET21a_GFP1–10 plasmid and grown overnight in a preculture of LB medium supplemented with 150 mg/L ampicillin. The preculture was used to inoculate 500 mL LB medium supplemented with ampicillin to an $OD_{600}$ of 0.1 and the cells were incubated at 37°C until an $OD_{600}$ of ~0.6 was reached. Subsequently, gene expression was induced by adding 1 mM IPTG, following incubation at 37°C for 3 h. Cells were harvested by centrifugation, resuspended in cold TNG buffer [100 mM Tris-HCl (pH 7.4), 150 mM NaCl, 10% (v/v) glycerol] and lysed by sonication. A repeated centrifugation step was performed to recover the inclusion bodies. The pellet was resuspended in 1 × FastBreak™ Cell Lysis Reagent (Promega GmbH) in TNG buffer, sonicated and centrifuged, and the supernatant was discarded. This step was repeated three times. Subsequently, TNG buffer was added, followed by sonication and centrifugation to remove detergent. The supernatant was discarded, and these steps were repeated two times. TNG buffer was added to the inclusion bodies to reach a final concentration of 75 mg/mL, separated in 1 mL volumes and centrifuged. The pellet was frozen at −70°C. To refold the inclusion bodies, the pellet was dissolved in 1 mL 9 M urea, 5 mM DTT at 37°C. The resuspension was centrifuged to precipitate any misfolded GFP1–10 aggregates and subsequently 25 mL cold TNG buffer were added, gently mixed and passed through a 0.2 µM syringe filter (Sarstedt). The reporter solution was stored up to two months at −70°C.

**Screening of DNA libraries**

*E. coli* BL21 Gold (DE3) cells (Agilent Technologies) were individually co-transformed with pEVOL_AzoF-RS and one plasmid from each DNA library. To produce positive and negative controls, samples of the same expression host were co-transformed with pEVOL_AzoF-RS and either pET28a_HisF$^{s11}$ or pET28a_HisH$^{s11}$, or pET28aTEV_BsaI, which lacks a target gene insert,[14] respectively. Each co-transformed cell sample was then cultivated on an LB agar plate supplemented with 50 mg/L kanamycin and 30 mg/L chloramphenicol. To prepare precultures for expression, four colonies of each variant or control were used to inoculate 4 × 1.8 mL LB medium supplemented with the respective antibiotics in a 96-deepwell plate. The plate was covered with a gas permeable membrane (Moisture Barrier Seal 96, Azenta Life Science) and incubated overnight at 37°C in a microtiter plate shaker (Hei-MIX Titramax 1000 and Hei-MIX Incubator 1000, Heidolph Instruments). For heterologous gene expression of the enzyme library each preculture was diluted 1:16 in TB medium with the respective antibiotics to a final volume of 200 µL in two flat-bottom 96-well plates, generating technical duplicates of the four biological replicates. The *E. coli* cultures were incubated until $OD_{600}$ reached ~0.4–0.8 in the covered plate, then transferred to a canonical 96-well plate and heterologous gene expression was induced with 0.5 mM IPTG, 0.02 % L-arabinose, and 0.4 mM AzoF. After overnight incubation at 30°C in the microtiter plate shaker, which was covered to keep the plates in the dark, cells were harvested by centrifugation. This step and all subsequent steps were performed in the dark. Cell lysis with 100 µL 1 × FastBreak™ Cell Lysis Reagent (Promega GmbH) in 50 mM Tris-HCl (pH 7.5) and a heat step for purification at 60°C for 15 min, followed by centrifugation to separate cell debris yielded the individual variants in 2 × 4 biological replicates.

Each resulting enzyme library was screened using the real-time photocontrol activity assay described below. For this, the enzyme complex was assembled by adding 2 µM HisF or hAzoF in its TEQ to a total volume of 40 µL lysate containing one hAzoF$^{s11}$ (in its TEQ) or HisF$^{s11}$ variant, respectively, of the enzyme library. Glutaminase activity was then initiated by addition of 10 µL prepared enzyme complex in its TEQ state. To normalize the determined activity of each enzyme clone with its expression level, a split-GFP assay[1–3] was performed. The truncated GFP1–10 protein was prepared as described above, and 180 µL were mixed with 20 µL lysate in a black 96-well plate. After incubation at 4°C for at least 12 h, fluorescence intensity was measured using a plate reader (Tecan Infinite M200

Pro) with an excitation wavelength of 488 nm and an emission wavelength of 530 nm. The detector's amplification factor was set to optimum, and the mean fluorescence intensity was calculated from five measurement cycles. The activity *v* of each variant of the enzyme library was then normalized with its corresponding fluorescence intensity from the split-GFP assay. Finally, LRFs and reversibility were calculated from the means of all four replicates as described below (**Equation S2**).

**Real-time photocontrol activity assay**

Photocontrol of glutaminase activity was recorded in real-time using a coupled enzymatic assay for the detection of the ProFAR stimulated glutaminase activity of the HisH subunit (**Figure S5**). For this, L-glutamate oxidase (GOX, Sigma-Aldrich) and horse radish peroxidase (HRP, Sigma-Aldrich) were employed as auxiliary enzymes producing red-colored quinoneimine as final product. All reaction components—except the enzyme complex—were pre-mixed, and 90 µL were filled into each well of a transparent flat-bottom 96-well plate and incubated for at least 1 h at room temperature to reduce possible L-glutamate contaminations. Reaction conditions for the screening included low concentrations of auxiliary enzymes to reduce costs: 5 mM L-glutamine (saturated), 40 mU/mL GOX, 80 U/mL HRP, 3 mM 4-aminoantipyrine, 3 mM phenol, 70 µM ProFAR (saturated) in 20 mM Tris-HCl (pH 7.0 at 28°C). To obtain exact values, reaction conditions with purified enzymes contained increased auxiliary enzyme concentrations: 10 mM L-glutamine (saturated), 150 mU/mL GOX, 120 U/mL HRP, 3 mM 4-aminoantipyrine, 3 mM phenol, 70 µM ProFAR (saturated) in 20 mM Tris-HCl (pH 7.0 at 28°C). For reactions in the absence of the allosteric stimulator ProFAR the concentrations of enzyme complexes were increased to 2.4 µM.

After recording the baseline absorbance of the reaction components for several minutes at 505 nm and 28°C in a plate reader (Tecan Infinite M200 Pro), the reaction was initiated by addition of the prepared enzyme complex in its TEQ state. For reactions with purified enzymes 0.05 µM for hAzoF and all single mutants, and 0.1 µM for all double and triple mutants were employed. The reaction was then followed throughout the lag phase, typical for coupled enzymatic assays, and several minutes of the pseudo-linear steady state phase. Measurements were then paused, and the reactions were irradiated with 365 nm (2 min per plate with the screening setup or 2 s per well with the individual setup for purified enzymes) to establish the *Z*-enriched $PSS^{365}$. This step was repeated a second time using 420 nm irradiation (2 min per plate with the screening setup or 4 s per well with the individual setup for purified enzymes) to reestablish an *E*-enriched $PSS^{420}$. To determine activity values *v* the quasi-linear phases of each reaction were fitted with a linear regression model and the resulting slopes *m* were converted into *v* using the Lambert-Beer law (**Equation S1**),

$$v = \frac{\Delta c}{\Delta t} = \frac{\Delta A}{\Delta t \times \varepsilon(quinoneimine_{505nm}) \times d} = \frac{m}{\varepsilon(quinoneimine_{505nm}) \times d} \quad \text{(S1)}$$

with *Δc* as the change in quinoneimine concentration over time (*Δt*), *ΔA* as the corresponding change in absorbance, $\varepsilon(quinioneimine_{505nm})$ as the extinction coefficient of quinonimine with the value 6400 $M^{-1}cm^{-1}$,[26] and *d* as the pathlength with an approximate value of 2.67 mm as calculated from technical specifications of the 96-well plate. For reactions using purified enzymes, normalization using the applied enzyme concentration obtained $v/E_0$ values. Mean values and their standard deviations (SD) were calculated from all applied replicates and LRFs comparing TEQ and $PSS^{365}$ were calculated from the mean *v* or $v/E_0$ values using **Equation S2**, with *y* as the respective activity constant.

$$LRF = \frac{y_{PSS^{365}}}{y_{TEQ}} \quad \text{S2}$$

Reversibility (revs.) was determined as the percentage by which activity decreased after exposure to 420 nm light using **Equation S3** with *y* as the respective mean activity constant (unrounded).

$$revs.[\%] = 1 - \frac{y_{PSS^{420}} - y_{TEQ}}{y_{PSS^{365}} - y_{TEQ}} * 100 \quad \text{S3}$$

**Z'-factor determination**

To estimate the data quality of the screening assay, the Z'-factor was determined according to **Equation S4**,[27]

$$Z' - factor = 1 - \frac{3\,(\sigma_p + \sigma_n)}{|\mu_p - \mu_n|} \quad \text{S4}$$

with $\sigma_p$ and $\sigma_n$ as the standard deviations of positive and negative control, respectively, and $\mu_p$ and $\mu_n$ as the means of positive and negative control, respectively. Generally, a Z'-factor of 1 is considered ideal, with values between 0.5 and 1 indicating an excellent assay. Values between 0 and 0.5 suggest a marginal assay and a Z'-factor below 0 indicates that the screening is essentially impossible.[27]

To determine the Z'-factor for the screening assay developed to evaluate LRFs, the entire screening process was performed for two 96-well plates containing the positive and negative control, respectively. As positive control an enzyme complex consisting of HisF and lysate from hAzoF$^{s11}$, and as negative control an enzyme complex consisting of HisF and lysate from HisH$^{s11}$ was used. Glutaminase activity measurements and evaluations were conducted as described for the screening and for the real-time photocontrol activity assay.

**Alanine screen of ProFAR-stimulated glutaminase activity**

To select residues for the semi-rational design that are located at the interface and relevant for allostery, an alanine screen was performed. For this, the crystal structure of apo HisH_HisF (PDB: 1GPW) was compared with the crystal structure of holo HisH_HisF (PDB: 7AC8). Five residue positions were identified that are located at the interface with a considerable change of the sidechain conformation rather than the backbone conformation/location. The selected residue positions in HisF and HisH were mutated to alanine using site-directed mutagenesis and the resulting variants were produced using heterologous gene expression as described above. ProFAR stimulated glutaminase activity of the obtained HisF variants in complex with HisH or the obtained HisH variants in complex with HisF was measured with the coupled enzymatic assay (**Figure S5**) using the following reaction conditions: 10 mM L-glutamine (saturated), 20 mU/mL GOX, 25 U/mL HRP, 1 mM 4-aminoantipyrine, 1 mM phenol, 100 µM ProFAR (saturated) in 20 mM Tris-HCl (pH 7.0 at 28°C). The reactions were started by addition of 0.2 µM enzyme complex and performed in technical triplicates at 505 nm and 28°C. Activity values $v$ and $v/E_0$ were determined as described above using the extinction coefficient of 6400 $M^{-1}$ $cm^{-1}$.[26]

**Cycle performance of enzymatic photocontrol**

To show the reversibility of photocontrol in multiple circles of irradiation, further increase of the concentrations of auxiliary enzymes in the real-time photocontrol activity assay was required to minimize the lag phase (300 mU/mL GOX, 190 U/mL HRP). The measurements were performed as described for purified enzymes in the real-time photocontrol activity assay with the following adaption. When reaching the linear steady state phase, the measurement was paused and irradiated six times with alternating 365 nm and 420 nm irradiation. The reactions were measured in technical duplicates at 505 nm and 28°C for ~1.5 h. Negative control reactions (single measurements) were run in parallel that were not irradiated. The activity values *v* and $v/E_0$ as well as the LRF and revs. were obtained as described above using the extinction coefficient of 6400 $M^{-1}$ $cm^{-1}$.[26]

**Steady-state kinetics of ProFAR-stimulated HisH-activity**

To determine steady-state constants of the ProFAR stimulated glutaminase activity the coupled enzymatic assay as described above was utilized (**Figure S5**). Reaction conditions included: 0.05–20 mM L-glutamine, 100 mU/mL GOX, 120 U/mL HRP, 3 mM 4-aminoantipyrine, 3 mM phenol, 70 µM ProFAR (saturated) in 20 mM Tris-HCl (pH 7.0 at 28°C). The reaction was initiated by addition of the prepared enzyme complex (TEQ: 0.05 µM for hAzoF_HisF and hAzoF_fI73P; 0.1 µM for hAzoF_fI75K; 0.2 µM for hAzoF_fL2D-fI75K; 0.3 µM for hAzoF_fL2D; 0.5 µM for hAzoF_fI73P-fI75K; 0.6 µM for hAzoF_fL2D-fI73P, and 1.2 µM for hAzoF_fL2D-fI73P-fI75K; $PSS^{365}$ and $PSS^{420}$: 0.05 µM for hAzoF_HisF, hAzoF_fI73P, and hAzoF_fI75K; 0.1 µM for hAzoF_fL2D; 0.2 µM for hAzoF_fL2D-fI73P, hAzoF_fL2D-fI75K, and hAzoF_fI73P-fI75K; 0.6 µM for hAzoF_fL2D-fI73P-fI75K). To determine steady-state constants for all variants in the TEQ state the reaction was performed without irradiation. To determine steady-state constants for all variants in the $PSS^{365}$ and $PSS^{420}$ the reaction mixture was irradiated for 2 s per well with 365 nm or 4 s per well with 420 nm (individual irradiation setup) directly after addition of the enzyme complex. For $PSS^{420}$ the samples were irradiated first with 365 nm and second with 420 nm. For each substrate concentration two technical replicates were measured. $v/E_0$ values were obtained as described above for the real-time photocontrol activity assay. The $v/E_0$ values of the duplicates were then plotted against the substrate concentration *c(S)* and fitted together in a concatenated non-linear regression model following the Michaelis-Menten law (**Equation S5**) to determine $k_{cat}$ ± standard error of fit (SE) and $K_m$ ± SE values.

$$v/E_0 = \frac{k_{cat} \times c(S)}{K_m + c(S)} \qquad \text{S5}$$

The catalytic efficiency $k_{cat}/K_m$ ± SE was deduced by fitting the duplicate values using a converted version of **Equation S5** (**Equation S6**).

$$v/E_0 = \frac{k_{cat}/K_m \times c(S)}{1 + \frac{k_{cat}/K_m \times c(S)}{k_{cat}}} \qquad \text{S6}$$

**UV/Vis analysis**

UV/Vis spectra of 25 µM enzyme complex in 20 mM Tris-HCl (pH 7.0 at 28°C) or 20 µM hAapF in 50 mM KP (pH 7.5), 100 mM NaCl were recorded in a UV-transparent 96-well plate in the range of 240–600 nm using a plate reader (Tecan Infinite M200 Pro). Spectra were either measured in the TEQ state or after subsequent irradiation with 365 nm or 420 nm for different durations (individual irradiation setup). All spectra were baseline corrected at 600 nm. To determine the rates of isomerization *k* the absorbance values at the π→π* transition were plotted against the irradiation time. Exponential decay of the absorbance was then fitted with **Equation S7** to obtain the rate of isomerization $k^{365}$,

$$y = y_i + A\,e^{-k^{365}t} \qquad \text{S7}$$

in which *t* is the irradiation time, $y_i$ is the *y* value at infinite times (also named plateau) and *A* is the span of the exponential curve between $y_i$ and $y_0$ (the *y* value when time = 0). Exponential growth of the absorbance was fitted with **Equation S8** to obtain the rate of isomerization $k^{420}$.

$$y = y_0 + A\,(1 - e^{-k^{420}t}) \qquad \text{S8}$$

The half-life of isomerization $t_{1/2}$ was then derived from each isomerization rate *k* with **Equation S9**.

$$t_{1/2} = \frac{\ln(2)}{k} \qquad \text{S9}$$

**Estimation of *E*:*Z* ratios from UV/Vis spectra**

Estimation of *E*:*Z* ratios followed an expanded procedure based on previously published protocols.[28–30] First, several spectra spanning various time-points between the TEQ state and $PSS^{365}$ were selected from one experiment and the π→π* and n→π* peaks were deconvoluted between 310 nm and 600 nm in Origin 2024 with the multiple peak fit tool using the Gaussian function shown in **Equation S10**,

$$y = A\,e^{\frac{(x-x_c)^2}{2w^2}} \qquad \text{S10}$$

in which $y$ is the measured absorbance, $A$ is the amplitude of the Gauss curve, $x$ is the wavelength, $x_c$ is the wavelength at the center of the Gauss curve, and $w$ is the width of the Gauss curve at $\frac{A}{2}$. As a result, a cumulative Gauss fit of both peaks was created that simulates the respective UV/Vis spectrum. Next, a first estimation of the *E*:*Z* ratios was obtained by using the cumulative Gauss fits. The cumulative Gauss fit of the TEQ spectrum was used as reference, assuming that the TEQ state comprises 100% of *E* isomer. The spectrum of 100% *Z* isomer was simulated to determine the fraction of *E* in another selected cumulative Gauss fit with the simple curve math tool in Origin 2024 using **Equation S11**,

$$A_Z = \frac{A_i - A_E \times f_E}{1 - f_E} \qquad \text{S11}$$

in which $A_Z$ is the simulated spectrum of the 100% *Z* isomer, $A_E$ is the cumulative Gauss fit of the TEQ (100% *E*) reference spectrum, $A_i$ is the selected cumulative Gauss fit, and $f_E$ is the fraction of the *E* isomer. The fraction of $f_E$ was obtained in this simulation by manual adjustment until the signal of the π→π* absorbance band approximated zero.

Finally, a standard curve for each experiment was generated by plotting the estimated $f_E$ values against the absorbance value at the π→π* maximum of the cumulative Gauss fit from the original spectra. Linear regression of this standard curve obtained a linear fit equation that was used to estimate $f_E$ of any spectrum within the experiment that were then utilized to determine rate constants.

**Fluorescence analysis of complex formation with HisF$^{CouA}$**

Dilution series with HisF$^{CouA}$ variants, in which the fluorescent UAA CouA was incorporated in position K99 in HisF variants, were performed to evaluate complex formation with HisH or hAzoF. For the measurements, a dilution series of HisH or hAzoF was prepared (0–1000 nM) and mixed in a 96-well plate with either HisF$^{CouA}$ variant (100 nM) in 50 mM Tris-HCl (pH 7.5). Subsequently, CouA fluorescence was monitored in a plate reader (Tecan Infinite M200 Pro) at 25°C using 325 nm excitation and 424 nm emission wavelengths. The measurements were carried out twice with the same detector's amplification factor for two technical replicates of each HisH or hAzoF concentration. Besides looking at the TEQ state of the enzymes, the measurements were also performed with pre-irradiated hAzoF (2 s per well with 365 nm). Fluorescence signals (*F*) of all replicates were plotted as a function of the HisH or hAzoF concentration (*L*) and fitted together with a hyperbolic equation in Origin 2024 using **Equation S12** to obtain $K_D$ values,

$$F = F_{start} + \frac{1}{2} \times (F_{max} - F_{start}) \times \left(1 + \frac{L + K_D}{E} - \sqrt{\left(1 + \frac{L + K_D}{E}\right)^2 - 4 \times \frac{L}{E}}\right) \quad \text{S12}$$

in which $F_{start}$ is the initial fluorescence signal, $F_{max}$ is the fluorescence intensity at saturation and $E$ is the fixed concentration of HisF$^{CouA}$ (100 nM).

**Determination of $K_{ac}^{ProFAR}$ values**

$K_{ac}^{ProFAR}$ values were measured using the coupled enzymatic assay (**Figure S5**) for the glutaminase reaction. Reaction conditions included: 10 mM L-glutamine (saturated), 100 mU/mL GOX, 120 U/mL HRP, 3 mM 4-aminoantipyrine, 3 mM phenol, 0–100 µM ProFAR in 20 mM Tris-HCl (pH 7.0 at 28°C). The reaction was initiated by addition of the prepared enzyme complex (0.05 µM for HisH_HisF and hAzoF_HisF; 0.1 µM for hAzoF_fL2D and hAzoF_fI73P-fI75K) in its TEQ or PSS$^{365}$, for which the enzyme complexes were pre-irradiated (10 s per well with 365 nm; individual irradiation setup) before addition to the reaction. Determined $v/E_0$ values (as described for the real-time photocontrol activity assay) were plotted against the ProFAR concentration and fitted in Origin 2024 with the hyperbolic **Equation S12** with $F$ as $v/E_0$ values, $F_{start}$ as initial $v/E_0$ values, $F_{max}$ as $v/E_0$ values at saturation, $E$ as fixed enzyme complex concentration, $L$ as ProFAR concentration and $K_D$ as $K_{ac}^{ProFAR}$.

**Determination of $K_{ac}^{hAzoF}$ values and $k_{cat}^{app}$ of HisF**

To evaluate the influence of hAzoF on HisF, the coupled HisH_HisF activity was determined by following the turnover of PRFAR to AICAR and ImGP in the presence of glutamine as a source of ammonia (supplied by HisH; **Figure S20**). To this end, the decline of the PRFAR absorbance signal at 300 nm was recorded at 25°C using UV-transparent 96-well plates and a plate reader (Tecan Infinite M200 Pro). Reaction conditions included: 1 µM HisA (to turn over ProFAR to PRFAR), 70 µM ProFAR (saturated), 10 mM Gln (saturated) in 20 mM Tris-HCl (pH 7.0 at 28°C). To determine the $K_{ac}^{hAzoF}$ values, a dilution series of hAzoF was prepared and was added to the reaction components to obtain final concentrations of 0.01–0.75 µM. Moreover, hAzoF was either used in its TEQ, PSS$^{365}$ (30 s 365 nm) or PSS$^{420}$ (30 s 365 nm and subsequent 60 s 420 nm) via pre-irradiation using the individual irradiation setup. After recording the baseline absorbance of the reaction components for several minutes, the reaction was initiated by addition of 0.1 µM HisF variant. Technical duplicates were measured for each concentration of hAzoF. To determine activity values $v$, the quasi-linear phases of each reaction were fitted with a linear regression and the resulting slopes *m* were converted into *v* using the Lambert-Beer law (**Equation S1**) with $\Delta c$ as the turnover of PRFAR over time ($\Delta t$), $\Delta A$ as the corresponding change in absorbance, $\varepsilon(PRFAR\text{-}AICAR_{300nm})$ as the extinction coefficient with the value 5637 $M^{-1}cm^{-1}$,[31] and *d* as the pathlength with an approximate value of 2.67 mm. The obtained *v* values for the duplicates were plotted against the hAzoF concentration and fitted together with the hyperbolic **Equation S12** with *F* as *v* values, $F_{start}$ as initial *v* values, $F_{max}$ as *v* values at saturation, *E* as fixed HisF variant concentration, *L* as hAzoF concentration and $K_D$ as $K_{ac}^{hAzoF}$. To determine $k_{cat}^{app}$ the $F_{max}$ values were normalized with the enzyme concentration.

**Determination of $K_{ac}^{HisF}$ values and $k_{cat}^{app}$ of ProFAR stimulated hAzoF**

To evaluate the influence of HisF on HisH, the coupled enzymatic assay for the glutaminase reaction as described the real-time photocontrol activity assay was used. Reaction conditions included: 10 mM L-glutamine (saturated), 80 mU/mL GOX, 100 U/mL HRP, 3 mM 4-aminoantipyrine, 3 mM phenol, 80 µM ProFAR (saturated) in 20 mM Tris-HCl (pH 7.0 at 28°C). Moreover, hAzoF (0.1 µM) was added to the assay components, either in TEQ or $PSS^{365}$ (pre-irradiated 30 s 365 nm with the individual irradiation setup). After recording the baseline, the reactions were initiated by addition of a dilution series of HisF monomer (0.001–1 µM). Activity values *v* were determined as described above and $K_{ac}^{HisF}$ values were obtained using the same method as described for $K_{ac}^{hAzoF}$, and $k_{cat}^{app}$ values correspond to $F_{max}$ values normalized with the enzyme concentration.

**Circular dichroism analysis**

Circular dichroism (CD) spectra in the far-UV range of 195−280 nm were recorded in a Jasco J-815 spectrophotometer with five accumulations. The spectra were measured with 30 µM enzyme complex in 50 mM KP (pH 7.5) in a 0.02 cm cuvette at 25°C. The curves were smoothed in Origin 2024 (OriginLab). Data were normalized to obtain the mean residue ellipticity as described in ref[32].

**Differential scanning calorimetry**

Differential scanning calorimetry (DSC) measurements were performed in a VPDSC differential scanning microcalorimeter (MicroCal, Malvern Instruments) with fixed reference and sample cells (0.545 mL each). Each sample was degassed for several minutes before measurement. Proper equilibration of the calorimeter was initially ensured by conducting several buffer-buffer baselines. DSC runs were then performed using 20 µM enzyme complex in 50 mM KP (pH 7.5) and 100 mM NaCl that was heated from 40 to 120°C at a ramp rate of 1 K $min^{-1}$. Overpressure was applied to prevent boiling above 100°C. The change in heat capacity with an increase in temperature was recorded, the buffer signal was subtracted from the protein data, and baselines were corrected using the spline interpolation option in Origin 2024. The denaturation midpoint $T_m$ was determined as the temperature at the maximum of the change in heat capacity.

**Mass spectrometry**

Recombinant *Thermatoga maritima* hAzoF and hAapF proteins were run on a 10% SDS-PAGE and stained with Coomassie G250 (SimplyBlue SafeStain, Lifetech). Protein bands were cut out from the gel, washed with 50 mM $NH_4HCO_3$, 50 mM $NH_4HCO_3$/acetonitrile (3/1), 50 mM $NH_4HCO_3$/acetonitrile (1/1) and lyophilized. After a reduction/alkylation treatment and additional washing steps, proteins were in gel digested with trypsin (Trypsin Gold, mass spectrometry grade, Promega) overnight at 37°C. The resulting peptides were sequentially extracted with 50 mM $NH_4HCO_3$ and 50 mM $NH_4HCO_3$ in 50% acetonitrile. After lyophilization, peptides were reconstituted in 20 µL 1% TFA and separated by reversed-phase chromatography. An UltiMate 3000 RSLCnano System (Thermo Fisher Scientific, Dreieich) equipped with a C18 Acclaim Pepmap100 preconcentration column (100 µm i.D.×20 mm, Thermo Fisher Scientific) and an Acclaim Pepmap100 C18 nano column (75 µm i.d.×250 mm, Thermo Fisher Scientific) was operated at flow rate of 300 nl/min and a 60 min linear gradient of 4 % to 40 % acetonitrile in 0.1 % formic acid. The LC was online-coupled to a maXis plus UHR-QTOF System (Bruker Daltonics) via a CaptiveSpray nanoflow electrospray source. Acquisition of MS/MS spectra after CID fragmentation was performed in data-dependent mode at a resolution of 60000. The precursor scan rate was 2 Hz processing a mass range between m/z 175 and m/z 2000. A dynamic method with a fixed cycle time of 3 s was applied via the Compass 1.7 acquisition and processing software (Bruker Daltonics). Prior to database searching with Protein Scape 3.1.3 (Bruker Daltonics) connected to Mascot 2.5.1 (Matrix Science), raw data were processed in Data Analysis 4.2 (Bruker Daltonics). A customized database comprising the *Thermatoga maritima* entries from UniProt as well as manually added sequences of the mutated HisH proteins and common contaminants, was used for database search with the following parameters: enzyme specificity trypsin with 2 missed cleavages allowed, precursor tolerance 10 ppm, MS/MS tolerance 0.04 Da. As general variable modifications, deamidation (0.984 Da) of asparagine and glutamine, oxidation of methionine (15.995 Da), carbamidomethylation of cysteine (57.021 Da), or propionamide modification of cysteine (71.037 Da) were set. Specific variable modifications for identification of unnatural amino acids were as follows:

AapF 136.075 Da, or AzoF 104.037 Da. Spectra of peptides containing unnatural amino acids were inspected manually.

**Molecular modelling system preparation**

In preparation for an extensive evaluation of hAzoF_HisF, hAzoF_fL2D, and hAzoF_fI73P-fI75K via molecular dynamics (MD) simulations, we validated that a comparable photocontrol effect is maintained in all three variants after removal of the N-terminal $His_6$-tag, which is difficult to simulate owing to its high flexibility (**Table S9**).

The X-ray structure available for HisH_HisF in its active form (PDB: 7AC8, chains E and F) was used to generate the starting structures for the six systems (hAzoF_HisF *E*, hAzoF_HisF Z, hAzoF_fL2D *E*, hAzoF_fL2D *Z*, hAzoF_fI73P-fI75K *E*, and hAzoF_fI73P-fI75K *Z*). The mutation hC84A was reverted manually. Geometries for substrate, cofactor and both isomers of the UAA were optimized performing cluster models at the GFN2-xTB level of theory. Partial atomic charges were derived using the RESP model,[33] fitted to the electrostatic potential (ESP) calculated at the B3LYP/6-31G(d) level of theory. The ESP was computed following the Merz-Singh-Kollman scheme,[34,35] and solvation effects were modeled using the Solvation Model based on Density (SMD) with diethyl ether as the solvent using Gaussian16.[36]

The MD parameters for L-Glutamine, ProFAR, and isomers of hAzo (*E*/*Z*) were generated with the antechamber and parmchk2 modules of AmberTools23 (Amber 22)[37] using the second generation of the general amber force field (GAFF2).[37,38]

The protonation states were predicted using PROPKA.[39,40] The enzyme structures were solvated in a pre-equilibrated truncated octahedral box of 12 Å edge distance using the OPC water model and neutralized by the addition of explicit counterions (i.e., $Na^+$) using the AmberTools23 (Amber22) leap module.[37] All MD simulations were performed using ff19SB forcefield.[41]

**MD simulation details**

MD equilibration phase was done following the protocol described by Roe and Brooks with small differences fine-tuned to our systems.[42] The bonds involving hydrogen are constrained by the SHAKE algorithm during the non-minimization steps. Long-range electrostatic effects were modeled using the particle mesh-Ewald method.[43] For Lennard-Jones and electrostatic interactions, a 10 Å cutoff was applied. The MD protocol starts with the minimization phase of 3000 steps of the steepest descent method followed by 7000 steps of the conjugate gradient method with a positional restrain (i.e., a force constant of

5.0 kcal mol$^{-1}$ Å$^{-2}$) to the protein heavy atoms. In the following heating phase, a temperature increment from 25 to 301.15 K during 20 ps of MD simulation time, a Langevin thermostat with a collision frequency of 5 ps$^{-1}$, and a positional restrain (i.e., a force constant of 5.0 kcal mol$^{-1}$ Å$^{-2}$) to the protein heavy atoms are performed. A minimization and heating of all atoms in the system is the following step. This starts with two minimization stages of 1000 steps of the steepest descent method followed by 1500 steps of the conjugate gradient method each with a positional restrain (i.e., force constant of 2.0 kcal·mol$^{-1}$ Å$^{-2}$ in the first minimization and 0.1 kcal·mol$^{-1}$ Å$^{-2}$ in the second) to the protein heavy atoms. Following, a third minimization phase of 1500 steps of the steepest descent method followed by 3500 steps of the conjugate gradient method without any positional restraint is performed. The system is then heated in accordance with the previously established procedure. Finally, a five-round equilibration phase at the NPT ensemble with a constant pressure of 1 atm is performed. The first four rounds were done with the Berendsen barostat, whereas the fifth one was done with a Monte Carlo barostat. For all equilibration rounds, Langevin thermostat with a collision frequency of 1 ps$^{-1}$ was used. A positional restraint to the protein-heavy atoms with a force constant of 1.0 and 0.5 kcal·mol$^{-1}$ Å$^{-2}$ was applied to the first and second equilibration rounds, respectively. In the third round of 10 ps equilibration, a positional restraint to the backbone-heavy atoms with a force constant of 0.5 kcal·mol$^{-1}$ Å$^{-2}$ was used. The fourth and fifth equilibration of 10 ps and 1 ns, respectively, were performed without any restraint. The production runs were performed at the NVT ensemble with the Langevin thermostat with a collision frequency of 1 ps$^{-1}$ during 150 ns for all systems. A total of three replicas of equilibration and production runs were performed reaching a total simulation time of 0.75 µs/system (5 replicas × 150 ns). The MD trajectories were analyzed using the Python packages MDTraj,[44] pytraj,[45] cpptraj,[46] MDAnalysis,[47] Shortest Path Map,[48,49] and NCIPLOT.[50]

**Conformational landscape (CL) reconstruction**

MD simulations allow the sampling of the population distribution of biomolecules by integrating Newton’s laws of motion. However, due to the vast number of atoms involved in the MD simulations, this probability distribution of molecular states is represented in an extremely high-dimensional space. This is usually solved by focusing on a selected set of degrees of freedom (DOF) relevant to the process of interest. In our case we used: the distance between hH178 and atom NE2 with hC84 and atom name SG (*y* axis) and the distance between hH178 and atom NE2 and L-Glutamine and atom N2 (*x* axis in all Figures). High dimensional data obtained from MD simulations can be projected onto these DOFs to obtain probability distributions. So, a maximum in the distribution corresponds to the most frequently visited, thus the most stable conformation of the protein.
